# Notch signaling blockade links transcriptome heterogeneity in quiescent neural stem cells with their reactivation routes and potential

**DOI:** 10.1101/2024.11.01.621547

**Authors:** David Morizet, Isabelle Foucher, Alessandro Alunni, Laure Bally-Cuif

**Affiliations:** Zebrafish Neurogenetics Unit, Institut Pasteur, UMR3738, CNRS, Team supported by the Ligue Nationale Contre le Cancer, Paris (75015), France; Sorbonne Université, Collège doctoral, F-75005 Paris, France

**Keywords:** Adult neural stem cells, telencephalon, pallium, zebrafish, quiescence, Notch, Nr2f1, single-cell RNAseq

## Abstract

In the vertebrate brain, neural stem cell (NSC) quiescence is necessary for stemness maintenance. Using single cell RNA sequencing (scRNAseq) in the zebrafish adult telencephalon, we identified different molecular clusters of quiescent NSCs, interpreted to sign different quiescence depths(*1*). Here, we show that these clusters, when challenged *in vivo* with an inhibitor of Notch signaling -a major quiescence promoting pathway-, unfold different behaviors. Notably, deeply quiescent NSCs with astrocytic features display a unique activation phenotype that combines the maintenance of astrocytic markers with the rapid upregulation of activation and neuronal commitment genes, reminiscent to murine periventricular astrocytes activating upon lesion. In contrast, an NSC cluster predicted to be in the deepest quiescence state resists Notch blockade, and we demonstrate that the transcription factor Nr2f1b mediates this resistance to activation *in vivo*. These results together link the molecular heterogeneity of quiescent NSCs with *bona fide* biological properties and their molecular regulators.

## Introduction

Somatic stem cells (SCs) reside in many tissues where they maintain homeostasis and can be a source of adaptability. In tissues with little turnover such as muscles and the nervous system, SCs reside in a quiescent state, only dividing infrequently. Interfering with quiescence maintenance leads to increased production of progeny cells but ultimately results in SC exhaustion (*2–4*), suggesting that stemness goes hand in hand with quiescence. At the same time, a certain level of basal SC activity must be maintained to support the needs of the tissue. Understanding how quiescence is modulated to ensure SC maintenance and activity remains a key unresolved issue. Current evidence points to two possibly intertwined mechanisms accounting for dynamic SC maintenance overall: the involvement of several SC subpopulations with distinct properties and in particular different quiescence durations, or the transitioning of individual SC through changing quiescence depths that control their propensity for activation. Live imaging and clonal analyses of adult neural stem cells (NSCs) in the hippocampus identified subpopulations differing in their self-renewal ability and quiescence depth (*5, 6*). Moreover, adult NSCs that have divided once display an increased probability of dividing again, and a decreased probability of being maintained (*3, 7*). Thus, a model was proposed distinguishing dormant NSCs, which have never divided after being established, and resting NSCs, which have divided at least once and returned to quiescence. Finally, latent NSCs have also been described to refer to cells that normally do not proliferate nor generate neuronal progeny but can be induced to do so in a regenerative context (*8*). This diversity of function and/or potential among quiescent NSCs (qNSCs) is an evolutionarily conserved feature. Indeed, in the zebrafish adult pallium, clonal analyses and intravital imaging also suggest that two subpopulations of qNSCs with different outputs are present, displaying different quiescence durations and hierarchically organized (*9, 10*). Moreover, quiescence depth appears to be a dynamic property. For example, the expression of some markers by qNSCs is correlated with time spent in quiescence since their last division (*11*), and qNSCs re-enter the cell cycle with different delays upon blockade of Notch signaling, a major quiescence-promoting pathway (*12*). Thus, quiescence heterogeneity among qNSCs at any given time can reflect distinct sub-lineages or trajectory positions. To date, the molecular and functional correlates of these differences are poorly characterized.

Recently, single cell RNA sequencing (scRNAseq) has been used to characterize neurogenic niches in mouse (*5, 7, 13–19*). These studies have successfully identified transcriptional differences linked to regionalization and apparent continuums from deep quiescence to activation. However, so far, it has proven challenging to exploit these scRNAseq data to identify subpopulations of qNSCs intermingled in the same progenitor domain and associated with distinct quiescence depths. Moreover, the slow kinetics of qNSCs, which can remain quiescent for weeks, do not lend themselves to methods for trajectory reconstruction such as RNAvelocity, which perform best at inferring dynamics on the scale of hours (*20*). Using scRNAseq, we also recently described molecularly distinct qNSC clusters predicted to reside in different quiescence states in the zebrafish dorsal pallium (*1*). Our reanalysis of other published datasets including zebrafish telencephalic NSCs (*21–24*), with validations of gene expression *in situ* and comparisons across vertebrate species, supported our partition of qNSCs. Here, we aimed to establish the significance of this scRNAseq-based categorization in terms of qNSC properties, by directly probing the putative distinct quiescence depths taken by qNSCs -as defined by their propensity to reactivate upon release of quiescence-promoting cues- and their pathways to activation.

To do so, we experimentally nudged cells towards a more activated state and exploited the comparison between physiological and “activation-oriented” scRNAseq data. The Notch pathway plays a role in the establishment (*25*), maintenance (*26–30*) and molecular control of adult neurogenesis (*12, 31, 32*). Upon inhibition of Notch signaling, NSCs in the zebrafish telencephalon (*12*) and the murine hippocampus (*33*) or subependymal zone (SEZ) (*32*) quickly enter the cell cycle to proliferate. In zebrafish, qNSC reactivation following treatment with LY411575 (LY), a validated gamma-secretase inhibitor, is asynchronous and dependent on the length of treatment (*12*). Therefore, we reasoned that profiling qNSCs following a short LY treatment regimen, under which most NSCs do not yet enter the proliferating state, could circumvent the slow kinetics of qNSCs and reveal different reactivation speeds and/or trajectories among them. We identified conditions for subthreshold Notch inhibition *in vivo* and used these to perform scRNAseq. Overall, combining LY-treated and untreated NSCs first refines the classification of adult NSCs in the zebrafish pallium. It also reveals heterogeneous responses to the forced transition towards reactivation triggered by LY treatment and maps them to specific molecular identities, demonstrating the functional diversity of qNSCs and highlighting previously unrecognized activation trajectories in vertebrate NSCs. Finally, we discover a rare subpopulation of qNSCs which shows low sensitivity to Notch pathway inhibition, and we identify the transcription factor Nr2f1b as a positive regulator of this state characterized by deeper and more robust quiescence. Together, these results confirm that qNSC transcriptomic heterogeneities correlate with specific NSC quiescence/activation behaviors and identify one molecular regulator of a deeply quiescent substate.

## Results

### Transcriptionally-defined subpopulations of quiescent NSCs remain identifiable after short Notch pathway inhibition

To study the dynamics of transitions between different quiescent scRNAseq clusters, we searched for an experimental scheme to bypass the normally slow kinetics of these transitions while simultaneously avoiding that NSCs enter proliferation proper. We previously showed that inhibition of canonical Notch signaling with LY411575 (LY) is a potent pro-activation cue for zebrafish pallial NSCs (*31, 12*). After 48 hours of treatment, 60% of NSCs are positive for the proliferation marker protein Pcna, compared to 5-10% under normal conditions. We reasoned that, before this happens, there should be a time window when LY treatment already acts, displacing qNSCs towards activation without reaching proliferation. To test this hypothesis, we bathed adult fish (3-month post-fertilization -mpf-) in LY (or DMSO as control) for different durations (Fig. 1A). As a readout of a shift towards activation, we assessed the effect of LY treatment by quantifying the percentage of Pcna-positive (Pcna^pos^) NSCs and using the expression of *ascl1a*, which is expected to be repressed by Notch pathway effector genes when Notch signaling is active (*34*) (Fig. 1B-F, Fig. S1A-D). In mouse, high expression of the transcription factor Ascl1 also characterizes a pre-activated state and its upregulation precedes, and is necessary for, cell cycle entry (*35*). This analysis revealed that LY treatment efficiently induced *ascl1a* expression as early as 12 hours after initiation of the treatment and very obviously by 24 hours (Fig. 1D-E’) while at that time most cells had not yet entered proliferation (Fig. 1F). This was validated at both rostral and caudal levels (Fig. 1F and not shown). Thus, a short Notch pathway inhibition of 24 hours meets our two criteria.

**Figure 1.**
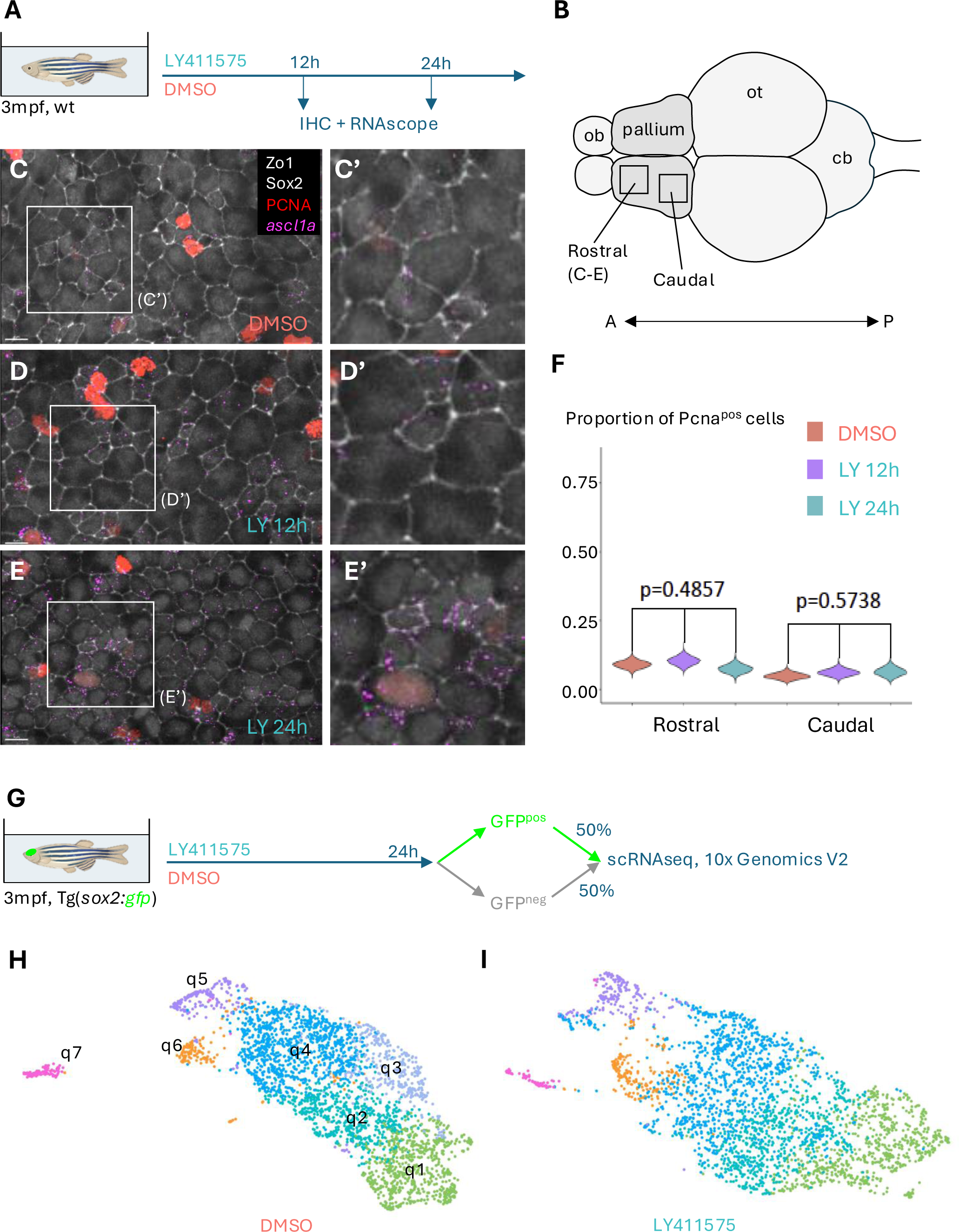
A brief inhibition of Notch signaling activates quiescent NSCs without depleting them in favor of cycling NSCs. **A,B**. Experimental scheme (A) of the treatment of 3mpf wildtype adult zebrafish with DMSO or LY411575 for 12h or 24h, followed by whole-mount immunohistochemistry (IHC) and *in situ* hybridization (RNAscope) and cell quantifications. A rostral and a caudal area of the pallial ventricular zone were analyzed, as depicted on the zebrafish brain cartoon (B) (dorsal view, A: anterior, P: posterior). **C-E.** Representative images of the pallial ventricular zone (rostral areas) after treatment either with DMSO (24h) (C) or LY for 12h (D) or 24h (E). C’-E’ are higher magnifications of the areas boxed in C-E. Whole-mount dorsal (apical) views, anterior left. Zo1 and Sox2 immunostaining (white) are used to identify progenitor cells with apical ventricular contact, Pcna (red) is used to label and count proliferating cells (quantifications shown in F) and *ascl1a* RNAScope (magenta) is used as a readout of decreased Notch activity and transition towards the pre-activated state. *ascl1a* is expressed at higher levels after LY treatment and higher after 24 hours than after 12 hours of treatment. See also Fig.S1D. Scale bars: 8µm**. F.** Quantification of the proportion of cycling cells (Pcna^pos^) after each treatment in the rostral or in the caudal part of the dorsal pallium. Indicated p-values are the result of a Kruskal-wallis test. Violin plots are built from bootstrapped random sampling of the measured proportions to estimate the distribution. **G.** Experimental scheme used to generate the scRNAseq dataset under conditions of LY treatment. The DMSO control dataset was reported in (*1*). **H.** Low-dimensional embedding of quiescent NSCs (qNSCs) in the control scRNAseq dataset colored by cluster (after (*1*)). **I.** Low-dimensional embedding of qNSCs in the scRNAseq dataset of LY-treated fish, colored by cluster and matching colors to those from the control dataset based on inferred equivalence.

Next, we used fish from the Tg(*sox2*:GFP) line (*36*), where GFP expression identifies NSCs and intermediate progenitors (IPCs) (*37*). We treated 3mpf adults with LY for 24 hours, immediately dissected the telencephalon, sorted GFP^pos^ cells and mixed them with an equal quantity of GFP^neg^ cells to enrich in NSCs without overlooking potentially relevant *sox2*^neg^ cells (Fig. 1G). We then performed droplet-based single cell RNAseq, as conducted for the physiological dataset (*1*). We identified the different cell types using well established markers (Table S1) and will focus below exclusively on NSCs and IPCs. Proliferating cells formed a distinct cluster which allowed us to isolate non-proliferating NSCs to subcluster them. We built a consensus matrix from the results of several clustering algorithms (*38–43*). With this approach, the frequency at which cells are grouped together with different methods serves as a distance measure to compute a hierarchical tree from which final clusters can be obtained. In our previous study, we chose a conservative cutoff, which yielded 7 clusters in the DMSO dataset (Fig. 1H) (*1*). Two of these clusters (q6-q7) were made up of spatially segregated NSCs in the subpallium and at the boundary between pallium and subpallium. The other five (q5-q1) were made up of qNSCs from the pallium and were predicted, based on their molecular profile and comparison with prospectively isolated NSCs, to reside in a progressively deeper state of quiescence from q1 to q5. The same approach yielded 6 clusters in the LY dataset (Fig. 1I), failing to identify cells from q3 as an independent cluster likely due to their transcriptional proximity to q2 and q4 and to the effect of LY treatment that might have made it more difficult to distinguish between them. Next, we assessed the expression of combinations of genes which can identify the different NSC subpopulations in the zebrafish telencephalon (*1*). We found that the patterns of expression of these genes are very similar between DMSO and LY conditions and that equivalent clusters could readily be identified (Fig. S1E,F). Thus, we conclude that core cell identity is not substantially affected by short Notch inhibition and that direct comparisons between similar cells can be performed to assess the effects of LY treatment.

### Gamma-secretase inhibition helps uncover rare molecular subpopulations of qNSCs and reveals the multi-faceted roles of Notch signaling in their regulation

The results above show that a 24-hour LY treatment enables the study of transcriptomic trajectories of the different clusters of qNSCs under Notch blockade and their dependency on Notch signaling. We reasoned that, in addition, integrating scRNAseq datasets acquired with or without LY treatment could reveal functional differences among qNSCs and molecular subpopulations perhaps too rare to be observed under normal circumstances (*44*). Indeed, although inspection of the two datasets separately is valuable, jointly defining the cells from the control and treated datasets is needed to confirm cluster differences and to identify transcriptional changes linked with LY treatment in given clusters. In order to do so we turned to the Liger algorithm, developed to integrate scRNAseq datasets (*45, 46*). We previously showed that the zebrafish telencephalon contains regionally restricted or regionally enriched qNSC subpopulations. These include the *nkx2.1*^pos^ population from the ventral subpallium (Fig. 1H, q7), the *gsx2*^pos^ population from the dorsal subpallium (q6), and a population of qNSCs which is enriched caudally and expresses high levels of *pnp6* (q5) (*1*). These subpopulations could be detected in both untreated and treated datasets individually (Fig. S1E,F, dotted circles to q6 and q7 and arrow to q5) which allowed us to use them as an internal cellular control to fine-tune the parameters of the integration. This resulted in a joint embedding with both good mixing of the datasets and preservation of the intrinsic variability between cells in the data (Fig. S2A-C).

As expected given clustering sensitivity to cell numbers, joint clustering yielded more clusters than independent clustering on either dataset (Fig. 2A). We projected the cells belonging to all the jointly defined clusters back to the individual datasets to determine whether they were made up of cells that were originally close together and/or that would have been identified as belonging to an independent cluster with slightly less stringent cutoffs for the consensus clustering (see Methods). Ultimately, this led to the identification of rare subpopulations of cells (Fig. 2A). Specifically, clusters of cells predicted to be closer to activation (identified relative to the *ascl1a*^pos^ cells in the control dataset) could be subdivided into six clusters (q1a to q1f). When comparing untreated cells belonging to cluster q1a to untreated cells belonging to clusters q1b, q1e and q1f, we found that the latter expressed higher levels of proneural genes (*neurod*, *sox4/11*, *eomesa*), as well as genes associated with cell division or commitment (*ccnd1*, *ascl1a*) (Fig. S2D). Some of these q1 clusters (q1c in particular) also expressed genes likely related to the generation of specific neuronal types such as *zic2a* and *zic3* (*47*) (Fig. S2D). Also, we previously identified astrocyte-like NSCs, presumably deeply quiescent although constitutively neurogenic under physiological conditions (cluster q4, Fig 1H) (*1*). These are identified by high expression of genes such as *timp4.3* and *igfbp2a* (Fig. S1E). In the integrated analysis, we found that these were subdivided into two clusters q4a and q4b, the latter having a transcriptome with many features at intermediate levels between q4a and q5 (Fig. 2A, S2D). Finally, we also identified a new cluster, q8, containing only a few NSCs (≈2.4% of qNSCs), conserved in all datasets. Despite containing cells with a low number of UMIs it was still enriched for several genes (Fig. S2D) suggesting that the low number of UMIs did not come from a technical issue. Some quiescent cells can have low metabolic activity or rely less on polyadenylation of mRNA (*48*), which would lead to low UMIs with the scRNAseq method we used. Together, the integrated dataset helped us refine our initial clustering to resolve a higher level of transcriptomic NSC heterogeneity.

**Figure 2.**
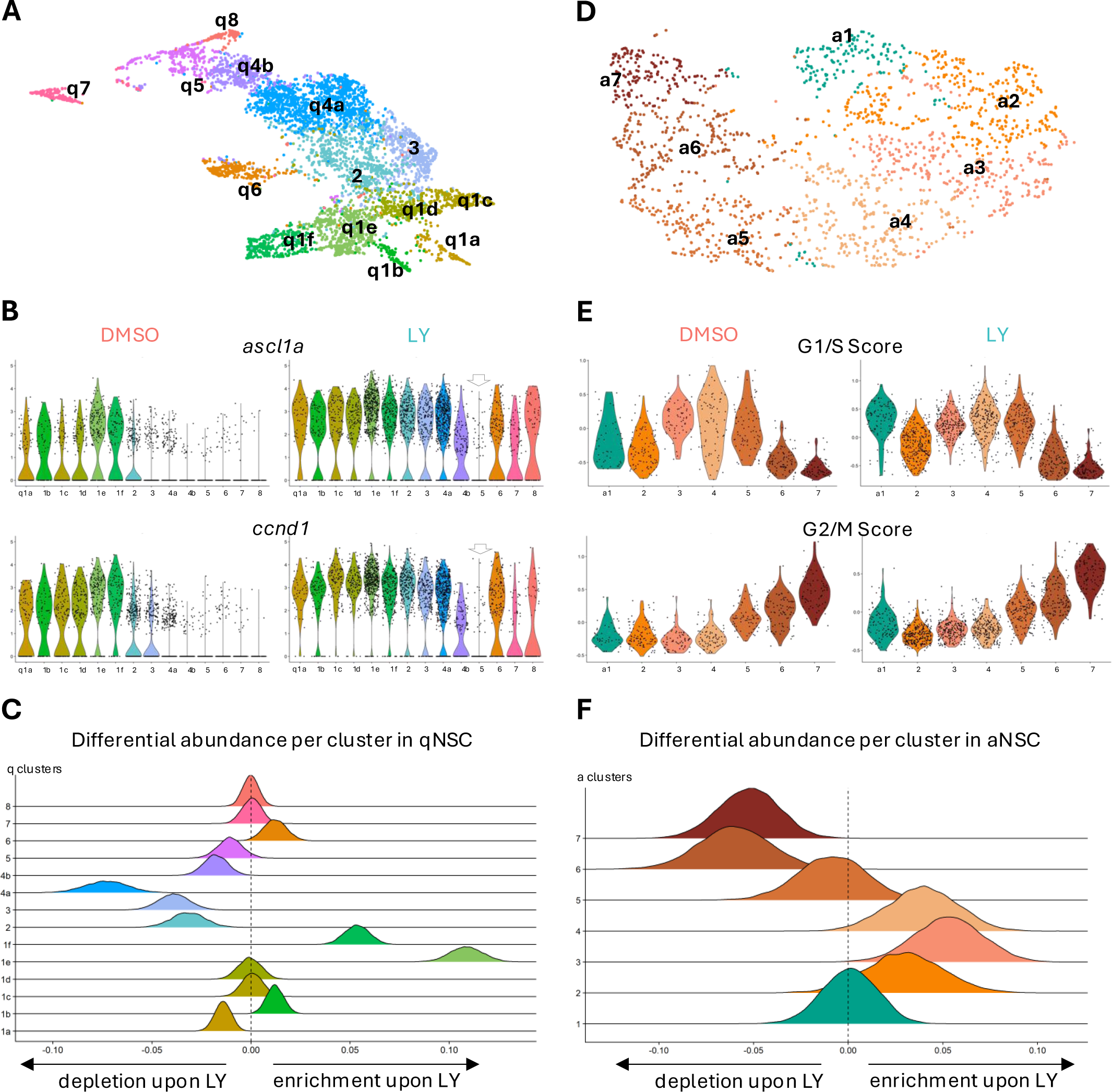
Data integration highlights Notch inhibition-induced changes in NSCs at the cellular and molecular levels. **A.** Low-dimensional embedding of qNSCs after integration of LY-treated and control datasets (Fig. 1H,I, Fig. S2A). Cells are colored based on the refined cluster annotations derived from integrated analysis; cluster q1 resolves into 6 clusters (1a-1f), cluster q4 resolves into 2 clusters (4a, 4b), and a previously overlooked cluster 8 is identified. **B.** Violin plot comparing the expression of *ascl1a* and *ccnd1*, genes associated with cells moving towards an activated state, across all clusters for control and treated cells. Clusters are colored as in Fig.2A and ordered from 1a to 8 from left to right. Y axis: number of reads normalized for sequencing depth. **C.** Depiction of the over or under-representation of distinct subpopulations of qNSCs between control and treated datasets. The clusters are color coded as in Fig. 2A. A curve displaced to the right represents an enrichment following LY treatment. The further a curve is shifted away from the central line, the more a given cluster is enriched (to the right) or depleted (to the left) in the treated dataset. **D.** Low-dimensional embedding of cycling cells after integration of LY-treated and control datasets. Cells are colored based on the refined cluster annotations derived from integrated analysis. Cluster 1 in green represents cycling NSCs and clusters 2 to 6 represent IPCs progressively more advanced in the cell cycle. **E.** Violin plot comparing the G1/S and G2/M scores across all clusters of cycling cells for control and LY-treated cells. Clusters are colored as in Fig. 2D. Absolute values do not reflect direct expression of genes and comparison must rely on relative values of each score. **F.** Depiction of the over or under-representation of distinct subpopulations of cycling cells between control and LY-treated datasets. The clusters are color coded as in Fig. 2D. A curve displaced to the right represents an enrichment following LY treatment. The further a curve is shifted away from the central line, the more a given cluster is enriched (to the right) or depleted (to the left) in the treated dataset.

Using this integrated dataset, we next probed for differences between the control and LY datasets in each commonly defined cluster. In the LY dataset, almost all NSCs expressed *ascl1a* whereas it was restricted to a minority of NSCs in the control dataset (Fig. 2B). Upregulation of *ascl1a* in these conditions is expected, as it is normally repressed by Her factors, orthologous to mammalian HES, which themselves are the effector genes of the canonical Notch pathway and indeed appeared to be downregulated in most clusters (Fig. S2F) upon LY treatment. This was accompanied by the upregulation of genes associated with entry into the cell cycle such as *ccnd1* (Fig. 2B), a target of Ascl1 in mice (*35*) which is necessary for progression through G1, confirming that NSCs had started transitioning towards activation. We then looked for differentially expressed genes between untreated and LY-treated cells in each cluster (Table S2). As expected in the LY-treated dataset we found broad downregulation of Notch target genes (Fig. S2F) and, in particular, of the *her4* and *her15* paralogs (orthologous to mouse *Hes5*) as well as *her9* and *her6* (orthologous to *Hes1*). This was accompanied by a broad upregulation of genes involved in initiating cell cycle, such as *pcna*, *mcm* genes or *ccnd1*, even though qNSCs remained distinct from the proliferative clusters. In several clusters (notably q2, q3, q4a, q4b and to varying degrees in q1 clusters), we also found downregulation of glial genes (Fig. S2F and Table S2) such as *slc1a2b*, *glula* and *metrn*, a glial cell differentiation regulator which promotes stemness in astroglia (*49*). Conversely, genes associated with neuronal differentiation such as *sox4a*, *sox11a, stmn1a* and *gadd45gb.1* were upregulated. Many of the changes that we observed when comparing treated and untreated cells in a given cluster mirror differences in expression between cells close to activation and more quiescent cells under normal conditions. This suggests that those differences recapitulate molecular events associated with the transition towards states of increased activation frequency in physiological conditions.

Finally, we asked whether the two datasets differed also in terms of cluster abundance, i.e., proportion of cells per cluster in each dataset. We designed a differential abundance test based on a Monte Carlo simulation aiming to identify whether some clusters represented a larger proportion of cells in a dataset than in another. This revealed a statistically significant enrichment in clusters q1b, q1e and q1f and a significant depletion in clusters q1a, q2, q3, q4a and q4b in LY-treated over control cells (Fig. 2C). Overall, activation-associated states are enriched at the expense of quiescence-associated states.

Together, Notch pathway inhibition in qNSCs promotes their expression of activation markers and triggers the expression of proneural genes at the expense of genes maintaining glial identity and stemness. Likely depending on the extent of these changes, this is accompanied, or not, with switches in cluster identity for individual cells.

### Notch signaling contributes to gating cell numbers and preventing neuronal commitment in proliferating populations

The Notch pathway is also active in proliferating cells where it controls the balance between differentiation and self-renewal (*12, 26, 27, 31*). Moreover, although we did not see increased PCNA immunostaining after 24 hours of LY treatment (Fig. 1C-F), transcriptionally defined cycling cells represented a larger proportion among stem and progenitor cells in the LY dataset (percentage of cycling cells among all NSCs and IPs: 31.02% upon LY treatment vs 14.36% under DMSO) and this increase affected both NSCs and IPCs (Fig. S3A,B). We thus extended our analyses to cycling cells.

First, we subclustered cycling cells from the untreated dataset. Despite all proliferating cells often being grouped together and called IPCs in scRNAseq datasets (*13, 14, 17*), we could distinguish between cycling NSCs and cycling IPCs (Fig. 2D). NSCs could be separated from IPCs due to higher NSC expression of genes such as *mfge8*, *fabp7*, *slc1a2* and *sparc* and lower expression of genes such as *insm1*, *elavl3*, *sox4* and *sox11* (Fig. S3C,D). We also re-analyzed datasets generated from the mouse SEZ which included a large population of cycling cells (*13, 17*) to identify conserved patterns of expression and highlight genes with a putatively important regulatory role. There too, we could distinguish between NSCs and IPCs (Fig. S3C). Cycling NSCs and IPCs also differed in the expression of several putative regulators of NSC behavior. In particular, *Hes5* was enriched in NSCs over IPCs in mouse and so was its ortholog *her4.1* in zebrafish (Fig. S3D), highlighting it as a potential mediator of Notch pathway-mediated promotion of self-renewal. Next, we curated a list of genes with periodic expression during the cell cycle using Cyclebase (*50*) to generate refined cell cycle scores (Material and Methods). This revealed IPCs in different phases of the cycle in both zebrafish and mouse (Fig. S3C). NSCs were too few for the latter distinction. Differences in gene expression between cells in late or early cell cycle belonging to the same cell type were modest in genes not directly involved in cycling or cycling-associated chromatin remodeling such *Dnmt1*, *Dnmt3* and *Hdac1*.

Second, we integrated the treated and untreated zebrafish datasets of cycling cells with Liger (Fig. 2D,E, Fig. S3E). Cycling NSCs formed a single cluster (cluster a1) while IPCs formed multiple clusters reflecting different levels of progression into the cell cycle (from cluster a2 in early cell cycle to cluster a7 in M phase). Differentially expressed genes (DEGs) overall were similar, albeit less numerous, to the ones identified for quiescent cells, and the number of DEGs was higher for NSCs than for IPCs (Table S3). In particular, the expression of several genes associated with radial glia, such as *fabp7a*, *glula* and *her4.1* was significantly and substantially decreased upon LY treatment, while expression of neuronal commitment genes such as *stmn1a* (in NSCs) and *sox4a* and *sox11a* (in IPCs early in the cell cycle) was increased (Table S3). Genes involved in early cell cycle phases, such as *ccnd1* and *mcm2*, were also increased upon LY treatment in cycling NSCs (Table S3). Finally, we tested for differential abundance which revealed that IPCs in early phases of the cell cycle are over-represented in LY-treated versus control samples, whereas IPCs in M phase are under-represented and the proportion of cycling NSCs (among cycling NSCs + IPCs) is not significantly changed (Fig. 2F). The proportional increase of a transcriptomic signature for early cell cycle phases in IPCs is most likely because they are freshly recruited and have not yet had the time to progress to the later phases, consistent with our inability to detect significant changes in Pcna^pos^ cells with immunostainings. It is unlikely that their progression becomes halted, as longer LY treatments that trigger proliferation in almost all stem and progenitor cells (*12*).

Together, these results reveal that Notch blockade for 24 hours in the zebrafish adult pallium does trigger cell cycle entry in a subset of NSCs and IPCs, detectable by scRNAseq, with a proportional enrichment for G1/S markers among IPCs. Notch blockade also exerts an effect on cycling cells with a tendency to increase expression of neuronal commitment genes.

Overall, our analysis uncovers differential responses of distinct subsets of NSCs to an activating stimulus, highlights multi-faceted roles of Notch signaling in promoting quiescence and stemness, inhibiting neuronal differentiation and maintaining glial identity. Among the deregulated genes following Notch pathway inhibition, some, such as *metrn* and *Hes5*/*her4.1,* may be downstream mediators of these different roles in NSCs.

### Astrocyte-like NSCs can reactivate in a manner reminiscent of murine latent NSCs

Our differential abundance analysis in qNSCs showed that the cell subpopulations most affected, in their relative amount, by LY treatments, belong to clusters q1e, q1f and q4a (Fig. 2C). We recently showed that cluster q4 (q4a + q4b) possesses astrocytic features and is closer to murine astrocytes than to murine radial glia-like (RG-like) cells (*1*). In zebrafish, these cells retain a neurogenic potential under physiological conditions, but pseudo-ordering, GSEA analysis and the overall mutual exclusivity of their highly expressed marker genes and activation markers suggest that they are deeply quiescent. We thus wanted to determine how these cells came to be depleted by LY treatment-mediated activation.

Under physiological conditions, q4 qNSCs can be identified by a set of genes associated with astrocytic functions. *igfbp2a* and *timp4.3* in particular are highly enriched in q4 cells compared to other qNSCs (*1*) (Fig. S1E). We looked for expression of these genes in the integrated dataset separately in treated and untreated cells, to determine whether q4 depletion results from a decrease in the number of cells expressing q4 markers or whether these cells are still present but cluster separately after treatment. In the LY dataset, cells expressing high levels of q4 markers in fact represented a large fraction of q1e and q1f NSCs suggesting that the second interpretation was correct and that these cells had been pushed to the q1e and q1f sub-states while maintaining q4 markers (Fig. 3A).

**Figure 3.**
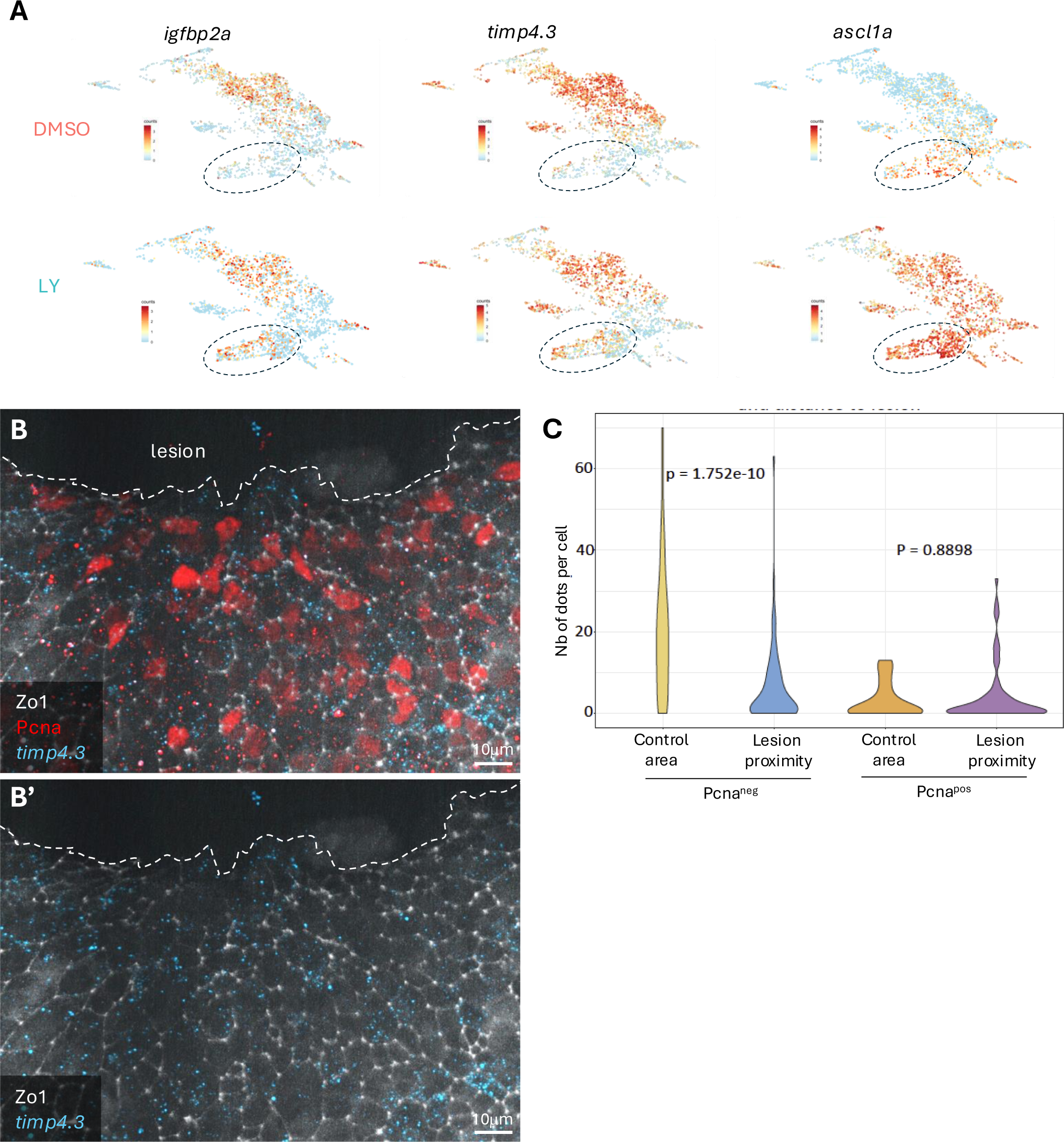
Zebrafish astrocyte-like cells respond to Notch inhibition and injury. **A.** Levels of expression of indicated genes in qNSCs from DMSO or LY-treated datasets projected on the integrated embedding of qNSCs. *igfbp2a* and *timp4.3* are markers of astrocyte-like cells and *ascl1a* is a marker of pre-activated cells and a readout of lack of Notch pathway activity. Clusters q1e and q1f are circled. In the LY dataset but not in the control, many cells co-express astrocyte-like NSC markers and *ascl1a*, in particular in clusters q1e and q1f. **B.** High magnification of the ventricular surface of a pallium close to a lesion (dorsal whole-mount view), processed for immunohistochemistry (ZO1 -white-, Pcna -red-) and smFISH (*timp4.3* - cyan-). Zo1 is used to delineate apical areas and identify cell contours. Proliferating cells are labelled with Pcna and appear with a red nucleus in greater number in cell rows neighboring the lesion. *timp4.3* expression was revealed using RNAScope; each dot represents one molecule of RNA. The top view shows all channels, the bottom views shows Zo1 and *timp4.3* only. Scale bar: 10µm. **C.** Quantification of *timp4.3* expression (number of dots per cell) Pcna^neg^ and Pcna^pos^ cells neighboring the lesion (“Lesion proximity”) or far away from the lesion (“Control area”). Represented p-values were calculated with a Wilcoxon signed rank test to avoid inflating sensitivity with a bootstrapped test. Control area: n=3 brains, 92 cells; Lesion proximity: n=3 brains, 149 cells. Overall *timp4.3* levels close to the lesion are decreased, with a significant decrease in Pcna^neg^ cells. In contrast, Pcna^pos^ cells tend to express *timp4.3* at higher levels close to the lesion than in control cells, but this difference is not statistically significant.

Next, we looked at the genes differentially expressed between treated and untreated NSCs of q1e and q1f. Despite expressing higher levels of glial markers in the LY-treated dataset (see Fig. 3A, Fig. S2D-F, Table S2), these qNSCs also express higher levels of proneural genes and genes associated with entry into the cell cycle (Fig. S2D-F, Table S2), like other clusters. However, although treated cells in q1e and q1f express higher levels of ribosomal genes than cells in q4a, we found that in contrast to q1a-d, LY-treated NSCs in q1e and q1f expressed lower levels of several ribosomal genes (Fig. S4A) than untreated NSCs from the same integrated clusters. This depletion being specific to q1e and q1f suggests that it is not a batch effect. Upregulation of ribosomal genes is a step that precedes entry into the cell-cycle (*51*) and appears to be conserved across other contexts (*52, 53*). Ribosome biogenesis is coupled to cell growth (*54*), in turn tightly linked to the ability to proliferate (*55, 56*). The limited ribosomal genes response of q1e and q1f, together with their maintained expression of several astroglial markers which are physiologically not co-expressed with activation markers, suggest that q4a NSCs pushed to clusters 1e and 1f by Notch inhibition undergo a sort of “rushed activation”.

We then sought to compare our data with data derived from studies conducted in mice, where astroglial cells that show peculiar patterns of activation were also described. Striatal astrocytes have been reported to possess a latent neurogenic potential: while they do not produce neurons under physiological conditions, they can be induced to do so by stimuli such as a stroke (*8, 16, 57*). This astrocytic activation requires Notch signaling to be attenuated and can be mimicked by genetic inhibition of the Notch pathway (*8, 58*). In addition, it has also been suggested that cells described as deeply quiescent RGL in the SEZ (referred to as qNSC1) can be forcefully reactivated upon injury (*16*). These cells have been described as corresponding to B1 cells in the SEZ with an RGL transcriptome but a methylome reminiscent of astrocytes (*57*). Although several datasets profiling the mouse SEZ have now been generated (*13, 15–17, 19, 57, 59–61*), most of them did not identify a cluster matching qNSC1. We reanalyzed these datasets (see also Supplementary text) and found that although such cells were present they had been classified either as astrocytes (*13*) or RGL (*15, 16, 57, 60*), with other qNSCs being classified either correctly as qNSCs (*13, 15, 16, 57*) or mistakenly as aNSCs (*60*) (Fig. S4B). We then used our previously identified astrocytic gene set (*1*) and Metaneighbor (*62*), and mapped SEZ astroglia to a telencephalon atlas(*63*) comprising both RGLs and parenchymal astrocytes. This suggested that the transcriptome of cells classified as qNSC1 is closer to that of *bona fide* local astrocytes rather than to RGLs (Fig. S4C,D). In accordance with this, a report in which these cells are classified as astrocytes while other cells are classified as RGLs showed that the latter comprise both B1 and B2 cells in the SEZ(*64*). Overall these results suggest that so-called qNSC1 are in fact parenchymal astrocytes close to the SEZ that can be distinguished from RGLs by both their methylome and their transcriptome. We thus treated murine latent NSCs as a single population of cells representing a subset of striatal astrocytes. The induced activation and maturation (*65–67*) of latent NSCs has recently been analyzed. As published (*58*), we confirmed that by 4 weeks of Rbpj deletion (mediating Notch inhibition), activated striatal astrocytes had downregulated many of the genes found in control astrocytes, and upregulated several genes found in activated NSCs. In contrast, at earlier timepoints, striatal astrocytes co-express astrocytic markers with genes associated with activation such as *Ascl1* and *Ccnd1*. We also probed for changes in gene expression in striatal astrocytes 2 days after experimentally induced stroke (*16, 57*). Under these conditions, the astrocyte transcriptome became closer to that of RGLs, while maintaining expression of some astrocyte-specific markers as previously reported (*16, 57*). The low number of cells profiled made it difficult to obtain statistically meaningful results for individual genes. However, we found that although ribosomal genes expression was upregulated in astrocytes that had been reactivated compared to resting astrocytes, many ribosomal genes were expressed at lower levels than in control RGLs at the same stage of progression along the activation trajectory. (Fig. S4A).

Given the similarities between astrocyte-like q4 and murine latent NSCs in their response to Notch pathway inhibition and the fact that Notch pathway inhibition triggers a similar response in latent NSCs as a brain lesion does, we next asked whether q4 cells also respond to injury in the zebrafish telencephalon. To do so we used a stab-wound model where a mechanical lesion was inflicted to the zebrafish pallium through the top of the skull. We combined *timp4.3* single molecule FISH (smFISH) quantification with immunostaining after lesion to assess the behavior of *timp4.3* high-expressing cells, mostly belonging to q4, after lesion (Fig. 3B). The maximum level of proliferation in such conditions is expected to be reached at 5 days post-lesion (5dpl) (*68, 69*) but we could reliably confirm that our lesion had been successful and indeed induced a response among NSPCs by monitoring the expression of cell cycle markers as early as 3dpl (Fig. S4E-G). At 3 dpl, 48% of the cells in the vicinity of the lesion (within the next 5 ± 2 cell rows) expressed Pcna, compared to 15% in cells away from the lesion used as control cells (Fig. S4G). Among Pcna^neg^ cells, the level of expression of *timp4.3* was significantly decreased in the vicinity of the lesion compared to control cells, suggesting that they had been recruited to deal with the lesion (Fig. 3C). The overall distribution of *timp4.3* expression also appeared increased in Pcna^pos^ cells near the lesion compared to control Pcna^pos^ cells, however this did not reach statistical significance (Fig. 3C, S4H). Together, the results from the lesions analyzed with the marker *timp4.3* suggest that q4 cells participate in regeneration, likely by going through a state where they co-express astrocytic markers and activation markers initially before downregulating astrocytic markers at later stages.

In sum, these analyses uncover the existence of a cryptic activation trajectory for astrocyte-like qNSCs in the zebrafish adult pallium. This trajectory is revealed by Notch inhibition and is not -or very rarely-used under physiological conditions. This phenomenon shares common features with medial striatal astrocytes close to the murine SEZ that act as latent NSCs: a persistent expression of astrocytic markers and limited upregulation of ribosome biogenesis in the early phases of activation, in cells that share astrocytic features but are efficiently recruited to regenerate the brain upon injury.

### A gamma-secretase-independent mechanism maintains Notch effector gene expression and quiescence in a subpopulation of NSCs

The expression of *ascl1a* and *ccnd1* was substantially increased in most NSCs after a 24-hour LY treatment, with the striking exception of one NSC cluster: q5 (Fig. 2B arrows, Fig. S1D, S2F). Quiescence in adult somatic stem cells is maintained by several signaling pathways, yet alteration of one of these pathways is often enough to trigger quiescence exit (*4, 70–77, 77, 78*). However, because a general and long-lasting release from quiescence leads to depletion of the NSC pool (*51*), being able to modulate the response of specific NSC subsets to the alteration of quiescence-promoting pathways holds the potential to spare, in part, the NSC pool. For these reasons, we sought to determine whether q5 highlights an NSC cluster resistant to Notch pathway inhibition and, if so, to shed light on how this resistance is mediated.

First, we assessed, in LY-treated versus control datasets, the expression of a large panel of genes dependent on Notch pathway activity as well as genes which are upregulated upon Notch pathway inhibition. This confirmed that q5 cells can maintain the expression of several Notch target genes (such as *her4.1*, *her8a*, *her9*, *hey1*, *fabp7a*) and repress the expression of several pro-activation genes (such as *ascl1a*, *ccnd1*, *sox4a*, *sox11b*, *stmn1a*) even after 24 hours of LY treatment (Fig. 2B, Fig. 4A, Fig. S2E).

**Figure 4.**
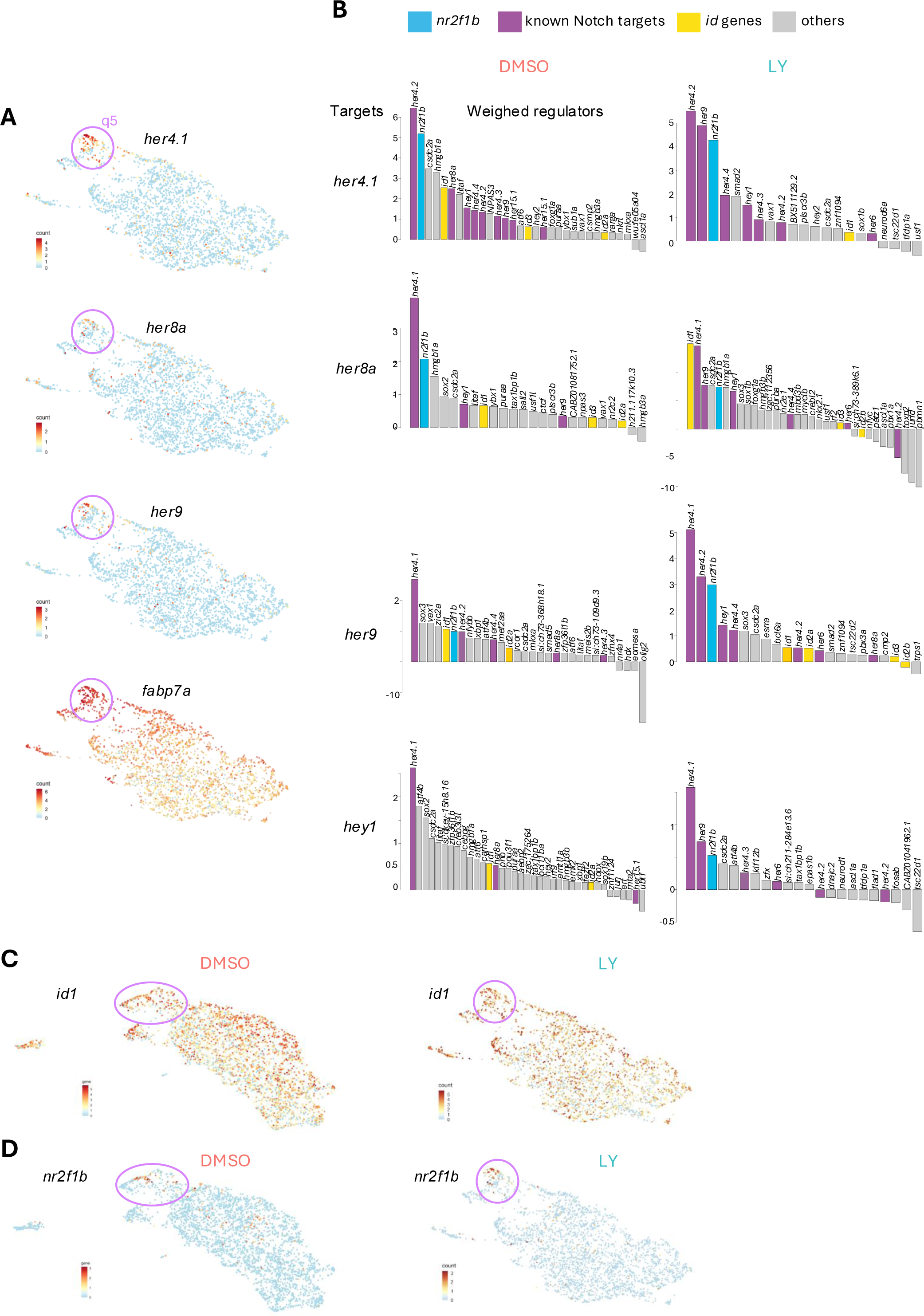
A machine learning approach predicts putative regulators of resistance to Notch signaling blockade in deeply quiescent pallial NSCs of cluster q5. **A.** Levels of expression of major Notch targets represented in the low-dimensional embedding of qNSCs from the LY-treated dataset. Cells belonging to cluster q5 are circled and show higher and almost exclusive expression of these genes. **B.** Barplots depicting the weight associated with putative regulator-target relationships inferred through gradient boosting and automatic filtering. The height of the bar is proportional to the contribution of the indicated transcription factor’s expression to the recovery of the pattern of expression of the target (*her4.1*, *her8a*, *her9*, *hey1*). When transcription factors were anti-correlated with their putative target we assigned a negative sign to the link. Bars are color-coded to highlight specific genes and gene families. **C,D.** Levels of expression of the genes encoding the best candidates regulators of q5 resistance to activation, *id1* (C) and *nr2f1b* (D), represented in the low-dimensional embedding of qNSCs from the control (left panels) and LY-treated (right panels) datasets.

To search for a putative mediator of this resistance to Notch pathway inhibition, we looked for factors that could maintain Notch effector genes’ expression. We inferred putative regulator-target relationships in both LY-treated and untreated conditions by using a gradient boosting approach implemented by GRNBoost2 (*79*). This method calculates a weight representing how much information on the expression of a given gene can be recovered by knowing the expression of a given transcription factor. We chose an adaptive threshold based on an elbow rule to refine lists of putative regulators (see Methods). We looked for transcription factors appearing as likely regulators of *her4.1*, *her9*, *her8a* and *hey1. her4.1* is the ortholog of *Hes5* which shows the highest correlation with inferred quiescence depth, *her9* is the ortholog of *Hes1* with the highest expression in qNSCs, *her8a* is phylogenetically closer to *Hes6* but was found to play a role in inhibiting neurogenesis in zebrafish (*80*) and *hey1* is another Notch effector gene enriched in qNSCs and associated with stemness maintenance (*25, 30*). We found that only few transcription factors were consistently predicted as putative regulators for these genes in both LY-treated and untreated conditions. Specifically, Id1 and Nr2f1b/Coup-tf1b were the two main transcription factors highlighted by this analysis when accounting for the likely co-regulation of multiple Notch effectors (Fig. 4B). *id1* is expressed broadly in NSCs, reaching its highest levels in q5, while *nr2f1b* expression is highly restricted to q5 (Fig. 4C,D). *id1* encodes an HLH protein which can heterodimerize with bHLH transcription factors and decrease their affinity for DNA due to lack of a basic domain (*81*). Formation of heterodimers with HES/Her proteins inhibits their ability to bind to their own promoter and thus leads to de-repression of expression (*82*) and can result in prolonged expression even for low levels of Notch signaling (*83*). *nr2f1b* encodes a nuclear receptor, belonging to a family of transcription factors highly conserved throughout metazoan (*84*). In mice, deletion of *Nr2f1* leads to decreased expression of *Hes5* and to an increased sensitivity to gamma-secretase inhibitor treatment in the developing cochlea (*85*) . Moreover, it was recently shown that *NR2F1* can indirectly upregulate *HES1* expression in dormant human tumor cells, and *Fabp7* -another Notch target maintained in q5 (Fig. 4A)-is a direct target of Nr2f1 (*86*). Overall, this in silico analysis identified putative regulators with consistent expression patterns and data from the literature to support their potential role in promoting NSC resistance to LY treatment. We thus chose Id1 and Nr2f1b for further investigations.

### Nr2f1b is necessary and sufficient to promote resistance to Notch inhibition in adult NSCs

To determine whether Id1 and/or Nr2f1b mediate NSC resistance to Notch signaling inhibition, we first needed to identify potentially resistant NSCs *in situ* and to find ways to modulate *id1* and *nr2f1b* expression in a tractable and conditional manner. Because of the exclusive expression of *nr2f1b* in q5 qNSCs following a 24-hour LY treatment (Fig. 4D), we used *nr2f1b* as a marker of Notch-resistant NSCs. We subjected fish to a 24-hour DMSO or LY treatment, then assessed the expression of *ascl1a* and *nr2f1b* with smFISH in whole-mount pallia. We observed that, under physiological conditions, decreasing antero-posterior gradients of Pcna and *ascl1a* expression are anti-correlated with an increasing gradient of *nr2f1b* transcription (Fig. S5A). The induction of *ascl1a* expression after a 24-hour LY-treatment followed the same type of gradient with noticeably higher expression rostrally whereas *nr2f1b* remained enriched caudally (Fig. S5B), confirming that *ascl1a* expression occurs preferentially in territories with few *nr2f1b*^pos^ NSCs. Next, we extended Notch inhibition to 48 hours, which led to an increase in NSC proliferation measurable using Pcna immunohistochemistry, as previously observed (Fig. S6A) (*12*). Whole-mount preparations revealed that NSC activation (Pcna^pos^) displayed an anterior-high to posterior-low gradient mirroring the gradient of *ascl1a* expression observed after 24 hours of treatment. Since *nr2f1b*^pos^ NSCs are enriched caudally, we focused our analysis on this territory to compare them directly with their *nr2f1b*^neg^ neighbors (Fig. 5A). Quantification of the proportion of PCNA^pos^ NSCs after 48 hours of treatment showed that *nr2f1b*^pos^ cells expressed PCNA significantly less often than *nr2f1b*^neg^ NSCs (Fig. 5B). We further defined a response rate to report the likelihood for a given cell population to respond to Notch inhibition taking into account the proportion of proliferating cells in control and treated conditions. This revealed that the response rate of *nr2f1b*^pos^ cells is significantly lower than that of *nr2f1b*^neg^ cells (Fig. 5C), confirming the association between *nr2f1b* expression (q5 membership) and resistance to reactivation upon Notch blockade.

**Figure 5.**
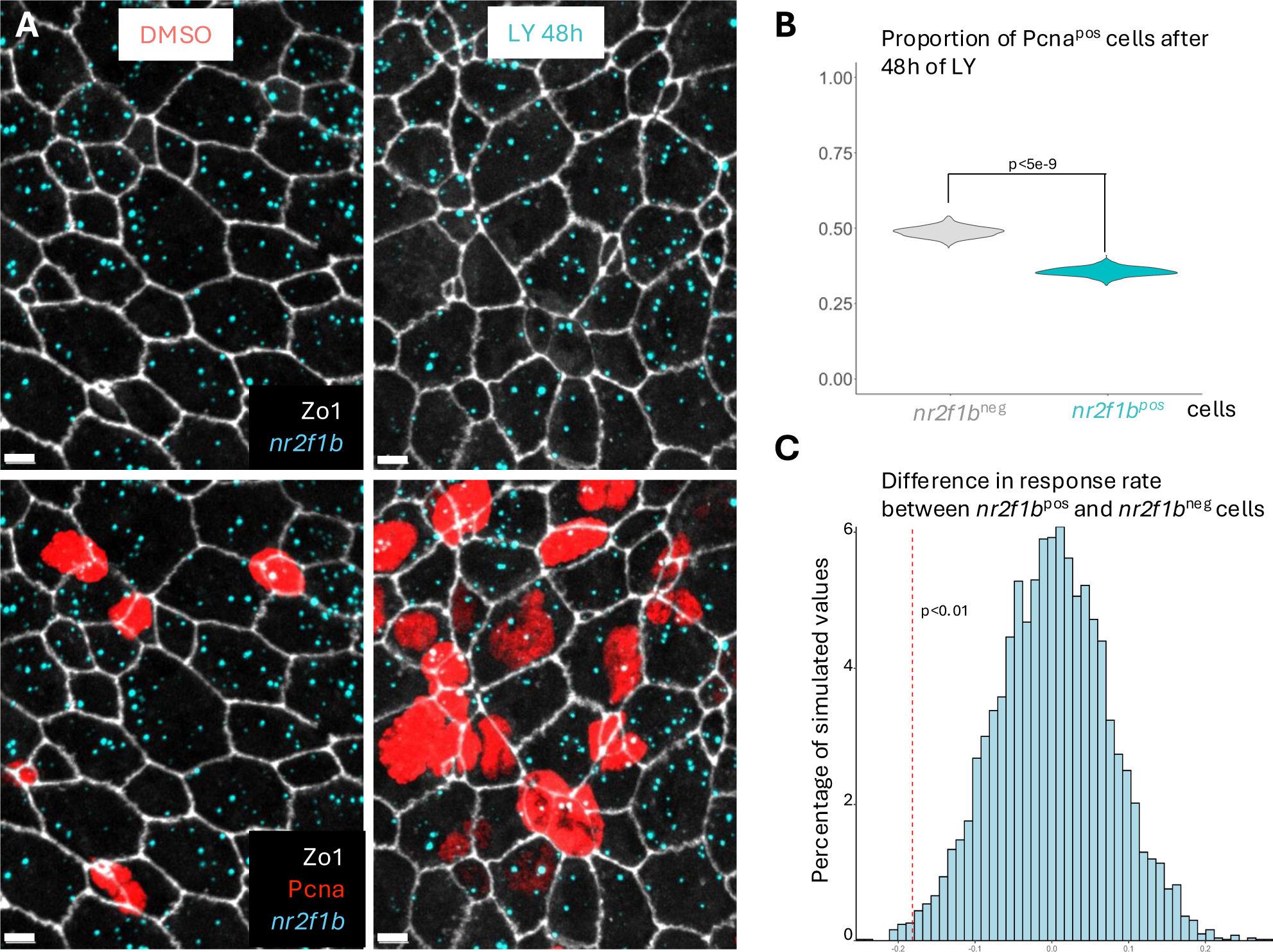
*nr2f1b* expression is associated with resistance to Notch inhibition. **A**. Confocal images of caudal areas in the dorsal pallium (see Fig. 1B) of fish treated for 48 hours with either DMSO or LY, immunostained for Zo1 (white, apical junctions) and Pcna (red, proliferation) and processed for smFISH for *nr2f1b* (cyan). Scale bars: 5 µm. **B.** Quantification of the proportion of cycling cells as a function of *nr2f1b* expression after 48 hours of LY treatment. *nr2f1b*^pos^ cells are less likely to be cycling than *nr2f1b*^neg^ cells. Reported p-value is derived from a chi square test. Violin plots are built from bootstrapped random sampling of the measured proportions to estimate the distribution but these estimated distributions are not used for statistical testing. DMSO: n=3 brains, 1263 cells; LY: n=3 brains, 1378 cells; *nr2f1b*^neg^ cells have zero *nr2f1b* mRNA dots. **C.** Difference in response rate (see definition of response rate in Methods) between *nr2f1b*^pos^ and *nr2f1b*^neg^ cells highlighting that *nr2f1b*^pos^ cells are less likely to respond to Notch inhibition. The histogram represents bootstrapped values to estimate a null distribution of the difference in response rates between *nr2f1b*^pos^ and *nr2f1b*^neg^ cells via Monte Carlo simulation. The dotted red vertical line represents the observed difference in response rate. 50 bootstrapped simulations with 1000 samples each were conducted, the p-value represented here corresponds to the maximum proportion of simulated values across all 50 bootstraps that were inferior to the observed difference.

We next proceeded to test the functional relevance of *id1* and *nr2f1b* under physiological conditions, focusing our analysis on the caudal part of the dorsal pallium.

*Id* genes expression is dependent on BMP signaling (*87*). *id1* expression in the zebrafish telencephalon is promoted by a conserved canonical cis-regulatory module containing a BMP-responsive element, and is inhibited by a treatment with the BMP signaling inhibitor DMH1 (*88*). The endogenous source of BMP in the zebrafish adult pallium was recently attributed to neurons (*89*), although other cells likely contribute. To decrease *id1* expression and assess whether this would affect NSC resistance to activation, we used DMH1 as previously reported (*88*). We treated fish from the *Tg(her4:dRFP)* line, a reporter of Notch signaling activity, with either DMSO, LY or DMH1 for 48 hours. Both LY and DMH1 treatments significantly decreased RFP intensity (Fig. S6A) consistent with the reported effect of DMH1 on *her4* expression (*89*) and demonstrating that DMH1 treatment was effective. In contrast, LY but not DMH1 led to an increase in NSC proliferation (Fig. S6A). We also compared proliferation after 48 hours of treatment with LY alone or joint treatment with LY and DMH1. This revealed that adding DMH1 does not change proliferation compared to LY treatment (Fig. S6B,C). Together, this suggests that the resistance of caudal NSCs to Notch pathway inhibition is not dependent on *id1* at least on this timescale.

We next tested the role of *nr2f1b*. In the absence of reliable ways to pharmacologically modulate its expression, we turned to the electroporation of fluorescently-tagged morpholinos (MO) in NSCs *in vivo*, as previously validated for several genes (*11, 12, 30*). A control MO, or a MO targeting *nr2f1b*, were intracranially injected into the pallial ventricle of 3mpf fish, followed by electroporation. This procedure primarily targets NSCs, which are in direct contact with the cerebrospinal fluid, and previous experiments demonstrated a maximal effect of MOs at 3-5 days post-electroporation (*11*). Electroporated fish were thus allowed to rest for 3 days before being treated with LY for 48 hours and analyzed (Fig. 6A), across the whole surface of the pallium. The proportion of proliferating cells among NSCs electroporated with the *nr2f1b* MO was significantly increased compared to control electroporated NSCs (Fig. 6B-C). Thus, *nr2f1b* downregulation decreases the ability of NSCs to remain quiescent when Notch signaling is inhibited.

**Figure 6.**
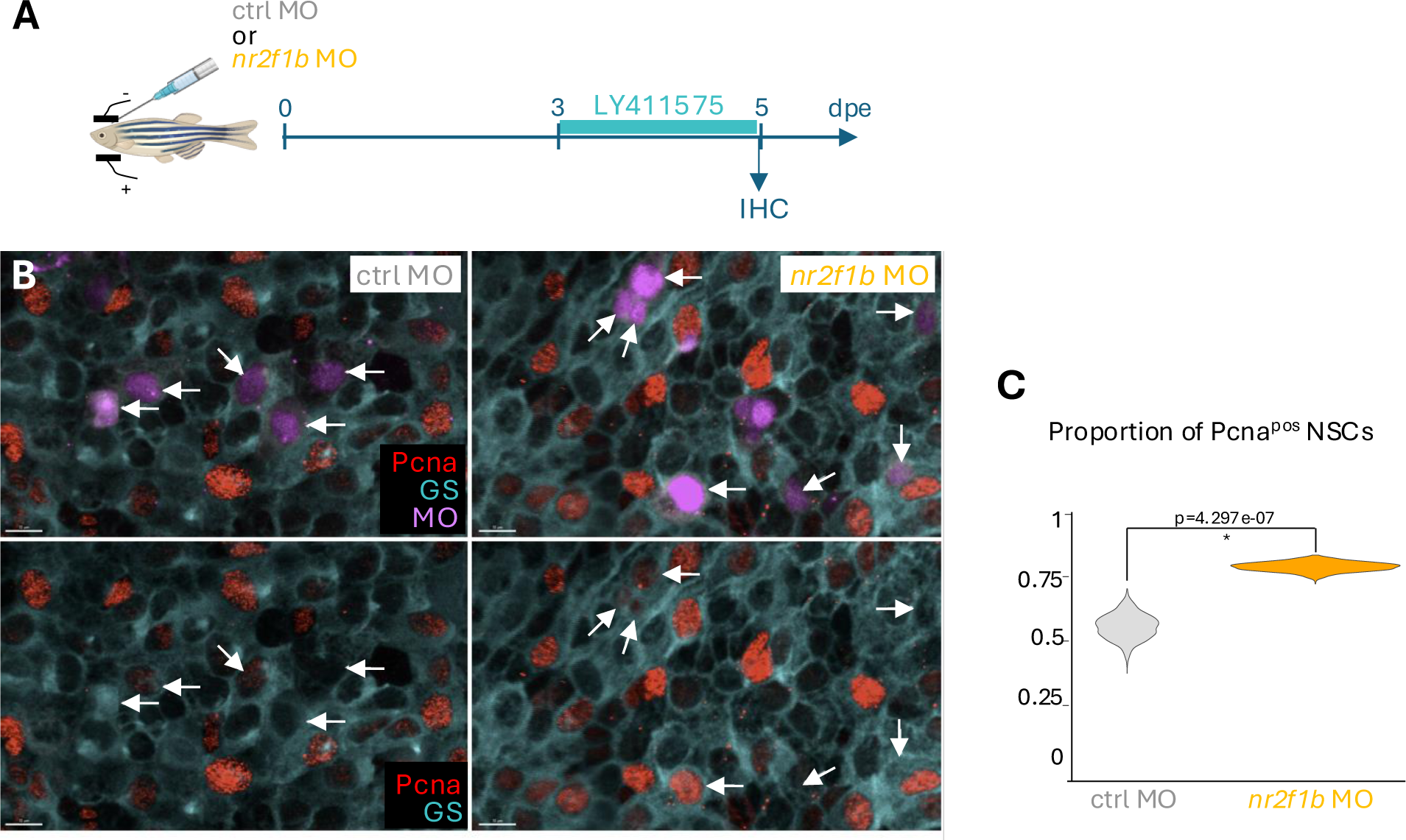
Endogenous *nr2f1b* expression is necessary for resistance to Notch signaling inhibition. **A.** Schematic of the experiment. After morpholino (MO) intracranial injection and electroporation, fish were allowed to rest for 3 days, then treated with LY for 2 days and sacrificed for quantification of the proportion of cycling NSCs. **B.** Example images of the pallial surface in fish electroporated with a control MO (left panels) and an *nr2f1b*-specific MO (right panels) followed by 48 hours of LY treatment (dorsal whole-mount views). The brains were processed for immunohistochemistry for Glutamine synthase (cyan, GS, labeling NSCs), Pcna (red, proliferation) and fluorescein (magenta, detecting the fluorescein-tagged MOs). Arrows point to electroporated NSCs. Scale bars: 10µm. **C.** Quantification of the proportion of proliferating NSCs as a function of the electroporated MO. NSCs electroporated with the *nr2f1b* MO are significantly more likely to proliferate after LY treatment than when electroporated with the control MO. Reported p-value is derived from a chi square test. Violin plots are built from bootstrapped random sampling of the measured proportions to estimate the distribution. Control MO: n=4 brains, 123 cells; *nr2f1b* MO: n=5 brains, 581 cells.

We next asked whether ectopically expressing *nr2f1b* would be sufficient to promote resistance to LY treatment. We electroporated 3mpf fish, as above, with a *pCMV:nr2f1b-P2A-nlsGFP* construct or a *pCMV:nlsGFP* control, allowed fish to rest for 3 days then treated them with LY for 24 hours to mimic the scRNAseq conditions (Fig. 7A). Here, we could make use of the slight leakiness of nlsGFP into the cytoplasm when highly expressed, allowing us to segment the NSC cell bodies and directly quantify the number of *ascl1a* dots per electroporated NSC with smFISH (Fig. 7B-C). This avoided the need for longer treatments to rely on Pcna expression as a proxy. This revealed that *nr2f1b* overexpression considerably hindered the upregulation of *ascl1a* after 24 hours of LY treatment (Fig. 7D-F). Thus, *nr2f1b* gain of function is enough to mediate NSC resistance to LY treatment, hence driving a cellular behavior similar to the behavior of q5 NSCs revealed with scRNAseq.

**Figure 7.**
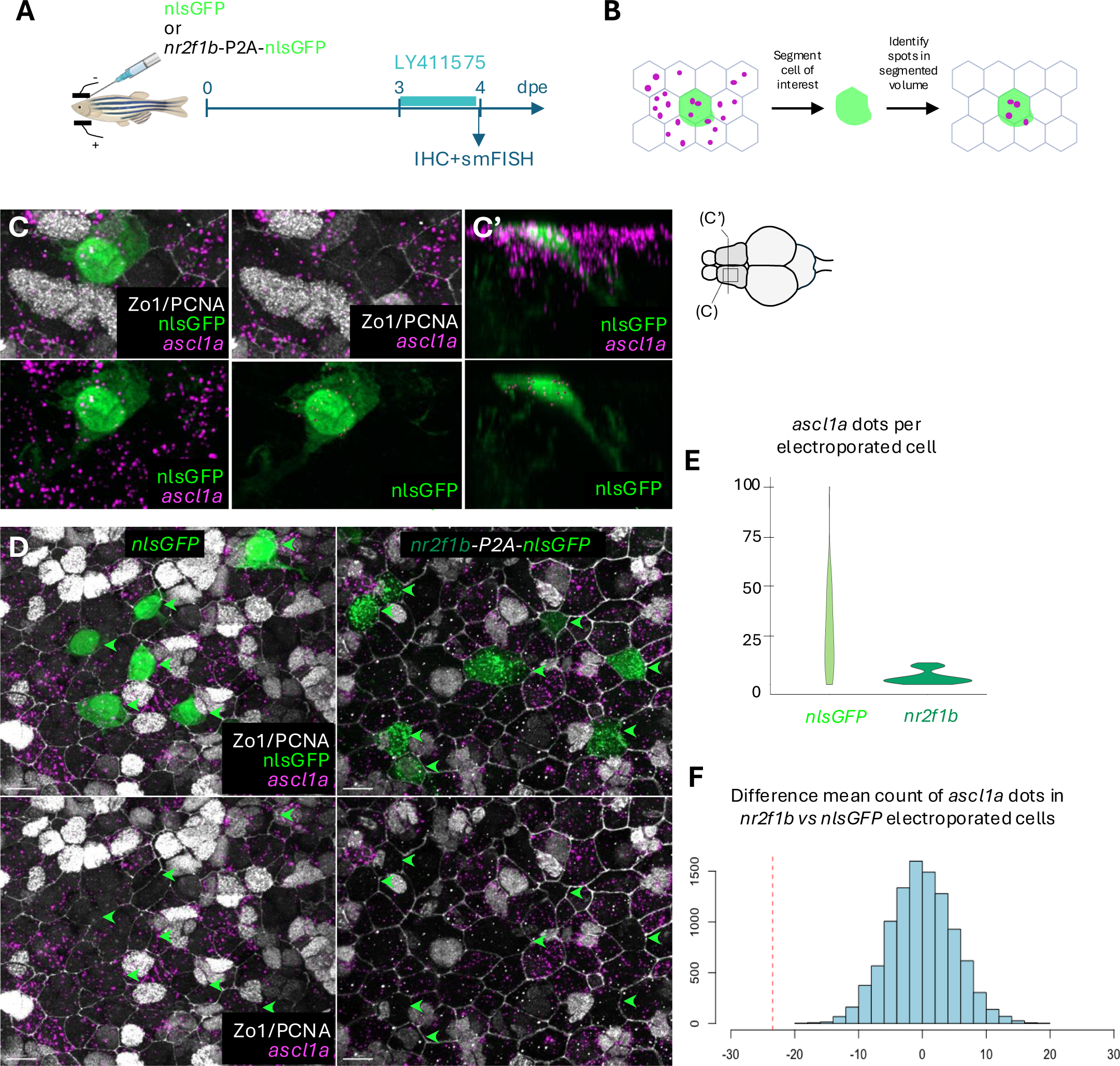
*nr2f1b* is sufficient to promote resistance to Notch pathway inhibition. **A.** Schematic of the experiment. After plasmid electroporations, fish were allowed to rest for 3, then treated with LY for 1 day and sacrificed for quantification *ascl1a* expression. **B.** Schematic of the approach used to quantify *ascl1a* molecules (magenta dots) in electroporated cells (green). NSC cell bodies do not always follow the shape of their apical membrane, precluding reliable automated quantification of signal with Zo1. However, the slight leakiness of nlsGFP leads to a filled cell body which can be segmented. Spots which fall into the the circumscribed volume are assigned to the cell in an unbiased way by detecting an elbow in the plot of pixel intensities in the *ascl1a* channel. **C.** Example of a cell with a complex morphology in which *ascl1a* dots were semi-automatically segmented. The first two images on top and the first image on the bottom show dorsal views of how the cell body extends beyond the Zo1^pos^ apical membrane and is surrounded in *ascl1a* dots. The third image on top shows an orthogonal view from the parenchyma displaying the complex morphology of the cell and the position of numerous *ascl1a* dots around it. The two last images on the bottom show the electroporated cells with only segmented *ascl1a* dots highlighted. Scale bars: 5µm. **D.** Example images of electroporated cells with *pCMV:nr2f1b-P2A-nlsGFP* or *pCMV:nlsGFP* after 24 hours of LY treatment. Arrows point to electroporated NSCs. Scale bars: 10µm. **E.** Violin plots of the number of *ascl1a* dots per cell in cells electroporated with *pCMV:nr2f1b-P2A-nlsGFP* or *pCMV:nlsGFP. pCMV:nlsGFP*: n=3 brains, 10 cells each; *pCMV:nr2f1b-P2A-nlsGFP*: n=3 brains, 10 cells each. **F.** Difference of mean test for the number of *ascl1a* dots in cells electroporated with *pCMV:nr2f1b-P2A-nlsGFP* or *pCMV:nlsGFP*. Values were randomly sampled 10000 times across all measures and assigned either an *nr2f1b* or control identity and the difference between the mean of counts assigned the *nr2f1b* identity with that of the counts assigned the control identity was computed. This generates a null distribution depicted by the histogram. It is centered on 0 and close to a normal distribution which confirms the validity of this approach to estimate the null distribution. The vertical line in red represents the true observed difference and is lower than any simulated value. Thus, cells electroporated with *nr2f1b* show significantly lower numbers of *ascl1a* mRNAs than control cells.

Nr2f1b can recruit co-repressors which are bound to Rbpj in the absence of Notch receptor cleavage and could thus lift the repressive effects of unbound Rbpj, leading to Notch-independent expression of Notch effectors (*90, 91*). This would be expected to maintain expression of dRFP driven by the *her4* promoter in *nr2f1b*^pos^ cells after LY treatment. However, we failed to detect an association between *nr2f1b* expression and RFP levels in *Tg(her4:dRFP)* fish after LY treatment (Fig. S6D). This suggests that *nr2f1b* acts through a mechanism or target sites independent of the Rbpj binding sites present in the promoter used for our *her4* transgenic line.

## Discussion

### Heterogeneity of adult quiescent neural stem cells

A major realization of the past few years has been that cells with different levels of stemness, fate restriction, responsiveness to external cues and proliferative activity coexist in adult stem cell niches in both zebrafish and mouse (*10, 92–94*). Differences in quiescence and long-term maintenance have been observed in NSCs in link with past proliferative activity (*3, 7*). Different transgenic lines also seem to label distinct populations (*5, 6, 10*) yet it hasn’t been possible to link them to specific molecular signatures. Here we combined an analysis at the single-cell level and an LY-induced phenotype with incomplete penetrance at the population level (*12*) to probe functional heterogeneity of adult NSCs with high resolution. This allows us to show that incomplete NSC activation at the population level cannot be not fully explained by a homogeneous activation probability resulting in partial response. Instead, NSCs are divided into subpopulations, some of which quickly respond to Notch pathway inhibition while others exhibit moderate to no response and a delay to enter the cell cycle. Our results further show that this behavioral heterogeneity is predictable based on scRNAseq cluster identity in control conditions (e.g., fast response of q2 and q3, hasted transition of q4a towards q1, no response of q5) adding support to the biological significance of these clusters, which we previously predicted to be associated with different quiescence depth.

In addition, LY treatment revealed cell states that we could not identify under control conditions. Specifically, cells belonging to a deeply quiescent cluster that transcriptionally resembles astrocytes (q4) are seemingly capable of advancing closer to the cell cycle without downregulating some of their marker genes. Whether this is a rare event that also happens under physiological condition or whether it requires a specific type of stimulus such as Notch inhibition is unknown. In the latter case, this would suggest that a form of “emergency” or rushed activation is possible given the appropriate stimulus, and that the topology of the quiescence phase is more complex than usually described. We also found that the latent NSCs described in several studies in mice are in fact likely to be the same population of medial striatal astrocytes and that the early phases of their reactivation, upon Notch blockade or injury, are similar to forced activation of q4 NSCs in zebrafish. Finally, our results suggest that zebrafish q4 NSCs may also participate in regeneration upon injury, ultimately losing their astrocytic markers in a manner reminiscent of murine latent NSCs. The zebrafish forebrain has long been established as a major model to gain insight into brain regeneration post-injury. Although this ability has been associated with an abundance of radial glial cells in zebrafish, these studies were conducted while being largely agnostic to the heterogeneous nature of zebrafish radial glia. Investigating the specific contribution of zebrafish astrocyte-like q4 NSCs in this process and comparing their path to activation with that of mammalian latent NSCs could further our understanding of regenerative neurogenesis and how to promote it.

### Distinct levels of Notch-dependency in adult neurogenesis

Although the importance of the Notch pathway in the regulation of adult neurogenesis and maintenance of the NSC pool has long been established (*29, 31, 95, 96*), the precise way in which it acts remains unknown on many levels. Here we could analyze molecular differences following Notch inhibition in all cell types of the neurogenic cascade simultaneously. In particular, while cycling IPCs and aNSCs are usually grouped together in a proliferating cell cluster, we could differentiate between them both in mice and zebrafish. Observing this, we found that progression through the cell cycle is accompanied not only by a change in ratios of expression between Notch receptors as previously described (*26, 31*) but also by a change in the relative expression of Notch effector genes. *Hes5* and its zebrafish orthologs are highly enriched in cycling NSCs over IPCs, while *Hes1* and its orthologs are only expressed at comparatively low levels. *Hes5* thus appears as an appealing putative intermediate for the maintenance of stemness in proliferating NSCs.

Notch signaling blockade has been shown to lead to both increased proliferation and neuronal differentiation, the latter being interpreted as the consequence of defects in self-renewal that take place in proliferating cells (*12, 26, 31*). Here we found that Notch inhibition already biases gene expression towards a molecular profile suggesting increased neuronal commitment even before NSCs start proliferating. This not only goes through upregulation of proneural genes such as *sox4a* but also through downregulation of astroglial genes. Direct regulation of astroglial genes by the Notch pathway has been demonstrated for *Blbp*/*Fabp7* and *Gfap* before (*97, 98*). Here we show that several other glial genes are downregulated quickly after Notch inhibition, including *metrn* which is a known regulator of astroglia identity (*49*). This suggests that in addition to promoting quiescence and repressing neuronal identity, the Notch pathway actively maintains glia identity in adult NSCs, which is consistent with its role in specifying astroglia in mammals (*99–101*), although we cannot say whether this effect is direct or dependent on intermediates. Of note, a parallel study aiming to characterize subpopulations of NSCs in the zebrafish telencephalon was recently published (*24*). Based on RNA velocity, the authors of this study mention two paths to neurogenesis from NSCs, either through a proliferative intermediate or through direct differentiation. They also sequenced a few cells after 48 hours of Notch inhibition. They were not able to integrate the control and treated datasets and instead relied on qualitative comparison of the two datasets analyzed separately. The main conclusion from this analysis was that a subset of NSCs, identified as being *snap25a*^pos^ and associated with direct differentiation, might be slightly enriched after Notch inhibition. We reanalyzed publicly available data from this study. Although we obtained qualitatively similar results, we were not confident in identifying *snap25a*^pos^ cells as directly differentiating NSCs. Thus, although the interpretation made in (*24*) is consistent with Notch having a pro-neurogenic effect as expected and as observed in our own data, our inability to replicate the results led us to not include a detailed analysis of *snap25a*^pos^ cells and direct differentiation in our own work.

### Co-operation of signaling pathways to ensure robust quiescence in a subset of NSCs

Contrary to most NSCs which quickly reactivated upon gamma-secretase inhibition, we revealed the existence of a subset of NSCs with low sensitivity to Notch inhibition. q5 NSCs were able to repress expression of *ascl1a* after 24 hours of LY treatment and proliferated less than other NSCs after a treatment of 48 hours. These cells could be less sensitive to signals from their neighbors and display delayed reactions upon pro-activation stimuli. Such properties would make them suited to act as a reserve, similarly to +4 cells in the intestinal crypts (*102, 103*). The existence of reserve zebrafish NSCs, which remain quiescent upon injury and can be identified by their upregulation of *id1* at 5 days post injury has been postulated previously (*89, 104*) . Although *id1* expression appeared highest in q5 NSCs, and was inferred as a predicted regulator of *her4.1* expression maintenance in these cells upon Notch blockade, our epistasis experiments suggest that in these conditions the action of BMP signaling does not contribute meaningfully to the resistance to Notch blockade. This might in part be due to our specific conditions. Indeed, a 5-day long treatment with DMH1 combined with LY treatment at a low concentration was shown to increase the effect on reactivation, and the additive effect of DMH1 is greater when the initial effect of LY is weaker (*89*). This is consistent with the role of Id proteins in modulating the stability of Notch effector genes: without Id proteins, Notch effectors can still function but their increased level of autorepression makes them more sensitive to even subtle changes in Notch signaling levels (*83*). In our case, because we used higher concentrations of LY (10µM instead of the 3µM which showed the maximum effect of DMH1), it is possible that the downstream effects are already maximized such that DMH1 can no longer have an additive effect.

Instead of Id1, we found that the resistance of q5 cells to Notch signaling inhibition is mediated at least in part by the transcription factor Nr2f1b. Its ortholog Nr2f1 is expressed in both mammalian telencephalic niches (*16, 105*) where it could play a similar role.

Of note, we also observed that among *nr2f1b*^neg^ cells, the ones in a caudal location are less sensitive to gamma-secretase inhibition than rostral NSCs. The choroid plexus, located in the posterior telencephalic region, is well situated to explain such gradients and secreted molecules from the choroid plexus are known to modulate NSC proliferation in the mouse SEZ (*106*). A scRNA-seq dataset encompassing cells from the tela choroida and the choroid plexus in the zebrafish brain (*22*) shows that besides BMP agonists, the choroid plexus also expresses high levels of Angptl1a -angiopoietin-like proteins being capable of acting as Notch agonists (*107, 108*)- and of the Wnt inhibitor Dkk3b -Wnt inhibitors promoting quiescence in murine NSCs (*109, 110*)-. Further studies will thus be required to get a clearer picture of the different levels of cooperation between signaling pathways to mediate the finely tuned spatiotemporal regulation of quiescence. Our results likely do not provide the full picture, also because we do not consider regulations happening at the protein level, as our identification of resistant cells is based on scRNAseq. Nevertheless our results suggest that Nr2f1 is able to promote a deep and refractory quiescent state through non-canonical modulation of Notch effectors. We note that Nr2f1 has been identified as a master regulator of glioblastoma and part of a core stemness module (*111*). It also induces dormancy in cancer stem cells from head and neck carcinoma and prostate cancer (*112*). Given the importance of Notch signaling in many cancers, the ability of Nr2f1 to promote a Notch-like phenotype in absence of canonical Notch signaling in adult NSCs suggests that it could be an efficient factor for cancer stem cell maintenance.

## Methods

### Fish lines and maintenance

All procedures relating to zebrafish (*Danio rerio*) care and treatment conformed to the directive 2010/63/EU of the European Parliament and of the council of the European Union. Zebrafish were kept in 3.5-liter tanks at a maximal density of five per litter, in 28.5°C and pH 7.4 water. They were maintained on a 14-hour light/10-hour dark cycle (light was on from 8 a.m. to 10 p.m.) and fed three times a day with rotifers until 14 days post-fertilization and with standard commercial dry food (GEMMA Micro from Skretting) afterwards. All fish used for experiments were between 3 and 4 months old. The transgenic lines *Tg(her4.1:dRFP)* (*113*) and *Tg(sox2:GFP)* (*36*) were maintained on an AB background.

### Pharmacological treatments

LY411575 (Sigma Aldrich) and DMH1 (Tocris) were resuspended in pure DMSO at 100µM and stored at -20°C and kept hidden from the light. For treatments, fish were transferred to new tanks at a density of one fish per 50mL of fish water. LY411575 was diluted to a final concentration of 10µM and DMH1 to a final concentration of 20µM. Regular DMSO was used as control for all experiments and to adjust total DMSO concentrations between conditions when necessary. Because LY411575 has been reported to be light sensitive, all treatments were carried out in the dark for all conditions. For treatments lasting over 24 hours, fish were given food and allowed to eat for 1 hour before their water was renewed.

### Euthanasia

Fish were euthanized in ice-cold water (temperature comprised between 1° and 2°C) for 10 min, according to a special dispensation and following the guidelines of the Ministry of Superior Education, Research, and Innovation.

### Dissociation and cell sorting

Cell sorting was conducted three days in a row to collect replicates, using twenty 3-month old adults from the *Tg(sox2:GFP)* line (*36*) on each day after 24 hours of treatment with LY411575. DMSO-treated brains (reported in (*1*) and serving as the control dataset) were processed at the same time. Brains were dissected in Ringer’s solution. The telencephalon was separated from the midbrain and the olfactory bulbs were removed. The two hemispheres were separated and cut along the boundary between pallium and subpallium to enrich for pallial cells. Cell dissociation was carried out according to (*114*). Cells were then sorted on a FACSAria III. We used forward and side scatter and DAPI staining to distinguish live cells from debris and sorted cells on their GFP levels to enrich for *sox2*^pos^ cells while still including GFP^neg^ cells to not miss any relevant population. After sorting, encapsulation of cells and reverse transcription was immediately performed using the 10x Chromium Controller and Chromium Single Cell 3’ Kit v2 and then immediately frozen at -80°C until all replicates had been collected.

### Library preparation and sequencing

After reverse transcription all replicates were processed in parallel with the 10x Genomics v2 Kit. Afterwards, barcoding libraries were pooled and split over multiple lines of a HiSeqX and sequenced at a depth over 100k reads per cell using 2x150 paired-end kits with the following sequencing read recommendations: Number of Cycles: 26 cycles Read1 for cell barcode and UMI, 8 cycles I7 index for sample index and 98 cycles Read 2 for the transcript. This yielded a saturation above 95% for all libraries.

### Mapping and filtering of data

Initial analysis was conducted using the cellranger software (https://support.10xgenomics.com/single-cell-gene-expression/software/pipelines/latest/what-is-cell-ranger). Reads were demultiplexed and mapped to the GRCz11 zebrafish genome assembly from Ensembl. Datasets were first analyzed individually to determine whether they were fit for integration. First, we filtered cells based on the number of genes (nGene) and UMIs (nUMI) detected as well as on the relationship between nGene and nUMI. nGene is expected to be positively correlated with nUMI. Lower nGene than expected for a given nUMI value suggests low library complexity whereas higher nGene than expected for a given nUMI value suggests excessive library complexity and likely doublets. To filter on that criterion, we fit a loess regression curve for nGene∼nUMI with a span of 0.5 and of gaussian family and removed cells which residuals were beyond three standards deviation of the mean (*115*). Then we inspected the frequency of cells associated with a given nGene. This yielded a distribution with a narrow peak at low levels of nGene followed by a broader distribution centered around a peak close to nGene=900. We considered the first peak to be low quality cells and set a threshold to remove it from the rest which ended up being nGene=200. We also removed cells which had abnormally high nGene or nUMI based on the distribution on a plot of nGene as a function of nUMI. Then we removed genes expressed in fewer than 10 cells. Finally, we computed the percentage of the transcriptome that consisted of mitochondrial genes (percent.mito) and inspected plots of percent.mito as a function of nGene. These two variables show negative correlation because high percent.mito tends to be observed in low quality cells, in which fewer genes are detected. We set a threshold at 10% which removed a tail of cells with high percent.mito and low nGene. Subsequent inspection of the variation of gene expression as a function of technical variables such as nGene after regular scaling and normalization revealed that variance in gene expression was not dependent on technical factors.

Post-filtering we retained 17710 control cells and 18776 LY-treated cells. Analyses restricted to qNSCs included over 6000 cells.

### Identification of scRNAseq clusters and their markers

There was a very small batch effect between the replicates which could be corrected by simple linear regression and we thus did not use any additional batch correction method. Variable genes were identified by selecting genes exhibiting high variability given their level of expression without using a hard general threshold. We selected PCA components to include based on whether they explained over 1% of the variance across the first 100 principal components and whether that variance was flagged as significant by the JackStraw test. Initial clustering was performed on all the cells using a simple Smart Local Moving algorithm (*42*). We looked for and removed doublet clusters using three separate approaches. The doublet cells methodology from the scran package was used to score individual cells and the doublet cluster methodology to detect clusters which look like a mixture of two other clusters. An adapted DoubletFinder algorithm was used to generate doublets from randomly selected pairs of cells and detect cells which frequently co-clustered with such cells (*116*). This multi-pronged approach ensured that we did not keep any artefactual cluster at this stage. Several broad cell types were already subdivided into multiple clusters. We isolated each individual lineage and performed further clustering to identify fine-grained cell subpopulations. Substantial heterogeneity was apparent in neurons and radial glia. To ensure that we did not overcluster, we developed a consensus clustering approach. We pooled results from Bayesian model mixture, smart local moving algorithm, multilevel algorithm, walktrap, spinglass and density peaks. kNN graphs used for the graph clustering approaches were themselves built using edges weighed either as in SNN-Cliq (*43*) or Phenograph (*40*). From this we obtained a consensus matrix with the Cluster-based Similarity Partitioning Algorithm (*39*). From this consensus matrix we drew a final hierarchical clustering and used a conservative cutoff to identify robust clusters. We then subjected each pair of neighboring clusters to differential gene expression detection to determine whether they should be merged.

### Data Integration

We performed data integration using the rliger package (*46*). It implements an integration method based on iterative non-negative matrix factorization described in (*45, 46*). This non-negative matrix factorization identifies both dataset-specific and shared metagenes and uses the latter to generate a joint low-dimensional embedding. Liger compares favorably to other integration methods (*117*), produces interpretable results thanks to the use of non-negative matrix factorization, and relies on few parameters, facilitating grid-search. We performed a grid search on the lambda and k parameters to optimize the integration, settling on a lambda of 1.25 with a k of 18 for qNSCs and a lambda of 5 with a k of 8 for the aNSCs. Clusters with a strong identity, in particular those that are regionally defined and can thus be reliably identified, were used as controls to control for incorrect and in particular excessive merging of the datasets. Following dimensional reduction with non-negative matrix factorization the integrated dataset was subjected to the same type of analysis as individual datasets, but clustering and calculation of UMAP and tSNE were performed on the integrated embedding rather than on principal components.

### Differential abundance of scRNAseq clusters

Detecting different proportions in single cell RNAseq studies is not straightforward. Comparisons of proportions based simply on calculating an average between replicates does not take advantage of the number of cells being profiled, whereas reporting a single point estimate of proportions across all cells does not provide a statistical assessment of whether differences are meaningful or not. We calculated differential abundance based on a bootstrapped difference of proportion test. Briefly, for both conditions and for each cluster, we randomly sampled with replacement a number of entries equal to the number of cells of that condition which can either belong to the cluster or not with a probability that is equal to the proportion of the cells of that condition that belong to that cluster. We then calculated the difference in simulated proportions for an arbitrary number of times. This results in a vector of simulated proportion difference which is expected to reflect what would be observed if the single cell RNAseq was repeated many times. If there are no differences, we expected the distribution of simulated proportions to be centered on 0. We calculated the differences in proportions as LY – Control and used 10000 simulations. We then used the Bonferroni method to correct for multi-testing.

### Calculation of cell cycle scores

G1/S and G2/M scores were determined from scRNAseq data using the method developed in (*118*) and implemented in the Seurat package. First, genes belonging to different phases of the cell cycle were curated using Cyclebase3.0 (*118*) and orthologs in mouse and zebrafish were identified using the ortholog tables that we previously generated (*1*). Next, the average expression of these genes was computed across all cycling cells and used to divide genes in 24 bins of similar average expression. For each gene belonging to a geneset of interest, 100 genes belonging to the same average expression bin were randomly selected ensuring that the set of control genes would have a similar overall distribution of levels of expression. Finally, a score was attributed to each cell based on the difference between the average expression of the genes belonging to the G1/S or G2/M scores and the average expression of control genes.

### Identification of putative regulators of q5 resistance to Notch blockade

The list of putative regulators was restricted to all zebrafish genes known or predicted to encode proteins with a transcription factor activity. We filtered the data matrices of both control and LY treated datasets to retain only genes expressed in at least 1% of all cells or 10% of the cells of at least one cluster. We then obtained weighted matrices linking transcription factors to potential targets using GRNBoost2’s python implementation (*79*). When implemented as part of the SCENIC pipeline this step is usually followed by refinement of inferred gene modules, by determining whether the promoter of putative targets contain sequences close to the binding motif of the associated transcription factor (*119*). However, this method is only available for drosophila, mice and humans (*119*) as generalization of the cis-regulatory motif scoring approach was not successful in zebrafish (S. Aerts, pers. comm.). Instead, based on the empirical observation that, for a given target, only a few putative regulators have a high associated weight and that this weight then plateaus at low level for many regulators, we devised a method based on automatically detecting the elbow point in weight values to refine gene modules. Rather than doing it based on an arbitrary threshold on the number of regulators to keep, this selection method flexibly and automatically detects the break point in a plot of sorted weight and retains all the putative regulators with higher weights.

### Preparation of tissue for immunohistochemistry

Whole brains or whole telencephala were dissected from 3-to 4-month-old fish after euthanasia as described above in cold PBS. They were immediately placed in cold 4% PFA and fixed on a rotating platform at 4°C overnight. The next day, brains were dehydrated through sequential washes in 25%, 50%, 75% and finally 100% methanol (mixed in PBS the first three solutions) and then stored at -20°C. For labelling, brains were first rehydrated through washes in 75%, 50% and 25% methanol in PBS and finally PBS + 0.1% Tween 20. They were subsequently treated with a bleaching solution made up of 0.5X SSC, 3% H202, 0.05% formamide diluted in DNAse and RNAse-free water and then washed four times in PBS + 0.1% Tween 20.

For immunostaining, brains were then treated with histoVT 1x at 65°C for one hour for antigen retrieval, incubated in PBS + 0.1% Tween 20 + 5% normal goat serum + 1% DMSO and finally incubated overnight at 4°C in the same buffer with primary antibodies. The next day they were washed in PBS + 0.1% Tween 20 five times before being stained with the secondary antibody. The antibodies used are listed in Table S4.

For RNAScope staining (https://www.bio-techne.com/reagents/rnascope-ish-technology), after bleaching and washing in PBS + 0.1% Tween 20, brains were pre-incubated at 40°C in a solution containing 5X SSC, 25% formamide, 0.1% Tween 20, 50µg/µL of heparin and 2.5mM of citric acid for at least one hour. They were then incubated overnight in the same conditions with primary probes from the RNAScope Hiplex kit and subsequently processed according to the kit’s instructions. Afterwards immunostaining was described as above starting from the incubation with the blocking buffer. Information on probe sequences and detection channels used are listed in Table S5.

### Image acquisition and processing

Whole mount images were acquired on LSM700 and LSM710 laser-scanning confocal microscopes with 40x oil objectives with numerical apertures equal to 1.4 and 1.3 respectively. Images were then stitched with ZEN’s proprietary software and analyzed with Fiji or Imaris. Imaris’ in-built function were used to segment GFP surfaces and identify spots for RNAScope.

### Quantification of Pcna^pos^ cells

In the case of experiments without electroporations, square regions in the xy plane, with a side length between 125µm and 150µm, were selected in the indicated area (either rostral or caudal). The number of PCNA^pos^ cells at the surface of the ventricle was taken to reflect the number of proliferating NSCs and divided by the total number of intact NSCs in the region of interest to obtain the fraction of proliferating NSCs.

In the case of experiments with electroporations, counting was performed among cells labeled by the MO or the plasmid and present at the ventricular surface.

To assess the response of cells to LY and compare this response in *nr2f1b*^pos^ cells and *nr2f1b*^neg^ cells, we calculated the percentage of proliferating cells in each population and each condition but also defined a response rate, as a way to directly illustrate the effect of LY treatment on each population. Rather than directly using the ratio of PCNA^pos^ cells between LY and DMSO treatments in each population, the response rate takes into account the number of cells “at risk”, drawing inspiration from a widespread approach in the medical field and is defined for each population as (%PCNA^+^_(LY)_-%PCNA^+^_(DMSO)_)/%PCNA^-^_(DMSO)_. This way, the response rate reflects the proportion of cells that were recruited by LY treatment divided by the population of cells that could be recruited. This allowed us to circumvent the drawbacks of directly calculating a ratio between the proportion of PCNA^pos^ cells in LY-vs DMSO-treated fish. Indeed, such an approach is not suited when baseline percentages differ significantly as is the case here. For example, an absolute gain of activated cells of 5% in two populations with a baseline proliferation of 5% and 15% respectively results in ratios of 3 and 1.33 respectively even though a larger proportion of the cells that had the potential to be activated responded in the second population. The response rates take this into account. Because we could not conduct paired experiment to directly measure the response rate, as that would have necessitated reading out PCNA immunostaining and *nr2f1b* expression levels before and after treatment, we estimated it using a Monte Carlo simulation from separate counts on control and treated cells.

### Cloning

The full coding sequence of *nr2f1b* was amplified via PCR with Phusion Plus polymerase and cloned to generate a *pCMV:nr2f1b-P2A-nlsGFP* from *pCMV:hey1-P2A-nlsGFP*. *pCMV:nlsGFP* and *pCMV:hey1-P2A-nlsGFP* had been generated previously (*30*).

### Ventricular injections and electroporations

For electroporations, fish were first anesthetized in 180mg/L of MS222/Tricaine. A hole was opened in their skull with a sterile needle and a capillary introduced for intraventricular injection. They were then placed in a separate dish with their head between electrodes and administered 4 electric pulses for electroporation (50 V, for 50ms each and separated by 1 second). They were then put in an individual dish with 3g/L of NaCl in fish water and they received frequent gentle pulses of water through the mouth to help with waking up.

To selectively block Nr2f1b protein production, we electroporated a previously validated lissamine-tagged splice-blocking morpholino (*120*): 5’ - CCCACACAAGATGTACTCACCTTCG - 3’ . A lissamine-tagged morpholino which does not target any zebrafish gene was used as control: 5’ - AGAGCAACTGAACTCACTCACGTTC - 3’. In both cases we used a 1mM concentration.

For gain of function experiments the same procedure was followed with the *pCMV:nr2f1b-P2A-nlsGFP* and *pCMV:nlsGFP* plasmids at 0.15fmol/µL.

### Pallial lesions

For lesions fish were anesthetized in the same way as for electroporations and a BD Microfine 29G needle with a 0.33mm diameter was inserted through the skull until all of the beveled end of the needle had penetrated the fish’ head. Fish were immediately moved to a new tank without necessitating a specific treatment to increase survival odds as baseline survival rates were already 100%.

### Statistical analysis

All statistical analyses were performed in R. For comparisons between three groups as in Fig. 1F we used a kruskal-wallis test. Comparisons between two sets of proportions were subjected to chi square tests. For the number of spots per electroporated cells we used a bootstrapped difference of the mean test. For the statistical test comparing distribution of values in different deciles in Fig. S4H we used the rogme package. All countings included at least 3 hemispheres, and exact numbers are provided in figure legends.

## Supporting information

Full supplementary material

## Acknowledgments

We thank the ZEN team for input, Nicolas Dray for his critical reading of the manuscript, and Sébastien Bedu and Nathan Guibert for expert zebrafish maintenance.

## Funding

Work in the L. B-C. laboratory was funded by the ANR (Labex Revive), La Ligue Nationale Contre le Cancer (LNCC EL2019 BALLY-CUIF), the Fondation pour la Recherche Médicale (EQU202203014636), CNRS, INSERM, Institut Pasteur and the European Research Council (ERC) (SyG 101071786 - PEPS).

## Author contributions

Conceptualization: DM, LBC; Methodology: DM, AA, IF; Funding acquisition: LBC; Project administration: LBC; Supervision: LBC; Writing – original draft: DM; Writing – review & editing: DM, LBC.

## Competing interests

Authors declare that they have no competing interests.

## Data availability

All sequencing data generated in this study have been deposited in the GEO database and are available at: https://www.ncbi.nlm.nih.gov/geo/query/acc.cgi?acc=GSE225863 for the control cells and at https://www.ncbi.nlm.nih.gov/geo/query/acc.cgi?acc=GSE274483 (reviewer token: **czqpcoyarrsbzyl)** for the LY-treated cells . Additional processed data have been deposited in the French data.gouv database and are available at: https://entrepot.recherche.data.gouv.fr/privateurl.xhtml?token=2ed67607-b912-4143-8021-ff3e82173d74. All other data needed to evaluate the conclusions in the paper are present in the paper and/or the Supplementary Materials.

## Notes

### Competing Interest Statement

The authors have declared no competing interest.

