## Supplementary material for "Notch signaling blockade links transcriptome heterogeneity in quiescent neural stem cells with their reactivation routes and potential": Full supplementary material

**Morizet et al.**

### Supplementary information

This file includes:

- Supplementary text: Reclassification of qNSC1 striatal astrocytes
- Legends for Supplementary Figures
- Legends for Supplementary Tables
- References for Supplementary data
- Supplementary Figures
- Supplementary tables

### Supplementary text

#### Reclassification of qNSC1 as striatal astrocytes

We reanalyzed the data from the original mouse study (*16*) with a pipeline developed by the Allen Institute suited to Smart-Seq2 data with few cells (*120*), and attempted to assign identities to the resulting clusters. The number of clusters and the classification of oligodendrocytes, neuroblasts and proliferating cells were in good agreement. However, we found that one of the quiescent astroglial clusters expressed high levels of genes associated with astrocytes rather than radial glia-like (RG-like) cells. We also re-analyzed another dataset which collected more cells and reported an identical subdivision of the data (*15*) and obtained the original annotations. Once again, we recovered clusters comparable to the published data but found that the cells initially annotated as qNSC1 had a transcriptional profile similar to that of astrocytes. In particular, they expressed high levels of *Aqp4*, *F3* and *S100b,* which are commonly used to distinguish astrocytes from RG-like cells, as well as *Timp4*, *Cxcl14* and *Slc4a4,* which we recently described as being enriched in astrocytes rather than RG-like cells in mouse (*1*). Several independent studies have now generated scRNA-seq atlases of the SVZ without explicitly reporting the presence of the qNSC1 cluster (*13*, *17*, *19*, *59*, *60*). We re-analyzed all of them to determine whether they contained clusters of cells matching the transcriptional signature of qNSC1 cells and how these cells had been classified. We ultimately excluded the data from (*17*) because their study was tailored to recover a large number of cells at the expense of library complexity and thus risked lacking sensitivity to distinguish between closely related cells. Shah et al. did not report the presence of astrocytes in their datasets but labeled qNSC as aNSC as evidenced by the pattern of expression of *Thbs4* (*60*). The large cluster of cells they labeled as qNSC expressed the same astrocytic genes as those mentioned above, thus their classification into qNSC and aNSC matched the qNSC1 and qNSC2 clusters from (*16*) and (*15*). On the other hand, both (*19*) and (*13*) reported the presence of astrocytes alongside NSCs and our re-analysis confirmed that these astrocyte clusters matched qNSC1 (Fig.S4B). Cebrian-Silla et al. (*13*) profiled a large number of cells with high gene detection per cell and identified several spatially distinct populations of qNSC which were closer to each other than to the astrocyte cluster and we independently reproduced the results from this analysis. Overall, this strongly suggests that the cells that qNSC1 cells are in fact astrocytes. We also recovered data from a whole telencephalon atlas (*63*) and used MetaNeighbor (*62*) to map the astroglial cells from (*15*) to those of the atlas. In this analysis, qNSC1 cells clustered together with telencephalic astrocytes and away from NSCs, supporting their astrocytic identity (Fig.S4C). A latent neurogenic potential, activated upon stroke, injury or Notch inhibition has been reported in striatal astrocytes (*8*) in particular in the medial striatum, close to the lateral wall of the SVZ. We thus asked if qNSC1 cells might be striatal astrocytes and re-analyzed data from a study on striatal astrocyte (*67*). Our analysis largely agreed with the reported clusters and we found that a subset of cells from the striatum of P90, but not P3 mice bearing a striking similarity to qNSC1 cells. One cluster in particular displayed higher expression of *Crym,* which is also enriched in qNSC1 over qNSC2 and is expressed in astrocytes in the medial striatum (*13*), and lower expression of several astrocytic genes such as *Agt*, *Sparc* and *Nnat*. Moreover, we reanalyzed scRNA-seq data generated simultaneously in cortical and striatal astrocytes (*58*). This confirmed that striatal astrocytes are enriched in *Cd9* and *Cd81* compared to cortical astrocytes, and thus that the differences that led to the classification of qNSC1 as RG-like cells might be linked with heterogeneity among astrocytes instead (Fig.S4D). Thus, we concluded that cells originally classified as deeply quiescent RG-like cells are in fact a population of astrocytes from the striatum, likely enriched medially and which emerges between P3 and P90.

A recent multi-omics study including qNSC1 and qNSC2 was recently published (*57*), in which the authors conclude that the methylome of the cells classified as qNSC1 was closer to that of control astrocytes than to that of RG-like cells. Although they renamed them “vSVZ astrocytes" they also proposed that these cells might be B1 cells, which are RG-like cells. A recent report characterized both B1 and B2 cells in the SEZ (*64*), and the markers identified as labelling them belong to the cells that were previously classified as qNSC2 and which can also be mapped throughout multiple datasets according to our analyses. Thus, so-called qNSC1 and qNSC2 can be reliably distinguished not only via their methylome but also via their transcriptome, and qNSC1 cells are actually astrocytes while qNSC2 cells are bona fide RG-like cells. Importantly this does not call into question the functional properties that had been identified for these cells, except for the interpretation that qNSC1 cells reactivate by becoming "qNSC2" cells. Indeed, we consider it highly unlikely that parenchymal cells turn into RG-like cells. Instead it is more likely that they go through a state with a transcriptome similar to that of NSCs while remaining parenchymal astrocytes, as had been previously postulated (*58*). Overall this interpretation reconciles multiple studies on the SEZ with hitherto diverging results, unifies the description of reactive neurogenesis and is supported by methylomic as well as both regular and spatial (*61*) transcriptomic data and consistent with in situ validation with RNAScope (*64*). For these reasons, we believe that the initial distinction between qNSC2 and qNSC1 should instead be treated as a distinction between RG-like cells and local astrocytes, respectively, the latter likely being the same population as the population of medial striatal astrocytes previously identified as latent NSCs, which is how we considered them in our final analyses.

### Legends for Supplementary Figures

**Figure S1 - *ascl1a* is upregulated but major qNSC subpopulation markers display similar patterns following 24 hours of DMSO and LY treatment. A.** Low dimensional embedding of qNSCs in a DMSO control scRNAseq dataset colored by cluster (same as in Fig.1H, from (*1*))**. B.** Low dimensional embedding of qNSCs in the LY-treated scRNAseq dataset colored by cluster (as in Fig.1I)**. C,D.** Levels of expression of *ascl1a* in the DMSO (C) and LY (D) datasets. **E.** Levels of expression of cluster-indicating genes in the control dataset. Rare clusters identified by *nkx2.1* (q7) and *gsx2* (q6) are circled to highlight them, and q5 is indicated by an arrow. **F.** Levels of expression of the same genes in the LY-treated dataset.

**Figure S2 - Integrative analysis highlights novel qNSC subclusters and differentially expressed genes between LY-treated and untreated qNSCs in each cluster**. **A.** Low dimensional embedding of the joint qNSCs scRNAseq dataset colored by condition (control in red and LY-treated in blue). **B,C.** Projections of individual datasets (B: DMSO, C: LY) on the low-dimensional embedding color-coded by cluster. This highlights that the two datasets thoroughly mix but that several clusters remain clearly identifiable. **D,E.** Heatmaps showing, for a selection of genes, relative mean gene expression per cluster in the DMSO (D) and LY (E) datasets. Because expression levels differ between genes, a scaling per gene was applied (for each gene, an expression level equal to the mean of expression in all clusters is set to zero, and clusters with expression higher or lower than the mean are positive or negative, respectively). **F.** Dotplot of a subset of differentially expressed genes between LY-treated and untreated cells in each qNSC cluster. A subset of genes known to be Notch targets (among which Notch effectors are highlighted in yellow) and/or associated with glial identity (highlighted in green), proneural activity (highlighted in orange), cell cycle entry (highlighted in pink) or chromatin remodeling (other genes) was chosen for visualization. Dot size is binary based on whether the adjusted pvalue of the MAST differential gene expression test with nUMI as a latent variable is below 0.05 or not. Dot color is determined by log2 fold change, with purple indicating upregulation in LY-treated cells and green indicating downregulation in LY-treated cells.

**Figure S3 - Analysis of cycling cells in the adult mouse SEZ and zebrafish pallium reveals comparable NSCs, IPCs and cell cycle phases**. **A.** Projection of the cycling cells of each individual zebrafish pallial dataset (DMSO-treated, top (*1*), and LY-treated, bottom -this study-) on the integrated embedding of cycling cells, color-coded by cluster derived from integrated analysis. Note the higher total number of cells in the LY-treated dataset. **B.** Percentage of proliferating cells per cell category (from top to bottom: all cells, NSCs and IPCs) in the DMSO versus LY datasets (color-coded). **C.** Low dimensional embeddings of cycling cells from the mouse SEZ (left) (from (*2*)) and zebrafish pallium (right) (from (*1*)) colored by levels of expression of genes enriched in cycling NSCs (*Ckb*, *Mfge8*) or IPCs (*Elavl3*, *Sox11*) and cell cycle scores. Color coding is distinct to separate genes from scores. In both cases two independant sources of variation can be identified transcriptionally and visualized on the lower dimensional embedding: an axis (horizontal) separating NSCs from IPCs and an axis (vertical) going from cells in early phases of the cell cycle to cells in late phases of the cell cycle. **D.** Plot of log2 Fold Change in zebrafish NSCs over IPCs in relation to log2 Fold Change in murine NSCs over IPCs after conversion to a common set of orthologs. A majority of genes that are detected as significant in both datasets fall in correlated quadrants. Among them, *Hes5* is the only effector of the Notch pathway with significantly higher expression in NSCs than in IPCs in both zebrafish and mice. **E.** Low dimensional embedding of the joint scRNAseq datasets of cycling cells for the DMSO- and LY-treated conditions, color-coded by condition (top left) (DMSO: red; LY-treated: cyan), by cluster (top right) (the two datasets thoroughly mix but)several clusters remain clearly identifiable., or showing only the cells of the DMSO dataset (bottom left) or the cells of the LY dataset (bottom right).

**Figure S4 - Zebrafish astrocyte-like NSCs and murine latent NSCs share features when forcefully reactivated by Notch inhibition or injury**. **A.** Left and middle panels: log2 Fold Change (log2FC) of the expression of ribosomal genes detected as significantly differentially expressed between LY-treated and untreated cells in clusters q1e (left) and q1f (middle). Cell transition to q1e and q1f upon LY treatment is not accompanied with a normal upregulation of ribosomal genes. Right panel: log2 Fold Change of the expression of ribosomal genes between cells found in "qNSC2" (see Fig.S4B) after injury and unperturbed "qNSC2" cells (*3*). Ribosomal genes were not sorted for statistical significance in this case because of the low power of statistical tests with as few cells as were profiled in these scRNAseq experiments with Smart-seq technology. **B-D.** Reanalysis of mouse scRNAseq datasets profiling latent neural stem cells, variably classified as dormant NSCs (qNSC1 in (*3*)), as the only qNSCs(*4*) or as astrocytes (*5*). Detailed reanalysis of these datasets, mapping to a murine telencephalon atlas encompassing both NSCs and astrocytes (*6*) and the use of specific markers that we previously identified as being reliable to distinguish between astrocytes and NSCs across neurogenic niches (*1*) suggested that they all correspond to the same subpopulation of mature medial striatal astrocytes. **B.** Identification and projection of a score specifically identifying qNSC1(*3*) cells to other adult mouse SEZ datasets (*4*, *5*). The cells displaying highest qNSC1 scores are putative astrocytes from these other datasets (see also Methods, and new conclusions on qNSC1 in (*7*)). **C.** Metaneighbour (*8*) mapping of non-proliferating astroglia between a SEZ dataset describing the presence of qNSC1 cells (*9*) and astroglia from the telencephalon extracted from a whole brain cell atlas (*6*). Contrary to qNSC2 and aNSC1, qNSC1 are closer to protoplasmic astrocytes than to RG-like cells. **D.** Compared expression of *Cd9*, a putative marker of qNSC1 between striatal and cortical astrocytes in the adult mouse (*10*). **E,F.** Example images of lesioned pallia at 3dpl (E) or 5dpl (F). E, F are Images represent large fields of view of 3D whole-mount hemispheres illustrating localized increase in proliferation around the lesion. Scale bar: 50µm. E′-F″ are close up views of ≈10µm thickness centered on the lesions at different depths. Scale bars: 15µm. E′, F′ are at the parenchymal depth where the lesion is best visible; E″, F″ are at the ventricular surface, showing the tela choroida, which was not removed. An abundant proliferative response can be observed at 3dpl and 5dpl, specifically in the NSC layer and just below (E′, F′). At 3dpl the lesion itself is still visible whereas at 5dpl the hole has been filled. **G.** Percentage of Pcna$^{neg}$ and Pcna$^{pos}$ cells in ventricular regions located far from the lesion (control area) or close to the lesion (lesion proximity). Cells close to the lesion proliferate more, as expected with a regenerative response. **H.** Relative *timp4.3* expression in Pcna$^{pos}$ cells between cells close to the lesion (control cells) vs far from the lesion (y axis) as a function of the decile of *timp4.3* expression in control cells (x axis). Representation of the shift function as calculated in (*11*). The x-axis represents *timp4.3* expression values, measured as number of dots in smFISH, in control cells binned by deciles and the y-axis represents the difference between the average *timp4.3* expression value in cells neighboring the lesion and in control cells for each decile of their respective distribution. The circles represent the point estimates of this difference for each decile and the vertical bars represent the confidence interval. A positive difference suggests that the average *timp4.3* expression is higher in the control cells than in cells neighboring the lesion. The top 10% values of *timp4.3* expression (gray area) appear on average higher in cells close to the lesion than in cells far away from the lesion, but the difference is not statistically significant.

**Figure S5 - *nr2f1b* and *ascl1a* are expressed in opposite gradients in the dorsal pallium and this prefigures the resistance to LY treatment.** Whole-mount dorsal view of a telencephalic hemisphere (anterior left) in fish treated with DMSO or LY for 24 hours and immunostained for ZO1 (white, apical junctions) and Pcna (red, proliferation) and processed for smFISH for *nr2f1b* (cyan) and *ascl1a* (magenta). *ascl1a* and Pcna expression are enriched rostrally in the dorsal pallium in control conditions whereas *nr2f1b* expression is enriched caudally. After 24 hours of LY treatment, *ascl1a* expression is more widespread but remains enriched rostrally (this also applies to Pcna after 48 hours, see Fig.S6A middle panel), opposite to *nr2f1b*. Scale bars: 50μm.

**Figure S6: Assessment of putative regulatory mechanisms explaining the resistance to Notch inhibition. A.** Confocal images of whole individual telencephalic hemispheres from Tg(*her4*:dRFP) fish treated for 48 hours with either DMSO (left panels), LY (middle) or DMH1 (right). The brains were immunostained for ZO1 (white, apical junctions), Pcna (red, proliferation) and dRFP (magenta). The top row shows all channels, the middle row shows the detection of ZO1 and RFP, and the bottom row shows ZO1 and Pcna. While both LY and DMH1 efficiently decrease dRFP, only LY increases proliferation. Scale bars: 50 μm. **B.** Confocal images of caudal areas in the dorsal pallium of fish treated for 48 hours with either DMSO (left panel), LY alone (middle) or LY and DMH1 together (right), and immunostained for ZO1 (white, apical junctions) and Pcna (red, proliferation). Scale bars: 15μm. **C.** Quantification of the association between type of treatment and likelihood to start cycling after 48 hours of treatment. No significant difference is detected between treatments with LY alone or LY + DMH1. Reported p-value is derived from a chi square test. Violin plots are built from bootstrapped random sampling of the measured proportions to estimate the distribution. **D.** Confocal images from dorsal views of the pallium of a Tg(*her4:dRFP*) fish after treatment with either DMSO (left) or LY (right) for 48 hours and immunostained for ZO1 (white, apical junctions) and dRFP (magenta, to reveal activity of the 3.4kb *her4* promoter, and processed for smFISH for *nr2f1b* (cyan). There is no correlation between *nr2f1b* expression and dRFP intensity. Scale bars: 5μm.

### Legends for Supplementary Tables

**Table S1. Marker genes used to define broad cell identities from the full scRNAseq dataset.**

**Table S2. Differentially expressed genes (DEGs) between DMSO and LY conditions in each cluster of quiescent NSCs/IPCs.** Positive LogFC when expression is higher in the DMSO dataset. pctDMSO, pctLY: proportion of cells expressing the gene in the DMSO and LY datasets, respectively.

**Table S3. Differentially expressed genes (DEGs) between DMSO and LY conditions in each cluster of proliferating NSCs/IPCs.** Positive LogFC when expression is higher in the DMSO dataset. pctDMSO, pctLY: proportion of cells expressing the gene in the DMSO and LY datasets, respectively.

**Table S4. Antibodies used in this study**

**Table S5. RNAscope probes used in this study**

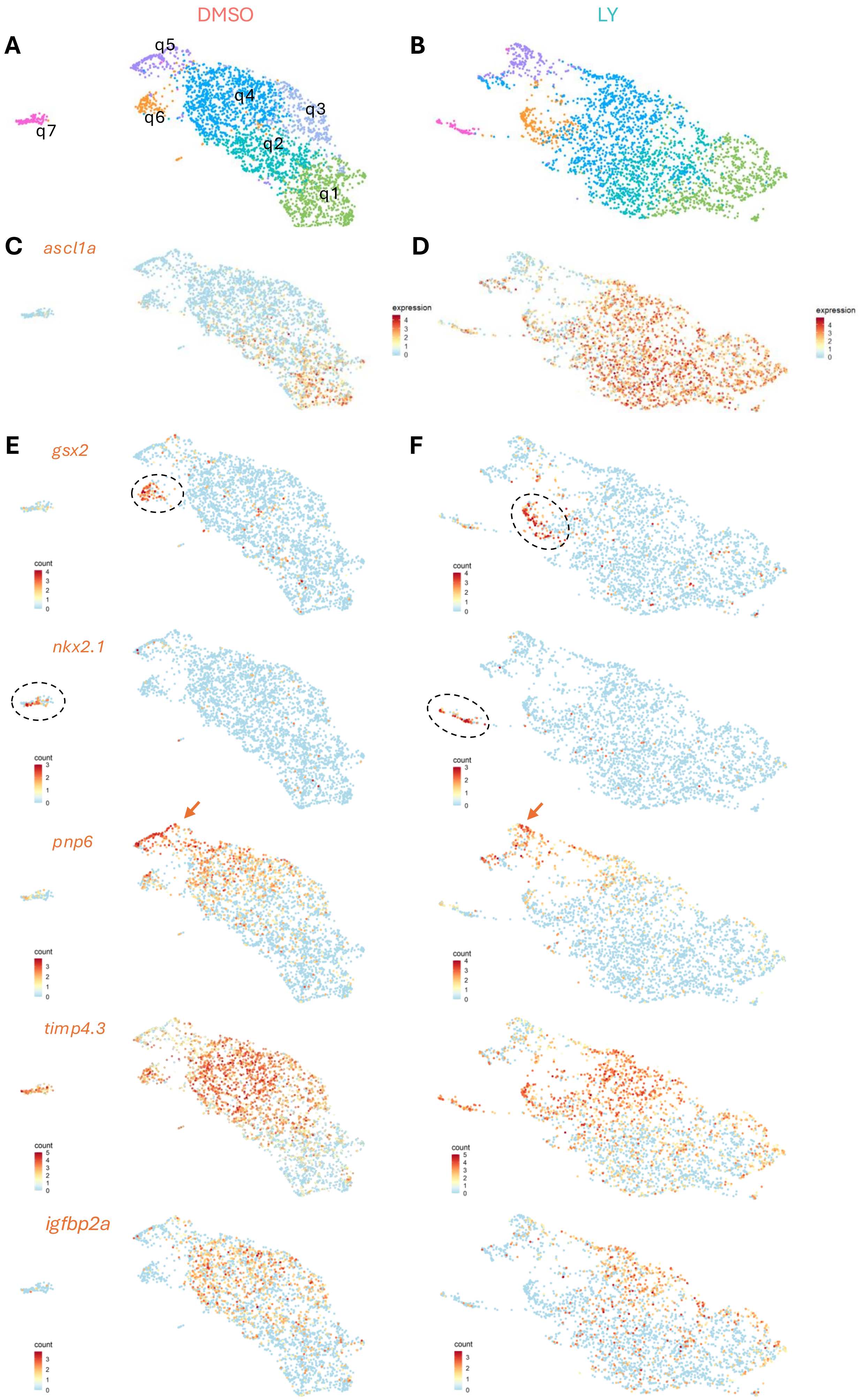


Morizet et al., Figure S1

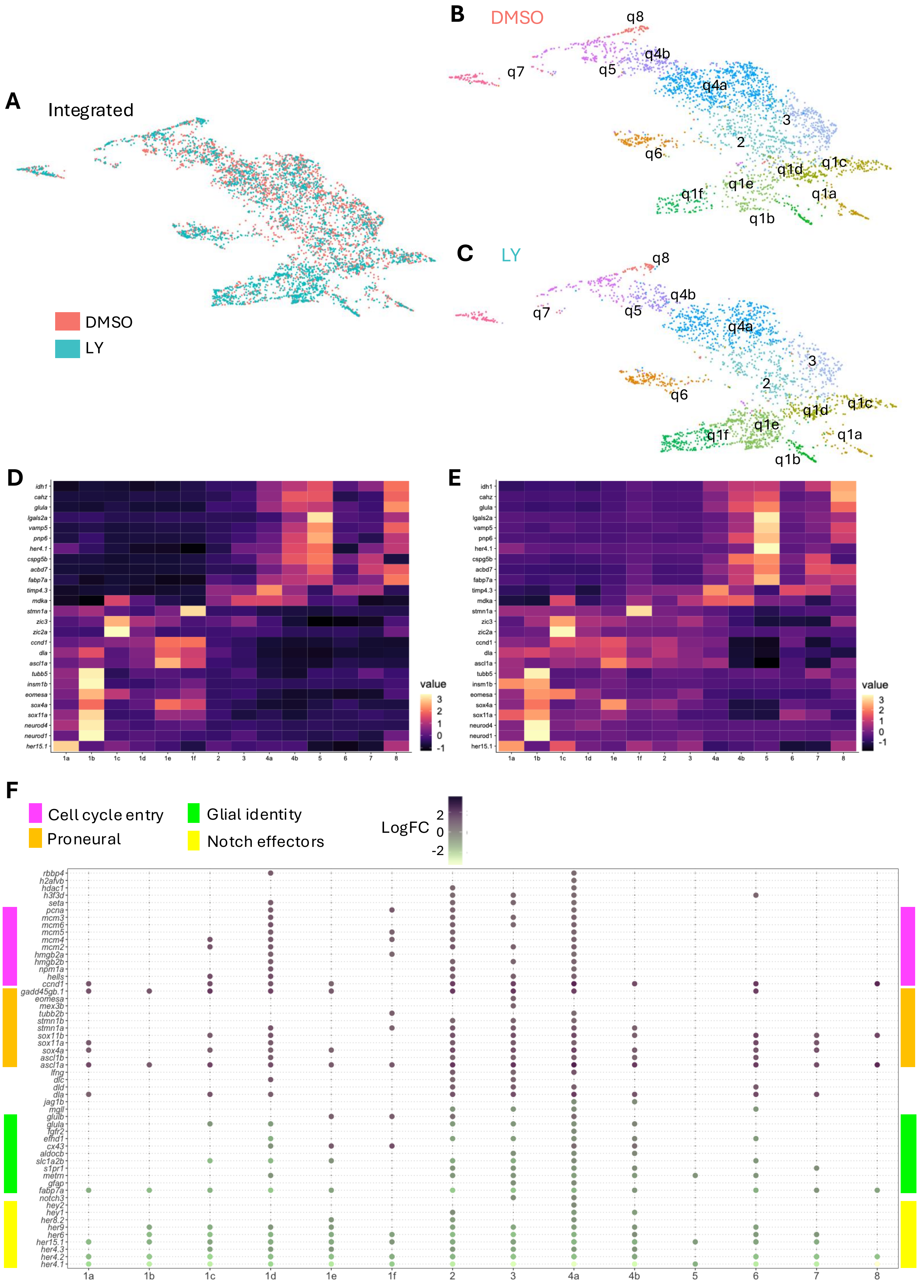


Morizet et al., Figure S2

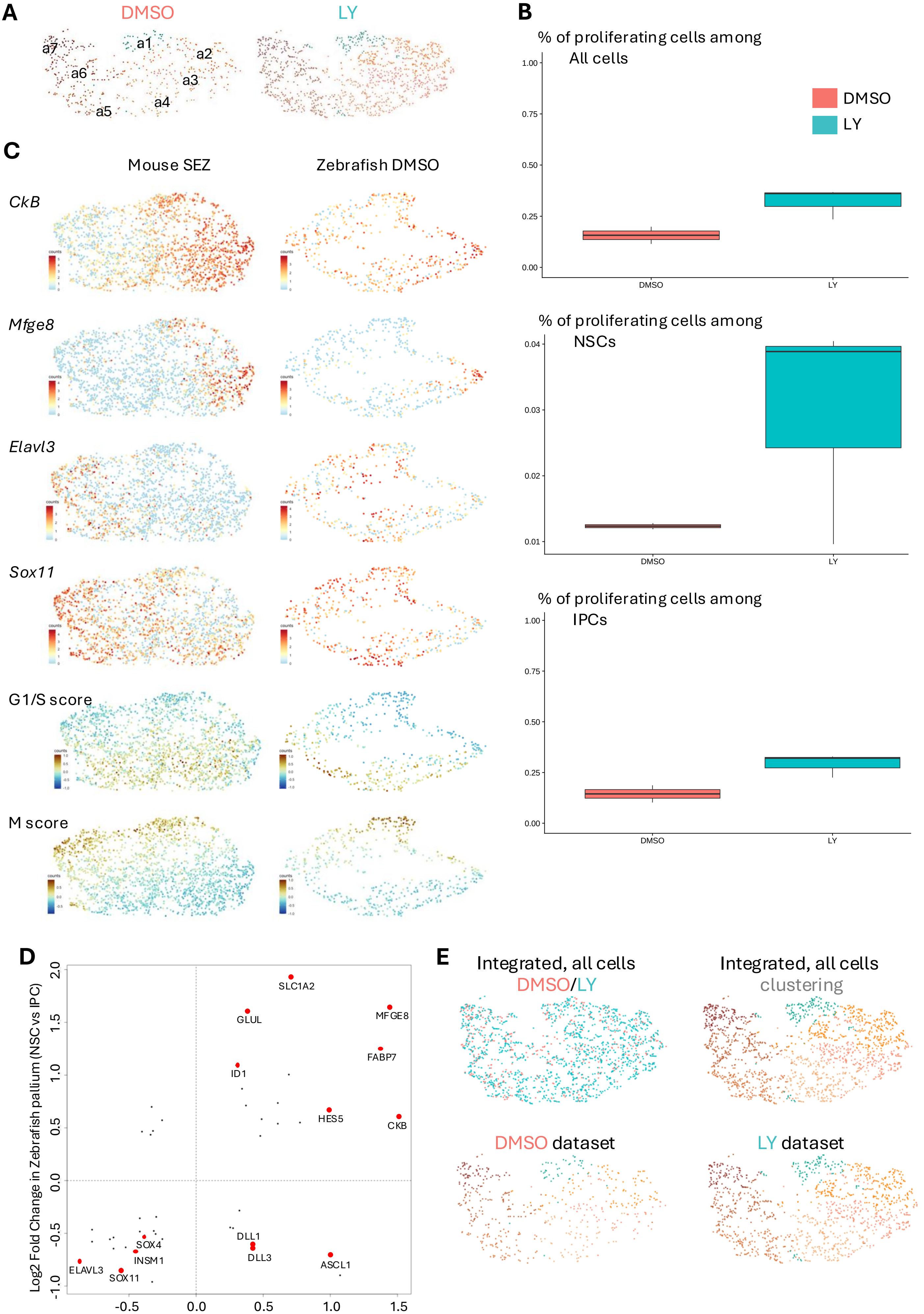


Morizet et al., Figure S3

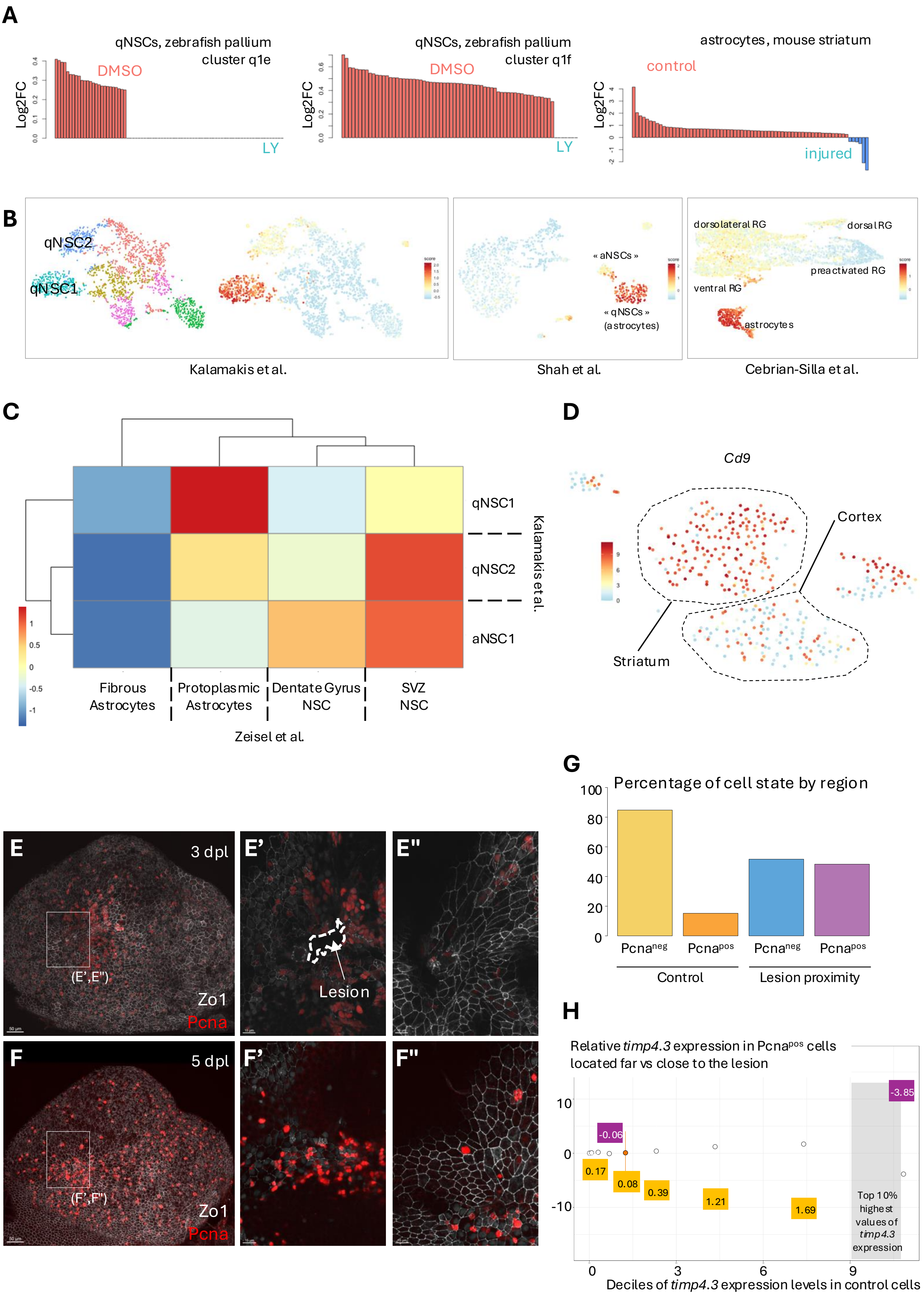


Morizet et al., Figure S4

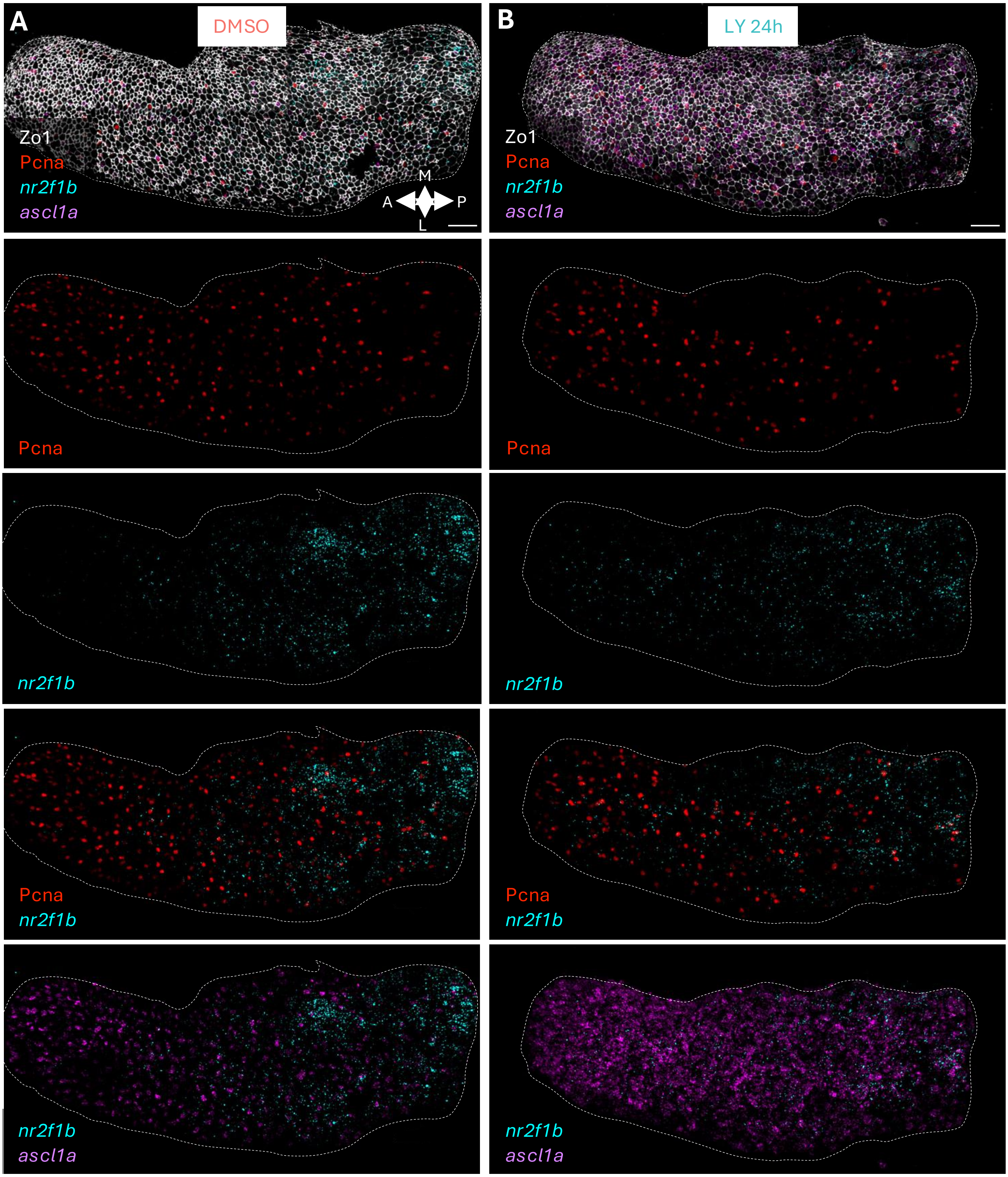


Morizet et al., Figure S5

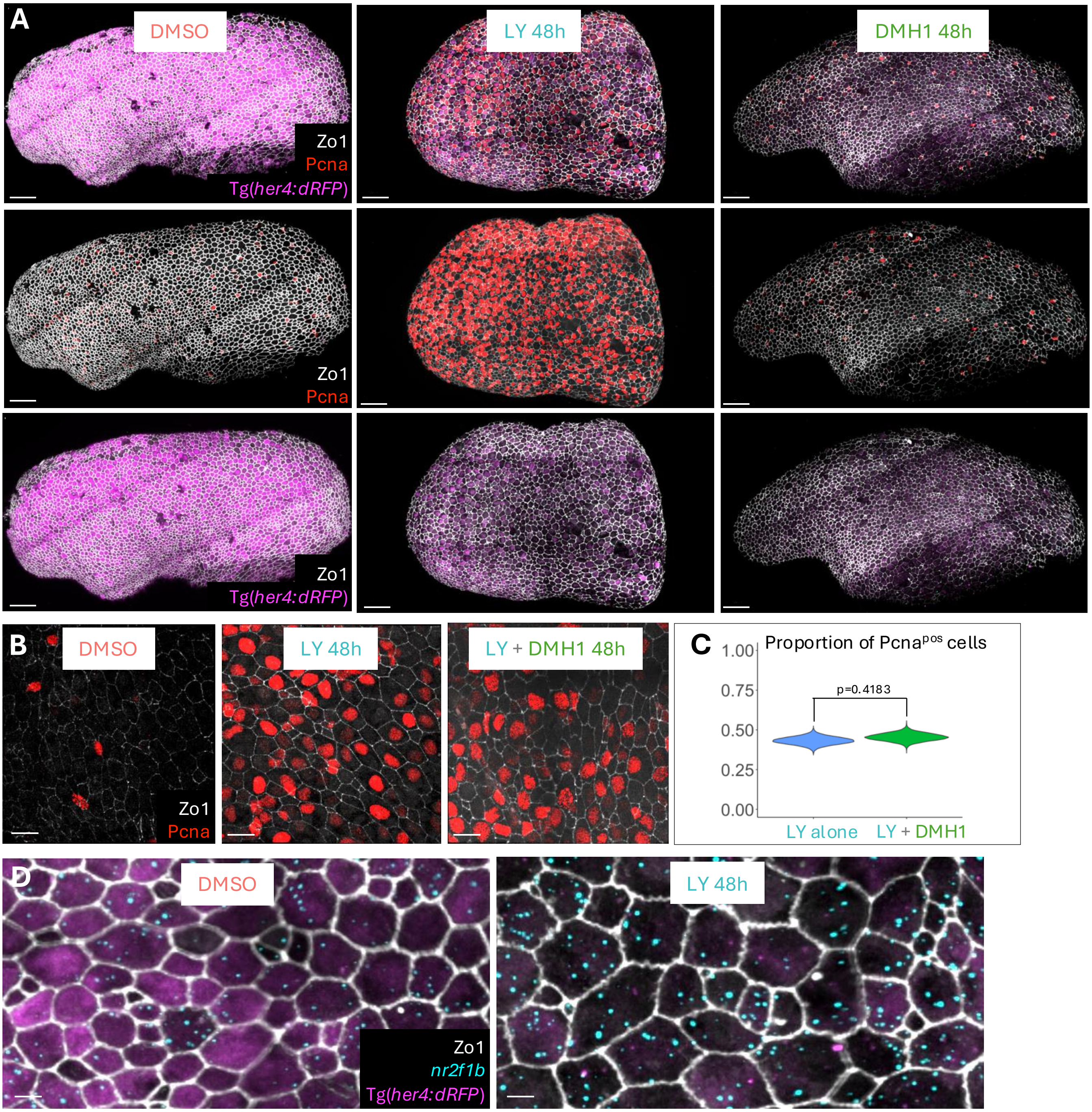


Morizet et al., Figure S6

| Broad cluster |
| --- |
| Radial Glia |
| Neurons (different types and at different levels of maturation) |
| Proliferating cells (mostly aNSCs, IPCs, and a few proliferating oligodendrocytes that could be remove |
| Oligodendrocytic lineage |
| Macrophage lineage |

| Markers |
|---|
| fabp7a, glula, cx43, gfap, s100b |
| tubb5, elavl3, neurod1, neurod6a, slc32a1, gad1b, slc16a6b, slc16a7a |
| mcm2, pcna, cdk1, |
| aplnra, aplnrb, sox10, olig2, mbpb, mpz, plp1a |
| cd74a, mfap4, mpeg1.1, irf8 |

Table S2

q1a_DMSOvsLY

| | p_val | avg_logFC | pctDMSO | pctLY | p_val_adj | gene_name |
|---|---|---|---|---|---|---|
| ENSDARG00000056732 | 2,65E-14 | 2,20866032 | 0,844 | 0,26 | 4,56E-10 | her4.1 |
| ENSDARG00000094426 | 3,33E-16 | 1,58405757 | 0,948 | 0,62 | 5,73E-12 | her4.2 |
| ENSDARG00000054562 | 1,81E-09 | 1,551653245 | 0,792 | 0,32 | 3,11E-05 | her15.1 |
| ENSDARG00000054560 | 8,97E-07 | 1,343212916 | 0,667 | 0,28 | 0,015435331 | her15.1 |
| ENSDARG00000007697 | 4,69E-14 | 1,287309917 | 0,99 | 0,94 | 8,06E-10 | fabp7a |
| ENSDARG00000056729 | 3,61E-08 | 1,221760965 | 0,844 | 0,72 | 0,000620898 | her4.2 |
| ENSDARG00000010791 | 4,17E-07 | -0,61187562 | 0,719 | 0,98 | 0,00716786 | dla |
| ENSDARG00000077811 | 1,82E-06 | -0,83470099 | 0,51 | 0,84 | 0,031326219 | sox11a |
| ENSDARG00000004588 | 7,90E-08 | -0,83815863 | 0,625 | 0,9 | 0,001359752 | sox4a |
| ENSDARG00000101637 | 3,81E-10 | -0,94773491 | 0,594 | 0,96 | 6,56E-06 | ccnd1 |
| ENSDARG00000016725 | 3,38E-08 | -1,04259164 | 0,292 | 0,76 | 0,000580819 | gadd45gb.1 |
| ENSDARG00000038386 | 2,17E-12 | -1,33318582 | 0,448 | 0,86 | 3,74E-08 | ascl1a |

Table S2

q1b_DMSOvsLY

| | p_val | avg_logFC | pctDMSO | pctLY | p_val_adj | gene_name |
|---|---|---|---|---|---|---|
| ENSDARG00000056732 | 3,44E-21 | 2,59243788 | 0,857 | 0,121 | 5,91E-17 | her4.1 |
| ENSDARG00000094426 | 6,71E-19 | 1,85807425 | 0,984 | 0,484 | 1,15E-14 | her4.2 |
| ENSDARG00000009822 | 5,78E-16 | 1,75555861 | 0,889 | 0,275 | 9,94E-12 | her4.4 |
| ENSDARG00000006514 | 1,52E-08 | 1,58301981 | 0,54 | 0,055 | 0,000261661 | her6 |
| ENSDARG00000007697 | 1,11E-12 | 1,5714622 | 1 | 0,725 | 1,90E-08 | fabp7a |
| ENSDARG00000056729 | 5,00E-15 | 1,44878792 | 0,921 | 0,593 | 8,61E-11 | her4.2 |
| ENSDARG00000054562 | 5,27E-08 | 1,34545857 | 0,619 | 0,154 | 0,000905816 | her15.1 |
| ENSDARG00000056438 | 6,12E-09 | 1,14666422 | 0,317 | 0 | 0,000105346 | her9 |
| ENSDARG00000038386 | 8,07E-07 | -0,85061642 | 0,619 | 0,791 | 0,013889289 | ascl1a |
| ENSDARG00000016725 | 1,98E-07 | -0,97780537 | 0,556 | 0,846 | 0,003399556 | gadd45gb.1 |

Table S2

q1c_DMSOvsLY

| | p_val | avg_logFC | pctDMSO | pctLY | p_val_adj | gene_name |
|---|---|---|---|---|---|---|
| ENSDARG00000056732 | 1,90E-25 | 2,616173225 | 0,669 | 0,043 | 3,28E-21 | her4.1 |
| ENSDARG00000094426 | 9,31E-27 | 2,093084337 | 0,847 | 0,198 | 1,60E-22 | her4.2 |
| ENSDARG00000056729 | 2,21E-19 | 2,047191285 | 0,702 | 0,164 | 3,81E-15 | her4.2 |
| ENSDARG00000009822 | 2,40E-20 | 1,98561948 | 0,645 | 0,069 | 4,12E-16 | her4.4 |
| ENSDARG00000006514 | 1,19E-17 | 1,678786415 | 0,597 | 0,078 | 2,05E-13 | her6 |
| ENSDARG00000102453 | 4,32E-27 | 1,603071224 | 0,944 | 0,5 | 7,43E-23 | slc1a2b |
| ENSDARG00000007697 | 9,50E-31 | 1,4044397 | 0,984 | 0,905 | 1,63E-26 | fabp7a |
| ENSDARG00000056438 | 4,04E-10 | 1,261398695 | 0,452 | 0,069 | 6,95E-06 | her9 |
| ENSDARG00000070770 | 8,56E-09 | 1,06974791 | 0,339 | 0,034 | 0,000147214 | her4.3 |
| ENSDARG00000054562 | 7,72E-10 | 1,045577685 | 0,637 | 0,284 | 1,33E-05 | her15.1 |
| ENSDARG00000095715 | 1,71E-06 | 0,767346356 | 0,653 | 0,319 | 0,029504421 | BX465834.1 |
| ENSDARG00000099776 | 3,55E-07 | 0,720097386 | 0,984 | 0,948 | 0,006110875 | glula |
| ENSDARG00000007275 | 6,24E-09 | 0,622539162 | 0,968 | 0,888 | 0,000107307 | si:ch211-251b21.1 |
| ENSDARG00000004588 | 2,10E-06 | -0,75642284 | 0,468 | 0,741 | 0,036045276 | sox4a |
| ENSDARG00000076532 | 2,61E-07 | -0,796822046 | 0,395 | 0,638 | 0,004493552 | si:ch211-222l21.1 |
| ENSDARG00000095743 | 2,66E-06 | -0,81758439 | 0,306 | 0,56 | 0,045717597 | sox11b |
| ENSDARG00000102798 | 3,13E-07 | -0,954965861 | 0,048 | 0,259 | 0,00538709 | mcm2 |
| ENSDARG00000010791 | 4,84E-15 | -0,962656212 | 0,581 | 0,905 | 8,33E-11 | dla |
| ENSDARG00000040041 | 7,80E-14 | -1,015490612 | 0,008 | 0,267 | 1,34E-09 | mcm4 |
| ENSDARG00000101637 | 1,94E-21 | -1,105263873 | 0,653 | 0,974 | 3,34E-17 | ccnd1 |
| ENSDARG00000057738 | 3,07E-09 | -1,200582822 | 0,105 | 0,345 | 5,27E-05 | hells |
| ENSDARG00000038386 | 3,87E-18 | -1,435359211 | 0,355 | 0,784 | 6,67E-14 | ascl1a |
| ENSDARG00000016725 | 8,86E-14 | -1,743460863 | 0,089 | 0,5 | 1,52E-09 | gadd45gb.1 |

Table S2

q1d_DMSOvsLY

| | p_val | avg_logFC | pctDMSO | pctLY | p_val_adj | gene_name |
|---|---|---|---|---|---|---|
| ENSDARG00000056732 | 6,02E-29 | 2,128234219 | 0,695 | 0,064 | 1,04E-24 | her4.1 |
| ENSDARG00000007697 | 8,93E-47 | 1,797737875 | 0,987 | 0,829 | 1,54E-42 | fabp7a |
| ENSDARG00000094426 | 7,46E-22 | 1,647682946 | 0,786 | 0,35 | 1,28E-17 | her4.2 |
| ENSDARG00000102453 | 6,02E-35 | 1,533812998 | 0,948 | 0,436 | 1,04E-30 | slc1a2b |
| ENSDARG00000056729 | 4,56E-16 | 1,463741023 | 0,682 | 0,271 | 7,84E-12 | her4.2 |
| ENSDARG00000006514 | 3,82E-14 | 1,400749755 | 0,435 | 0,043 | 6,57E-10 | her6 |
| ENSDARG00000009822 | 2,46E-14 | 1,343797265 | 0,61 | 0,164 | 4,23E-10 | her4.4 |
| ENSDARG00000043446 | 1,04E-18 | 1,323008956 | 0,721 | 0,207 | 1,79E-14 | efhd1 |
| ENSDARG00000054562 | 2,05E-15 | 1,269393727 | 0,604 | 0,157 | 3,52E-11 | her15.1 |
| ENSDARG00000007275 | 3,75E-26 | 1,156042022 | 0,935 | 0,593 | 6,46E-22 | si:ch211-251b21.1 |
| ENSDARG00000054560 | 2,21E-11 | 1,106034794 | 0,513 | 0,143 | 3,81E-07 | her15.1 |
| ENSDARG00000099776 | 3,72E-16 | 0,898278807 | 0,955 | 0,9 | 6,39E-12 | glula |
| ENSDARG00000070770 | 1,92E-06 | 0,860329251 | 0,266 | 0,043 | 0,033103537 | her4.3 |
| ENSDARG00000056438 | 5,83E-08 | 0,72510347 | 0,195 | 0,007 | 0,00100349 | her9 |
| ENSDARG00000030367 | 8,42E-08 | 0,67267842 | 0,214 | 0,014 | 0,001447772 | metrn |
| ENSDARG00000019856 | 3,87E-07 | 0,66780703 | 0,877 | 0,75 | 0,006654819 | atp1a1b |
| ENSDARG00000036036 | 3,56E-15 | 0,664206135 | 0,994 | 0,829 | 6,13E-11 | mdka |
| ENSDARG00000041799 | 7,73E-07 | 0,650580849 | 0,643 | 0,3 | 0,013304611 | cx43 |
| ENSDARG00000010008 | 8,14E-07 | 0,598426609 | 0,773 | 0,507 | 0,014012565 | vim |
| ENSDARG00000036080 | 5,69E-09 | 0,597866562 | 0,903 | 0,693 | 9,78E-05 | cd81a |
| ENSDARG00000037870 | 1,17E-07 | 0,4690808 | 0,955 | 0,871 | 0,002014518 | actb2 |
| ENSDARG00000063912 | 6,50E-08 | 0,368945209 | 1 | 0,943 | 0,001118451 | mt-co3 |
| ENSDARG00000002344 | 7,44E-07 | 0,344516442 | 0,994 | 0,864 | 0,012796556 | tubb4b |
| ENSDARG00000113103 | 4,04E-08 | -0,488963018 | 0 | 0,121 | 0,000695616 | BX530077.1 |
| ENSDARG00000040184 | 5,07E-07 | -0,499126615 | 0,468 | 0,736 | 0,008722281 | syncrip |
| ENSDARG00000101766 | 4,27E-11 | -0,522167405 | 0,929 | 0,964 | 7,35E-07 | ptmab |
| ENSDARG00000029722 | 2,15E-07 | -0,636460551 | 0,481 | 0,671 | 0,003697939 | hmgb2a |
| ENSDARG00000076532 | 5,76E-11 | -0,663094271 | 0,591 | 0,85 | 9,91E-07 | si:ch211-222l21.1 |
| ENSDARG00000104353 | 1,79E-06 | -0,699916806 | 0,195 | 0,393 | 0,030875987 | nop58 |
| ENSDARG00000019507 | 1,25E-06 | -0,72256909 | 0,026 | 0,207 | 0,021555112 | mcm5 |
| ENSDARG00000010791 | 4,84E-13 | -0,72436822 | 0,747 | 0,936 | 8,33E-09 | dla |
| ENSDARG00000004588 | 1,35E-08 | -0,730675863 | 0,604 | 0,821 | 0,000231905 | sox4a |
| ENSDARG00000101707 | 6,38E-09 | -0,745952238 | 0,052 | 0,257 | 0,000109786 | si:ch211-156b7.4 |
| ENSDARG00000031495 | 2,53E-09 | -0,746047837 | 0,461 | 0,664 | 4,35E-05 | seta |
| ENSDARG00000023299 | 3,82E-07 | -0,75369467 | 0,221 | 0,429 | 0,00657704 | snu13b |
| ENSDARG00000078155 | 1,25E-07 | -0,766098524 | 0,058 | 0,286 | 0,002149703 | inavaa |
| ENSDARG00000053990 | 8,02E-13 | -0,778134947 | 0,649 | 0,864 | 1,38E-08 | hmgb2b |
| ENSDARG00000029058 | 2,12E-07 | -0,781101049 | 0,136 | 0,357 | 0,003639635 | rbbp4 |
| ENSDARG00000101628 | 3,98E-07 | -0,793746751 | 0,071 | 0,314 | 0,006855251 | ascl1b |
| ENSDARG00000077811 | 5,76E-09 | -0,804353676 | 0,273 | 0,557 | 9,91E-05 | sox11a |
| ENSDARG00000054155 | 5,52E-09 | -0,8139678 | 0,13 | 0,35 | 9,50E-05 | pcna |
| ENSDARG00000102798 | 2,68E-08 | -0,840135814 | 0,136 | 0,386 | 0,000460815 | mcm2 |
| ENSDARG00000024204 | 2,77E-07 | -0,868159947 | 0,058 | 0,271 | 0,004770882 | mcm3 |
| ENSDARG00000040041 | 5,68E-10 | -0,902930239 | 0,039 | 0,286 | 9,77E-06 | mcm4 |
| ENSDARG00000002336 | 4,08E-09 | -0,911632635 | 0,026 | 0,271 | 7,02E-05 | dlc |
| ENSDARG00000101637 | 1,13E-22 | -0,918010963 | 0,747 | 0,971 | 1,95E-18 | ccnd1 |
| ENSDARG00000014329 | 3,16E-09 | -0,934005537 | 0,156 | 0,457 | 5,43E-05 | npm1a |
| ENSDARG00000004169 | 4,54E-11 | -0,96567749 | 0,149 | 0,436 | 7,82E-07 | stmn1a |
| ENSDARG00000095743 | 4,66E-12 | -0,965810717 | 0,403 | 0,729 | 8,02E-08 | sox11b |
| ENSDARG00000057683 | 2,86E-10 | -1,01170297 | 0,065 | 0,314 | 4,92E-06 | mcm6 |
| ENSDARG00000038386 | 2,93E-19 | -1,133806877 | 0,461 | 0,871 | 5,04E-15 | ascl1a |
| ENSDARG00000057738 | 2,82E-16 | -1,175427902 | 0,078 | 0,45 | 4,85E-12 | hells |
| ENSDARG00000016725 | 1,06E-23 | -1,464914964 | 0,227 | 0,764 | 1,83E-19 | gadd45gb.1 |

Table S2
q1e_DMSOvsLY

| | p_val | avg_logFC | pctDMSO | pctLY | p_val_adj | gene_name |
|---|---|---|---|---|---|---|
| ENSDARG00000056732 | 5,17E-63 | 2,482765666 | 0,636 | 0,04 | 8,89E-59 | her4.1 |
| ENSDARG00000094426 | 5,07E-55 | 1,673435392 | 0,87 | 0,39 | 8,73E-51 | her4.2 |
| ENSDARG00000056729 | 3,18E-45 | 1,595936405 | 0,799 | 0,37 | 5,47E-41 | her4.2 |
| ENSDARG00000009822 | 3,45E-34 | 1,449029983 | 0,707 | 0,21 | 5,93E-30 | her4.4 |
| ENSDARG00000006514 | 3,76E-26 | 1,236942791 | 0,446 | 0,07 | 6,47E-22 | her6 |
| ENSDARG00000054562 | 2,70E-22 | 1,216097965 | 0,484 | 0,12 | 4,65E-18 | her15.1 |
| ENSDARG00000054560 | 1,75E-16 | 1,061469688 | 0,457 | 0,15 | 3,01E-12 | her15.1 |
| ENSDARG00000007697 | 2,85E-37 | 1,034936264 | 0,973 | 0,89 | 4,91E-33 | fabp7a |
| ENSDARG00000056438 | 1,67E-11 | 0,806244685 | 0,174 | 0,02 | 2,87E-07 | her9 |
| ENSDARG00000069675 | 4,36E-09 | 0,736793696 | 0,212 | 0,05 | 7,49E-05 | her8.2 |
| ENSDARG00000070770 | 1,59E-09 | 0,726391095 | 0,321 | 0,1 | 2,73E-05 | her4.3 |
| ENSDARG00000102453 | 1,80E-11 | 0,607803738 | 0,902 | 0,73 | 3,10E-07 | slc1a2b |
| ENSDARG00000101794 | 3,07E-08 | 0,501014125 | 0,56 | 0,39 | 0,00052816 | atp6v0e1 |
| ENSDARG00000053365 | 1,74E-07 | 0,409042689 | 0,69 | 0,52 | 0,002992355 | rpl31 |
| ENSDARG00000036629 | 3,22E-12 | 0,402727201 | 0,946 | 0,89 | 5,53E-08 | rps14 |
| ENSDARG00000015128 | 2,87E-12 | 0,393934435 | 0,951 | 0,86 | 4,93E-08 | rpl27 |
| ENSDARG00000011201 | 2,01E-15 | 0,392807391 | 0,967 | 0,95 | 3,47E-11 | rplp2l |
| ENSDARG00000009212 | 2,90E-06 | 0,350432607 | 0,766 | 0,65 | 0,04988695 | ppiaa |
| ENSDARG00000036875 | 2,82E-12 | 0,344632347 | 0,978 | 0,96 | 4,86E-08 | rps12 |
| ENSDARG00000057556 | 1,13E-07 | 0,330468776 | 0,929 | 0,85 | 0,001936093 | rpl17 |
| ENSDARG00000099104 | 1,05E-06 | 0,329142375 | 0,962 | 0,86 | 0,018070028 | rpl24 |
| ENSDARG00000041232 | 2,69E-09 | 0,324898244 | 0,957 | 0,91 | 4,63E-05 | rps29 |
| ENSDARG00000043509 | 1,51E-12 | 0,322058639 | 0,989 | 0,98 | 2,60E-08 | rpl11 |
| ENSDARG00000021838 | 7,20E-07 | 0,299434946 | 0,848 | 0,76 | 0,012380631 | rps23 |
| ENSDARG00000051783 | 2,37E-08 | 0,297989121 | 0,995 | 0,96 | 0,000407974 | rplp0 |
| ENSDARG00000006413 | 2,00E-08 | 0,297448416 | 0,973 | 0,89 | 0,000343306 | rpl38 |
| ENSDARG00000099380 | 3,03E-07 | 0,294159441 | 0,951 | 0,91 | 0,005210071 | rpl13 |
| ENSDARG00000034291 | 2,69E-11 | 0,286272706 | 0,989 | 0,99 | 4,63E-07 | rpl37 |
| ENSDARG00000021864 | 5,47E-11 | 0,282076853 | 1 | 0,99 | 9,41E-07 | rplp1 |
| ENSDARG00000046157 | 3,88E-08 | 0,281293437 | 0,913 | 0,88 | 0,000667653 | RPS17 |
| ENSDARG00000018334 | 2,59E-07 | 0,277191269 | 0,978 | 0,94 | 0,004453185 | rpl35 |
| ENSDARG00000070849 | 4,04E-08 | 0,27040958 | 0,978 | 0,93 | 0,000694362 | rps15 |
| ENSDARG00000088030 | 3,76E-07 | 0,269816669 | 0,967 | 0,92 | 0,006470724 | rpl35a |
| ENSDARG00000042566 | 9,18E-10 | 0,268262725 | 0,995 | 0,99 | 1,58E-05 | rps7 |
| ENSDARG00000037350 | 2,31E-06 | 0,26676671 | 0,957 | 0,89 | 0,039723859 | rpl9 |
| ENSDARG00000055996 | 2,58E-06 | 0,264904821 | 0,973 | 0,96 | 0,044305989 | rps8a |
| ENSDARG00000019181 | 9,48E-08 | 0,26315578 | 0,984 | 0,95 | 0,001631235 | rpsa |
| ENSDARG00000058105 | 3,78E-07 | 0,256306272 | 0,973 | 0,93 | 0,006504557 | rpl36a |
| ENSDARG00000009285 | 3,47E-07 | 0,253039954 | 0,978 | 0,96 | 0,005974312 | rpl15 |
| ENSDARG00000058451 | 2,70E-06 | 0,250162015 | 0,957 | 0,88 | 0,046381331 | rpl6 |
| ENSDARG00000101766 | 6,32E-07 | -0,256299442 | 0,962 | 0,96 | 0,010872372 | ptmab |
| ENSDARG00000052082 | 3,74E-07 | -0,337714407 | 0,712 | 0,82 | 0,006431996 | gabarapb |
| ENSDARG00000025147 | 1,06E-07 | -0,375589248 | 0,761 | 0,83 | 0,001817223 | cd63 |
| ENSDARG00000038068 | 2,50E-09 | -0,427513691 | 0,63 | 0,83 | 4,30E-05 | ddx5 |
| ENSDARG00000043608 | 1,84E-07 | -0,473749974 | 0,435 | 0,59 | 0,003167663 | eif4ebp1 |
| ENSDARG00000101637 | 2,06E-14 | -0,484754263 | 0,87 | 0,97 | 3,55E-10 | ccnd1 |
| ENSDARG00000015349 | 1,48E-06 | -0,603398408 | 0,174 | 0,31 | 0,025532645 | mfge8a |
| ENSDARG00000041623 | 2,66E-11 | -0,620364258 | 0,777 | 0,87 | 4,58E-07 | mt2 |
| ENSDARG00000004588 | 9,54E-20 | -0,637480363 | 0,853 | 0,95 | 1,64E-15 | sox4a |
| ENSDARG00000100795 | 9,31E-07 | -0,655339912 | 0,19 | 0,34 | 0,016019071 | timp4.3 |
| ENSDARG00000100003 | 6,22E-19 | -0,684357316 | 0,679 | 0,88 | 1,07E-14 | glulb |
| ENSDARG00000038386 | 4,82E-23 | -0,8006631 | 0,734 | 0,91 | 8,29E-19 | ascl1a |
| ENSDARG00000020239 | 1,64E-07 | -0,810119625 | 0,071 | 0,23 | 0,002813837 | lpin1 |
| ENSDARG00000016725 | 6,82E-18 | -0,82994192 | 0,467 | 0,77 | 1,17E-13 | gadd45gb.1 |
| ENSDARG00000041799 | 5,48E-19 | -0,831832234 | 0,674 | 0,81 | 9,43E-15 | cx43 |

Table S2
q1f_DMSOvsLY

| | p_val | avg_logFC | pctDMSO | pctLY | p_val_adj | gene_name |
|---|---|---|---|---|---|---|
| ENSDARG00000056732 | 9,02E-30 | 1,885981917 | 0,636 | 0,09 | 1,55E-25 | her4.1 |
| ENSDARG00000009822 | 2,17E-14 | 1,344166538 | 0,589 | 0,29 | 3,74E-10 | her4.4 |
| ENSDARG00000094426 | 3,91E-20 | 1,287972068 | 0,785 | 0,41 | 6,73E-16 | her4.2 |
| ENSDARG00000006514 | 1,74E-08 | 1,010110601 | 0,383 | 0,15 | 0,000299537 | her6 |
| ENSDARG00000056729 | 2,11E-11 | 1,002980997 | 0,682 | 0,34 | 3,63E-07 | her4.2 |
| ENSDARG00000054562 | 1,12E-07 | 0,74126455 | 0,533 | 0,28 | 0,001925464 | her15.1 |
| ENSDARG00000011201 | 1,81E-21 | 0,701807056 | 1 | 0,93 | 3,12E-17 | rplp2l |
| ENSDARG00000006413 | 9,69E-18 | 0,672195769 | 1 | 0,91 | 1,67E-13 | rpl38 |
| ENSDARG00000039347 | 5,10E-15 | 0,593464775 | 0,972 | 0,88 | 8,78E-11 | rps24 |
| ENSDARG00000077717 | 1,65E-14 | 0,589956708 | 0,972 | 0,93 | 2,83E-10 | rpl29 |
| ENSDARG00000104011 | 9,96E-08 | 0,580306396 | 0,748 | 0,6 | 0,001714016 | rps17 |
| ENSDARG00000041435 | 4,10E-09 | 0,577457059 | 0,869 | 0,73 | 7,06E-05 | uba52 |
| ENSDARG00000014690 | 5,95E-15 | 0,576009842 | 0,991 | 0,93 | 1,02E-10 | rps4x |
| ENSDARG00000030408 | 1,19E-12 | 0,574260161 | 0,953 | 0,8 | 2,04E-08 | rps26l |
| ENSDARG00000041232 | 2,24E-15 | 0,573245124 | 0,981 | 0,88 | 3,85E-11 | rps29 |
| ENSDARG00000021838 | 1,97E-10 | 0,546692169 | 0,935 | 0,81 | 3,39E-06 | rps23 |
| ENSDARG00000025850 | 9,31E-13 | 0,54132966 | 0,991 | 0,94 | 1,60E-08 | rps21 |
| ENSDARG00000010516 | 6,64E-11 | 0,530351312 | 0,981 | 0,9 | 1,14E-06 | rpl21 |
| ENSDARG00000058451 | 3,52E-13 | 0,527811405 | 0,953 | 0,91 | 6,05E-09 | rpl6 |
| ENSDARG00000042905 | 7,66E-13 | 0,525654876 | 1 | 0,91 | 1,32E-08 | rpl10a |
| ENSDARG00000058105 | 4,32E-12 | 0,524707751 | 0,991 | 0,92 | 7,44E-08 | rpl36a |
| ENSDARG00000088030 | 3,55E-12 | 0,509633738 | 1 | 0,92 | 6,11E-08 | rpl35a |
| ENSDARG00000036875 | 2,15E-15 | 0,507901974 | 0,981 | 0,98 | 3,70E-11 | rps12 |
| ENSDARG00000005791 | 1,49E-11 | 0,506657192 | 0,991 | 0,91 | 2,56E-07 | rpl28 |
| ENSDARG00000053058 | 8,27E-12 | 0,500441489 | 0,972 | 0,9 | 1,42E-07 | rps11 |
| ENSDARG00000053457 | 3,99E-12 | 0,49896786 | 0,991 | 0,96 | 6,86E-08 | rpl23 |
| ENSDARG00000041619 | 3,24E-12 | 0,498473804 | 0,991 | 0,86 | 5,57E-08 | rack1 |
| ENSDARG00000099380 | 2,16E-13 | 0,497087287 | 0,991 | 0,91 | 3,72E-09 | rpl13 |
| ENSDARG00000042566 | 4,96E-14 | 0,495940967 | 1 | 0,98 | 8,54E-10 | rps7 |
| ENSDARG00000010160 | 2,83E-14 | 0,495591878 | 0,991 | 0,97 | 4,86E-10 | rps15a |
| ENSDARG00000045487 | 1,87E-12 | 0,49238986 | 1 | 0,97 | 3,21E-08 | rps16 |
| ENSDARG00000051783 | 3,28E-09 | 0,488862272 | 0,991 | 0,96 | 5,64E-05 | rplp0 |
| ENSDARG00000011405 | 1,29E-12 | 0,480335242 | 0,981 | 0,98 | 2,22E-08 | rps9 |
| ENSDARG00000035860 | 3,69E-15 | 0,473571991 | 1 | 1 | 6,35E-11 | rps28 |
| ENSDARG00000023298 | 2,09E-11 | 0,469370507 | 0,991 | 0,97 | 3,60E-07 | rps27.1 |
| ENSDARG00000103007 | 1,58E-09 | 0,468403814 | 0,963 | 0,8 | 2,71E-05 | rps3 |
| ENSDARG00000057556 | 4,96E-09 | 0,466331002 | 0,953 | 0,89 | 8,54E-05 | rpl17 |
| ENSDARG00000035692 | 3,68E-09 | 0,465345531 | 0,991 | 0,91 | 6,33E-05 | rps3a |
| ENSDARG00000055996 | 9,74E-11 | 0,463627105 | 0,991 | 0,93 | 1,68E-06 | rps8a |
| ENSDARG00000043509 | 3,22E-11 | 0,463590275 | 1 | 0,98 | 5,53E-07 | rpl11 |
| ENSDARG00000034291 | 4,97E-14 | 0,462499982 | 1 | 0,99 | 8,55E-10 | rpl37 |
| ENSDARG00000021864 | 4,56E-13 | 0,461981265 | 1 | 0,99 | 7,85E-09 | rplp1 |
| ENSDARG00000015128 | 2,65E-09 | 0,459463047 | 0,963 | 0,89 | 4,56E-05 | rpl27 |
| ENSDARG00000036316 | 1,03E-13 | 0,458928085 | 1 | 0,98 | 1,78E-09 | rpl39 |
| ENSDARG00000018334 | 6,57E-11 | 0,453589207 | 0,991 | 0,94 | 1,13E-06 | rpl35 |
| ENSDARG00000014867 | 5,06E-11 | 0,452067178 | 0,991 | 0,97 | 8,70E-07 | rpl8 |
| ENSDARG00000043453 | 4,18E-10 | 0,450757486 | 0,981 | 0,94 | 7,18E-06 | rps5 |
| ENSDARG00000056119 | 3,07E-08 | 0,449847957 | 0,935 | 0,81 | 0,000528895 | eef1g |
| ENSDARG00000036044 | 1,03E-13 | 0,446642001 | 1 | 0,99 | 1,78E-09 | rps20 |
| ENSDARG00000036629 | 5,37E-10 | 0,445461792 | 0,944 | 0,92 | 9,25E-06 | rps14 |
| ENSDARG00000036298 | 3,03E-08 | 0,442427658 | 0,972 | 0,9 | 0,000521078 | rps13 |
| ENSDARG00000046157 | 6,82E-09 | 0,434703684 | 0,972 | 0,9 | 0,000117279 | RPS17 |
| ENSDARG00000099104 | 5,36E-07 | 0,429890436 | 0,944 | 0,87 | 0,009220001 | rpl24 |
| ENSDARG00000034897 | 2,14E-07 | 0,427206833 | 0,935 | 0,81 | 0,00368797 | rps10 |
| ENSDARG00000029500 | 1,69E-11 | 0,423801457 | 1 | 0,98 | 2,91E-07 | rpl34 |
| ENSDARG00000006691 | 7,24E-08 | 0,419756148 | 0,981 | 0,84 | 0,00124614 | rpl12 |
| ENSDARG00000099022 | 4,65E-08 | 0,406668115 | 0,981 | 0,93 | 0,000799181 | faua |
| ENSDARG00000037350 | 1,97E-07 | 0,388389959 | 0,972 | 0,89 | 0,003389673 | rpl9 |
| ENSDARG00000054818 | 3,13E-10 | 0,383972959 | 1 | 0,99 | 5,38E-06 | rpl32 |
| ENSDARG00000032725 | 1,12E-07 | 0,383526684 | 0,963 | 0,92 | 0,001931664 | rps27a |
| ENSDARG00000037071 | 5,29E-07 | 0,382106462 | 0,935 | 0,84 | 0,009103674 | rps26 |
| ENSDARG00000100588 | 2,49E-10 | 0,381174873 | 1 | 0,98 | 4,28E-06 | rpl36 |
| ENSDARG00000009285 | 2,46E-09 | 0,38080647 | 1 | 0,98 | 4,24E-05 | rpl15 |
| ENSDARG00000025581 | 1,29E-07 | 0,37588188 | 0,991 | 0,96 | 0,002222275 | rpl10 |
| ENSDARG00000029150 | 1,16E-07 | 0,373624509 | 0,991 | 0,97 | 0,00199155 | hsp90ab1 |
| ENSDARG00000030602 | 3,68E-07 | 0,373014466 | 0,991 | 0,96 | 0,006337542 | rps19 |
| ENSDARG00000029533 | 7,97E-07 | 0,363644756 | 0,981 | 0,97 | 0,013709529 | rpl18 |
| ENSDARG00000100392 | 1,24E-06 | 0,361383873 | 0,972 | 0,94 | 0,021362247 | rps18 |
| ENSDARG00000019778 | 1,33E-07 | 0,359439574 | 0,991 | 0,99 | 0,002287223 | rps6 |
| ENSDARG00000092807 | 4,50E-13 | 0,351086889 | 1 | 1 | 7,74E-09 | RPL41 |
| ENSDARG00000019181 | 7,26E-07 | 0,347668554 | 1 | 0,95 | 0,012497901 | rpsa |
| ENSDARG00000019230 | 4,89E-09 | 0,347238296 | 1 | 0,98 | 8,42E-05 | rpl7a |
| ENSDARG00000070849 | 1,26E-06 | 0,343375408 | 0,991 | 0,95 | 0,021667378 | rps15 |
| ENSDARG00000020850 | 1,48E-07 | 0,340522896 | 1 | 1 | 0,002551015 | eef1a1l1 |
| ENSDARG00000025073 | 3,92E-07 | 0,334925079 | 1 | 0,98 | 0,006751371 | rpl18a |
| ENSDARG00000035871 | 6,55E-07 | 0,33367322 | 0,963 | 0,94 | 0,011271072 | rpl30 |
| ENSDARG00000077291 | 1,35E-06 | 0,306285736 | 1 | 0,98 | 0,023308149 | rps2 |
| ENSDARG00000021113 | 8,29E-07 | -0,453875042 | 0,813 | 0,92 | 0,014254829 | ptmaa |
| ENSDARG00000029722 | 9,89E-07 | -0,475391497 | 0,748 | 0,87 | 0,017008005 | hmgb2a |
| ENSDARG00000052082 | 4,19E-08 | -0,478860473 | 0,766 | 0,87 | 0,000720732 | gabarapb |
| ENSDARG00000068995 | 1,07E-07 | -0,504935093 | 0,729 | 0,86 | 0,001847712 | h2afx1 |
| ENSDARG00000015757 | 5,59E-07 | -0,535191356 | 0,252 | 0,55 | 0,009615622 | tmem50a |
| ENSDARG00000020708 | 4,42E-11 | -0,553933358 | 0,972 | 0,98 | 7,60E-07 | mdkb |
| ENSDARG00000039980 | 3,20E-07 | -0,556916304 | 0,224 | 0,53 | 0,005496957 | ppt1 |
| ENSDARG00000039914 | 6,13E-08 | -0,567239891 | 0,832 | 0,93 | 0,001055049 | gapdhs |
| ENSDARG00000019507 | 6,86E-07 | -0,573919793 | 0,421 | 0,67 | 0,011804965 | mcm5 |
| ENSDARG00000040041 | 9,49E-07 | -0,573932906 | 0,439 | 0,69 | 0,016318969 | mcm4 |
| ENSDARG00000053262 | 7,55E-10 | -0,594399215 | 0,561 | 0,83 | 1,30E-05 | atp1b4 |
| ENSDARG00000102340 | 7,58E-07 | -0,606953941 | 0,346 | 0,62 | 0,013046701 | ptn |
| ENSDARG00000004169 | 3,25E-08 | -0,609120972 | 0,589 | 0,8 | 0,000559692 | stmn1a |
| ENSDARG00000021806 | 4,78E-07 | -0,625146543 | 0,477 | 0,71 | 0,008218484 | zfp36l2 |
| ENSDARG00000043257 | 7,40E-08 | -0,6404879 | 0,729 | 0,85 | 0,001272645 | ckbb |
| ENSDARG00000040162 | 2,23E-08 | -0,647916479 | 0,888 | 0,9 | 0,000382832 | zgc:165461 |
| ENSDARG00000069065 | 3,49E-08 | -0,670603127 | 0,355 | 0,66 | 0,000600087 | btg3 |
| ENSDARG00000104068 | 1,73E-07 | -0,679221757 | 0,57 | 0,72 | 0,002971441 | gstp1 |
| ENSDARG00000025147 | 3,65E-13 | -0,684360508 | 0,757 | 0,89 | 6,28E-09 | cd63 |
| ENSDARG00000098591 | 3,78E-09 | -0,693151546 | 0,523 | 0,78 | 6,50E-05 | tubb2b |
| ENSDARG00000041623 | 1,95E-10 | -0,707891043 | 0,785 | 0,92 | 3,35E-06 | mt2 |
| ENSDARG00000002304 | 3,67E-07 | -0,718917411 | 0,215 | 0,49 | 0,006318206 | gins2 |
| ENSDARG00000093549 | 5,51E-11 | -0,732603948 | 0,57 | 0,83 | 9,48E-07 | selenop |
| ENSDARG00000052470 | 2,01E-07 | -0,756067097 | 0,112 | 0,39 | 0,003452526 | igfbp2a |
| ENSDARG00000100003 | 6,13E-14 | -0,764571923 | 0,682 | 0,9 | 1,05E-09 | glulb |
| ENSDARG00000100795 | 3,13E-07 | -0,792153596 | 0,271 | 0,58 | 0,005392259 | timp4.3 |
| ENSDARG00000094730 | 3,47E-08 | -0,845474181 | 0,318 | 0,59 | 0,000597216 | acbd7 |
| ENSDARG00000038386 | 1,89E-14 | -0,856893988 | 0,654 | 0,91 | 3,26E-10 | ascl1a |
| ENSDARG00000054155 | 3,48E-12 | -0,858249751 | 0,505 | 0,77 | 5,99E-08 | pcna |
| ENSDARG00000071626 | 1,85E-09 | -0,869174469 | 0,364 | 0,7 | 3,18E-05 | ptgdsb.2 |
| ENSDARG00000027088 | 2,12E-06 | -0,902754541 | 0,28 | 0,57 | 0,036486811 | ptgdsb.1 |
| ENSDARG00000007275 | 1,41E-15 | -0,96086584 | 0,897 | 0,91 | 2,42E-11 | si:ch211-251b21.1 |
| ENSDARG00000015349 | 5,12E-14 | -1,034791351 | 0,327 | 0,7 | 8,81E-10 | mfge8a |
| ENSDARG00000041799 | 8,53E-18 | -1,111368149 | 0,598 | 0,9 | 1,47E-13 | cx43 |

Table S2

g2_DMSOvsLY

| | p_val | avg_logFC | pctDMSO | pctLY | p_val_adj | gene_name |
|---|---|---|---|---|---|---|
| ENSDARG00000056732 | 3,25E-87 | 2,60953662 | 0,782 | 0,06 | 5,58E-83 | her4.1 |
| ENSDARG00000007697 | 5,92E-154 | 1,870938433 | 1 | 0,88 | 1,02E-149 | fabp7a |
| ENSDARG00000006514 | 1,45E-44 | 1,608938782 | 0,636 | 0,11 | 2,50E-40 | her6 |
| ENSDARG00000094426 | 3,80E-48 | 1,490537675 | 0,814 | 0,3 | 6,54E-44 | her4.2 |
| ENSDARG00000009822 | 8,29E-31 | 1,461277379 | 0,577 | 0,17 | 1,43E-26 | her4.4 |
| ENSDARG00000056729 | 1,95E-32 | 1,301494704 | 0,672 | 0,22 | 3,36E-28 | her4.2 |
| ENSDARG00000054562 | 2,75E-28 | 1,257448159 | 0,567 | 0,15 | 4,73E-24 | her15.1 |
| ENSDARG00000102453 | 1,19E-92 | 1,217121954 | 0,99 | 0,79 | 2,04E-88 | slc1a2b |
| ENSDARG00000054560 | 2,20E-27 | 1,10047069 | 0,565 | 0,16 | 3,79E-23 | her15.1 |
| ENSDARG00000056438 | 3,03E-19 | 1,062633784 | 0,254 | 0,02 | 5,22E-15 | her9 |
| ENSDARG00000043446 | 1,53E-40 | 1,038198733 | 0,902 | 0,54 | 2,62E-36 | efhd1 |
| ENSDARG00000031044 | 1,79E-19 | 0,959049895 | 0,513 | 0,16 | 3,08E-15 | lipg |
| ENSDARG00000069675 | 3,51E-14 | 0,922552146 | 0,191 | 0,01 | 6,03E-10 | her8.2 |
| ENSDARG00000099776 | 2,27E-39 | 0,834647447 | 0,998 | 0,97 | 3,90E-35 | glula |
| ENSDARG00000007275 | 1,55E-39 | 0,833145934 | 0,995 | 0,89 | 2,66E-35 | si:ch211-251b21.1 |
| ENSDARG00000070770 | 4,56E-15 | 0,800860447 | 0,306 | 0,06 | 7,84E-11 | her4.3 |
| ENSDARG00000068247 | 1,74E-12 | 0,77298634 | 0,582 | 0,3 | 2,99E-08 | luzp2 |
| ENSDARG00000030367 | 1,12E-12 | 0,743864851 | 0,262 | 0,05 | 1,93E-08 | metrn |
| ENSDARG00000077691 | 3,77E-11 | 0,737104469 | 0,355 | 0,11 | 6,49E-07 | slc13a5a |
| ENSDARG00000058966 | 6,88E-08 | 0,708241318 | 0,169 | 0,03 | 0,001183631 | zgc:112332 |
| ENSDARG00000036820 | 1,48E-10 | 0,702885038 | 0,562 | 0,3 | 2,55E-06 | mgll |
| ENSDARG00000029072 | 1,82E-10 | 0,680035383 | 0,369 | 0,13 | 3,13E-06 | klf6a |
| ENSDARG00000042690 | 3,67E-11 | 0,66186146 | 0,533 | 0,25 | 6,31E-07 | s1pr1 |
| ENSDARG00000099793 | 1,55E-09 | 0,65758388 | 0,296 | 0,08 | 2,66E-05 | cspg5b |
| ENSDARG00000023287 | 1,42E-09 | 0,653733372 | 0,289 | 0,08 | 2,44E-05 | hsd17b3 |
| ENSDARG00000095715 | 1,91E-09 | 0,63387481 | 0,428 | 0,19 | 3,29E-05 | BX465834.1 |
| ENSDARG00000013730 | 1,84E-08 | 0,617282658 | 0,597 | 0,34 | 0,000316613 | slc4a4a |
| ENSDARG00000101584 | 4,76E-18 | 0,585091435 | 0,878 | 0,67 | 8,18E-14 | CU467822.1 |
| ENSDARG00000069940 | 1,22E-10 | 0,580908886 | 0,548 | 0,27 | 2,11E-06 | ppap2d |
| ENSDARG00000055876 | 4,07E-08 | 0,580750669 | 0,328 | 0,12 | 0,000699486 | msmo1 |
| ENSDARG00000016868 | 3,87E-08 | 0,57912602 | 0,276 | 0,08 | 0,000666401 | rhobtb4 |
| ENSDARG00000007195 | 3,56E-08 | 0,578258704 | 0,66 | 0,42 | 0,000612713 | grm2b |
| ENSDARG00000019856 | 1,52E-24 | 0,532461332 | 0,993 | 0,92 | 2,61E-20 | atp1a1b |
| ENSDARG00000015059 | 7,20E-07 | 0,509100595 | 0,296 | 0,13 | 0,012393202 | daam1a |
| ENSDARG00000070538 | 1,42E-12 | 0,508046254 | 0,125 | 0 | 2,44E-08 | hey1 |
| ENSDARG00000100968 | 6,72E-09 | 0,496749835 | 0,13 | 0,01 | 0,000115633 | si:ch211-1a19.3 |
| ENSDARG00000039034 | 4,82E-10 | 0,49160247 | 0,8 | 0,57 | 8,29E-06 | marcksl1a |
| ENSDARG00000039352 | 4,43E-07 | 0,489043002 | 0,147 | 0,02 | 0,007612931 | pald1b |
| ENSDARG00000060072 | 4,52E-07 | 0,483793319 | 0,181 | 0,04 | 0,00777034 | abi3a |
| ENSDARG00000036080 | 1,77E-22 | 0,472977703 | 0,985 | 0,88 | 3,05E-18 | cd81a |
| ENSDARG00000036036 | 2,63E-20 | 0,467431795 | 0,978 | 0,91 | 4,53E-16 | mdka |
| ENSDARG00000040162 | 6,89E-14 | 0,461124347 | 0,961 | 0,88 | 1,18E-09 | zgc:165461 |
| ENSDARG00000043283 | 1,58E-06 | 0,452708238 | 0,357 | 0,16 | 0,027148062 | prrg1 |
| ENSDARG00000055129 | 3,36E-08 | 0,449895384 | 0,645 | 0,41 | 0,000577608 | pim3 |
| ENSDARG00000099236 | 2,52E-06 | 0,442499837 | 0,257 | 0,09 | 0,043407935 | sept-10 |
| ENSDARG00000035578 | 8,24E-08 | 0,442228278 | 0,577 | 0,35 | 0,001416903 | hs3st1l2 |
| ENSDARG00000103979 | 9,58E-07 | 0,410148199 | 0,149 | 0,03 | 0,016476946 | calr3a |
| ENSDARG00000062936 | 1,83E-06 | 0,406151711 | 0,127 | 0,02 | 0,031474521 | ndnf |
| ENSDARG00000019532 | 1,26E-06 | 0,405226843 | 0,46 | 0,23 | 0,021651939 | fads2 |
| ENSDARG00000088247 | 9,84E-07 | 0,404231199 | 0,506 | 0,27 | 0,016936778 | si:ch1073-303k11.2 |
| ENSDARG00000016363 | 2,36E-06 | 0,39579255 | 0,12 | 0,02 | 0,040603041 | her8a |
| ENSDARG00000019098 | 1,88E-06 | 0,309301264 | 0,824 | 0,65 | 0,032273198 | cd82a |
| ENSDARG00000037870 | 2,34E-06 | 0,274197538 | 0,946 | 0,85 | 0,04019201 | actb2 |
| ENSDARG00000027744 | 4,28E-07 | 0,273578854 | 0,851 | 0,67 | 0,007362804 | gadd45ba |
| ENSDARG00000087636 | 3,98E-07 | -0,259291019 | 0,868 | 0,93 | 0,006846055 | hmgn6 |
| ENSDARG00000056725 | 3,51E-08 | -0,285336304 | 0,868 | 0,92 | 0,00060436 | hmgb3a |
| ENSDARG00000070922 | 4,29E-07 | -0,314866162 | 0,697 | 0,75 | 0,007388398 | cnbpb |
| ENSDARG00000100003 | 1,31E-07 | -0,328617784 | 0,868 | 0,89 | 0,002258645 | glulb |
| ENSDARG00000062326 | 6,00E-07 | -0,35084672 | 0,665 | 0,75 | 0,010327612 | hmgn7 |
| ENSDARG00000032317 | 2,17E-06 | -0,376486413 | 0,399 | 0,5 | 0,037345826 | tox |
| ENSDARG00000103672 | 3,80E-14 | -0,382645567 | 0,851 | 0,9 | 6,54E-10 | cirbpa |
| ENSDARG00000045248 | 2,39E-12 | -0,386314609 | 0,826 | 0,92 | 4,11E-08 | h3f3d |
| ENSDARG00000092780 | 3,94E-07 | -0,390403788 | 0,017 | 0,1 | 0,006772475 | si:ch1073-170o4.1 |
| ENSDARG00000003920 | 1,99E-08 | -0,392889063 | 0,494 | 0,6 | 0,000341642 | setb |
| ENSDARG00000012399 | 4,48E-07 | -0,3956156 | 0,042 | 0,14 | 0,007709166 | adssl |
| ENSDARG00000036161 | 7,96E-11 | -0,409719053 | 0,641 | 0,73 | 1,37E-06 | hnrnpa0l |
| ENSDARG00000036162 | 2,37E-08 | -0,422744161 | 0,462 | 0,61 | 0,000407972 | hnrnpa0b |
| ENSDARG00000018961 | 1,58E-06 | -0,428505547 | 0,474 | 0,57 | 0,027217547 | gtpbp4 |
| ENSDARG00000003860 | 2,03E-07 | -0,42907932 | 0,357 | 0,48 | 0,003492495 | cbx3a |
| ENSDARG00000006219 | 6,75E-07 | -0,444293179 | 0,103 | 0,21 | 0,011620772 | gnl3 |
| ENSDARG00000078748 | 2,51E-08 | -0,445224107 | 0,249 | 0,42 | 0,000432473 | si:ch211-137a8.4 |
| ENSDARG00000069065 | 1,12E-07 | -0,446749832 | 0,274 | 0,43 | 0,001927305 | btg3 |
| ENSDARG00000068582 | 4,66E-08 | -0,451435546 | 0,271 | 0,41 | 0,000801877 | rnf44 |
| ENSDARG00000033655 | 9,83E-13 | -0,455540911 | 0,653 | 0,81 | 1,69E-08 | stmn1b |
| ENSDARG00000099865 | 2,82E-12 | -0,458014776 | 0,587 | 0,72 | 4,85E-08 | hnrnpabb |
| ENSDARG00000031495 | 1,02E-08 | -0,462582424 | 0,443 | 0,58 | 0,000174661 | seta |
| ENSDARG00000091831 | 9,43E-08 | -0,467575086 | 0,105 | 0,24 | 0,001622296 | ddx27 |
| ENSDARG00000056720 | 1,31E-06 | -0,470302465 | 0,071 | 0,18 | 0,022495813 | utp3 |
| ENSDARG00000002336 | 5,26E-08 | -0,474167409 | 0,017 | 0,12 | 0,000905694 | dlc |
| ENSDARG00000002710 | 1,91E-10 | -0,478851225 | 0,462 | 0,64 | 3,29E-06 | ncl |
| ENSDARG00000015427 | 6,65E-07 | -0,479717915 | 0,296 | 0,44 | 0,011448157 | hdac1 |
| ENSDARG00000055868 | 1,27E-06 | -0,480105566 | 0,225 | 0,33 | 0,021781993 | rsl1d1 |
| ENSDARG00000036675 | 2,39E-08 | -0,491507249 | 0,181 | 0,33 | 0,000410544 | hnrnpa1b |
| ENSDARG00000098221 | 7,30E-07 | -0,494283493 | 0,122 | 0,25 | 0,01255965 | st8sia1 |
| ENSDARG00000028335 | 2,83E-09 | -0,499785587 | 0,242 | 0,39 | 4,88E-05 | hmga1a |
| ENSDARG00000104227 | 4,27E-10 | -0,513083715 | 0,227 | 0,4 | 7,35E-06 | nop10 |
| ENSDARG00000012820 | 3,06E-09 | -0,515924993 | 0,291 | 0,42 | 5,26E-05 | nop56 |
| ENSDARG00000003281 | 2,22E-07 | -0,538154976 | 0,166 | 0,32 | 0,00381258 | pik3ip1 |
| ENSDARG00000008049 | 1,08E-11 | -0,538476763 | 0,301 | 0,53 | 1,85E-07 | si:dkey-42i9.4 |
| ENSDARG00000043608 | 4,35E-13 | -0,538556454 | 0,418 | 0,61 | 7,48E-09 | eif4ebp1 |
| ENSDARG00000104353 | 9,64E-10 | -0,54617836 | 0,164 | 0,32 | 1,66E-05 | nop58 |
| ENSDARG00000078155 | 6,23E-09 | -0,548649047 | 0,071 | 0,21 | 0,000107216 | inavaa |
| ENSDARG00000053912 | 2,37E-09 | -0,548990264 | 0,122 | 0,27 | 4,08E-05 | fbl |
| ENSDARG00000006837 | 1,66E-08 | -0,554787998 | 0,156 | 0,3 | 0,000285505 | mycn |
| ENSDARG00000101628 | 9,39E-09 | -0,558758058 | 0,027 | 0,14 | 0,000161527 | ascl1b |
| ENSDARG00000035993 | 2,96E-10 | -0,56026643 | 0,142 | 0,32 | 5,08E-06 | sumo3b |
| ENSDARG00000019507 | 7,18E-11 | -0,573696729 | 0,029 | 0,14 | 1,24E-06 | mcm5 |
| ENSDARG00000053990 | 3,27E-17 | -0,581410687 | 0,577 | 0,72 | 5,63E-13 | hmgb2b |
| ENSDARG00000024204 | 2,34E-11 | -0,59273415 | 0,037 | 0,18 | 4,03E-07 | mcm3 |
| ENSDARG00000041607 | 1,27E-07 | -0,594164208 | 0,13 | 0,28 | 0,002177134 | eif4ebp3l |
| ENSDARG00000101707 | 3,65E-10 | -0,594176072 | 0,076 | 0,21 | 6,28E-06 | si:ch211-156b7.4 |
| ENSDARG00000020239 | 3,50E-07 | -0,601495996 | 0,281 | 0,39 | 0,006015468 | lpin1 |
| ENSDARG00000054155 | 1,34E-12 | -0,640074596 | 0,071 | 0,23 | 2,31E-08 | pcna |
| ENSDARG00000076532 | 2,58E-15 | -0,645297361 | 0,408 | 0,64 | 4,44E-11 | si:ch211-222l21.1 |
| ENSDARG00000030161 | 9,03E-13 | -0,655760263 | 0,264 | 0,49 | 1,55E-08 | ppp1r14bb |
| ENSDARG00000040041 | 1,91E-16 | -0,684891551 | 0,034 | 0,19 | 3,29E-12 | mcm4 |
| ENSDARG00000057683 | 1,99E-14 | -0,689055539 | 0,029 | 0,19 | 3,42E-10 | mcm6 |
| ENSDARG00000020219 | 2,08E-13 | -0,690576133 | 0,071 | 0,24 | 3,58E-09 | dld |
| ENSDARG00000057738 | 3,98E-13 | -0,716936666 | 0,064 | 0,23 | 6,85E-09 | hells |
| ENSDARG00000101766 | 7,34E-38 | -0,729480285 | 0,846 | 0,95 | 1,26E-33 | ptmab |
| ENSDARG00000077473 | 4,55E-11 | -0,745780022 | 0,105 | 0,28 | 7,84E-07 | mych |
| ENSDARG00000093022 | 2,61E-16 | -0,749486691 | 0,178 | 0,38 | 4,49E-12 | selenoh |
| ENSDARG00000077811 | 2,13E-14 | -0,794318768 | 0,264 | 0,44 | 3,66E-10 | sox11a |
| ENSDARG00000037879 | 1,30E-24 | -0,797087977 | 0,472 | 0,73 | 2,23E-20 | lfng |
| ENSDARG00000014329 | 3,06E-17 | -0,825461469 | 0,103 | 0,33 | 5,27E-13 | npm1a |
| ENSDARG00000102798 | 1,29E-25 | -0,904402812 | 0,01 | 0,23 | 2,22E-21 | mcm2 |
| ENSDARG00000004169 | 7,19E-21 | -0,914791889 | 0,117 | 0,38 | 1,24E-16 | stmn1a |
| ENSDARG00000010791 | 5,62E-48 | -1,010502904 | 0,567 | 0,9 | 9,67E-44 | dla |
| ENSDARG00000095743 | 1,13E-31 | -1,018157895 | 0,306 | 0,66 | 1,94E-27 | sox11b |
| ENSDARG00000004588 | 3,34E-62 | -1,359584783 | 0,411 | 0,87 | 5,75E-58 | sox4a |
| ENSDARG00000101637 | 4,57E-83 | -1,558124585 | 0,457 | 0,91 | 7,86E-79 | ccnd1 |
| ENSDARG00000016725 | 1,73E-51 | -1,676931721 | 0,095 | 0,57 | 2,97E-47 | gadd45gb.1 |
| ENSDARG00000038386 | 9,18E-74 | -1,753737377 | 0,267 | 0,81 | 1,58E-69 | ascl1a |

Table S2
q3_DMSOvsLY

| | p_val | avg_logFC | pctDMSO | pctLY | p_val_adj | gene_name |
|---|---|---|---|---|---|---|
| ENSDARG00000056732 | 2,73E-90 | 3,010492083 | 0,885 | 0,06 | 4,70E-86 | her4.1 |
| ENSDARG00000007697 | 1,70E-95 | 1,731623985 | 0,997 | 0,84 | 2,93E-91 | fabp7a |
| ENSDARG00000094426 | 2,06E-61 | 1,728115174 | 0,931 | 0,35 | 3,55E-57 | her4.2 |
| ENSDARG00000006514 | 2,66E-34 | 1,718829782 | 0,615 | 0,1 | 4,57E-30 | her6 |
| ENSDARG00000009822 | 4,14E-40 | 1,685377071 | 0,773 | 0,21 | 7,12E-36 | her4.4 |
| ENSDARG00000056729 | 2,79E-38 | 1,398431023 | 0,799 | 0,25 | 4,81E-34 | her4.2 |
| ENSDARG00000102453 | 8,51E-67 | 1,274158734 | 0,994 | 0,75 | 1,46E-62 | slc1a2b |
| ENSDARG00000070770 | 2,23E-19 | 1,168269133 | 0,511 | 0,12 | 3,84E-15 | her4.3 |
| ENSDARG00000056438 | 9,23E-20 | 1,130673026 | 0,374 | 0,04 | 1,59E-15 | her9 |
| ENSDARG00000054562 | 5,42E-19 | 1,008383997 | 0,543 | 0,14 | 9,32E-15 | her15.1 |
| ENSDARG00000031044 | 4,61E-10 | 0,942514798 | 0,365 | 0,13 | 7,94E-06 | lipg |
| ENSDARG00000043446 | 4,99E-27 | 0,912334518 | 0,876 | 0,58 | 8,58E-23 | efhd1 |
| ENSDARG00000068247 | 5,41E-11 | 0,902443783 | 0,583 | 0,31 | 9,30E-07 | luzp2 |
| ENSDARG00000023287 | 1,04E-09 | 0,897563294 | 0,388 | 0,13 | 1,79E-05 | hsd17b3 |
| ENSDARG00000007275 | 1,77E-19 | 0,863199542 | 0,971 | 0,89 | 3,04E-15 | si:ch211-251b21.1 |
| ENSDARG00000099793 | 3,49E-09 | 0,854088974 | 0,33 | 0,08 | 6,00E-05 | cspg5b |
| ENSDARG00000054560 | 2,46E-12 | 0,807928422 | 0,526 | 0,2 | 4,23E-08 | her15.1 |
| ENSDARG00000099776 | 3,80E-19 | 0,753667124 | 0,994 | 0,95 | 6,54E-15 | glula |
| ENSDARG00000036820 | 6,73E-08 | 0,735550418 | 0,491 | 0,21 | 0,001158007 | mgll |
| ENSDARG00000007195 | 2,84E-08 | 0,718501026 | 0,569 | 0,31 | 0,00048804 | grm2b |
| ENSDARG00000030367 | 1,63E-09 | 0,711203549 | 0,279 | 0,06 | 2,81E-05 | metrn |
| ENSDARG00000100020 | 4,90E-07 | 0,701220212 | 0,411 | 0,17 | 0,008428993 | pim1 |
| ENSDARG00000042690 | 2,64E-10 | 0,696457042 | 0,661 | 0,35 | 4,55E-06 | s1pr1 |
| ENSDARG00000040942 | 8,93E-08 | 0,687176714 | 0,253 | 0,05 | 0,001535478 | pnp6 |
| ENSDARG00000078659 | 8,82E-09 | 0,628919224 | 0,431 | 0,16 | 0,000151705 | tmem176 |
| ENSDARG00000061817 | 7,24E-11 | 0,628711788 | 0,509 | 0,2 | 1,25E-06 | kif1aa |
| ENSDARG00000019856 | 3,69E-16 | 0,625354333 | 0,977 | 0,88 | 6,36E-12 | atp1a1b |
| ENSDARG00000069940 | 6,69E-09 | 0,614586665 | 0,624 | 0,36 | 0,00011507 | ppap2d |
| ENSDARG00000095715 | 2,28E-07 | 0,610591781 | 0,483 | 0,23 | 0,003923246 | BX465834.1 |
| ENSDARG00000102340 | 5,28E-07 | 0,605505783 | 0,736 | 0,48 | 0,009091147 | ptn |
| ENSDARG00000019702 | 2,68E-07 | 0,582267565 | 0,641 | 0,45 | 0,004604788 | aldocb |
| ENSDARG00000025301 | 1,65E-06 | 0,570208382 | 0,391 | 0,15 | 0,028412203 | gfap |
| ENSDARG00000101584 | 8,66E-11 | 0,554066963 | 0,833 | 0,59 | 1,49E-06 | CU467822.1 |
| ENSDARG00000056078 | 3,68E-07 | 0,552412865 | 0,184 | 0,02 | 0,006337267 | rftn2 |
| ENSDARG00000052139 | 3,68E-07 | 0,495754734 | 0,632 | 0,38 | 0,006322877 | notch3 |
| ENSDARG00000036036 | 3,90E-11 | 0,409598053 | 0,971 | 0,94 | 6,71E-07 | mdka |
| ENSDARG00000040162 | 3,81E-08 | 0,396111943 | 0,977 | 0,94 | 0,000654885 | zgc:165461 |
| ENSDARG00000028367 | 1,77E-06 | 0,341959038 | 0,155 | 0,03 | 0,03047505 | sult2st3 |
| ENSDARG00000103672 | 1,09E-06 | -0,323255851 | 0,839 | 0,9 | 0,01880523 | cirbpa |
| ENSDARG00000062326 | 2,40E-06 | -0,372372018 | 0,578 | 0,66 | 0,041259076 | hmgn7 |
| ENSDARG00000043608 | 1,04E-06 | -0,38679994 | 0,428 | 0,51 | 0,017828543 | eif4ebp1 |
| ENSDARG00000057683 | 9,89E-07 | -0,461240374 | 0,034 | 0,14 | 0,017019033 | mcm6 |
| ENSDARG00000045248 | 1,77E-12 | -0,465229638 | 0,799 | 0,87 | 3,04E-08 | h3f3d |
| ENSDARG00000024204 | 1,36E-07 | -0,478891766 | 0,017 | 0,12 | 0,002333731 | mcm3 |
| ENSDARG00000037879 | 6,45E-07 | -0,501136646 | 0,552 | 0,68 | 0,011101141 | lfng |
| ENSDARG00000031495 | 6,56E-08 | -0,504593324 | 0,379 | 0,55 | 0,001128494 | seta |
| ENSDARG00000020693 | 2,13E-06 | -0,509890795 | 0,589 | 0,67 | 0,036702969 | sesn1 |
| ENSDARG00000003281 | 2,50E-06 | -0,510724816 | 0,193 | 0,31 | 0,042953507 | pik3ip1 |
| ENSDARG00000076532 | 1,12E-06 | -0,512120368 | 0,371 | 0,53 | 0,019242809 | si:ch211-222l21.1 |
| ENSDARG00000037555 | 9,36E-07 | -0,520742288 | 0,032 | 0,14 | 0,016103242 | atoh8 |
| ENSDARG00000053990 | 3,09E-11 | -0,53809288 | 0,557 | 0,7 | 5,32E-07 | hmgb2b |
| ENSDARG00000002336 | 3,98E-09 | -0,555081002 | 0,009 | 0,13 | 6,85E-05 | dlc |
| ENSDARG00000102317 | 3,12E-09 | -0,555571171 | 0,437 | 0,48 | 5,36E-05 | rpl26 |
| ENSDARG00000028119 | 9,47E-10 | -0,559469152 | 0,356 | 0,55 | 1,63E-05 | sumo3a |
| ENSDARG00000033655 | 1,26E-09 | -0,570881309 | 0,578 | 0,73 | 2,17E-05 | stmn1b |
| ENSDARG00000079078 | 2,89E-06 | -0,576573262 | 0,172 | 0,28 | 0,049783879 | si:ch211-5k11.8 |
| ENSDARG00000058369 | 2,39E-07 | -0,578130453 | 0,032 | 0,15 | 0,004106335 | mex3b |
| ENSDARG00000102798 | 2,57E-09 | -0,611499622 | 0,017 | 0,14 | 4,42E-05 | mcm2 |
| ENSDARG00000006640 | 7,59E-07 | -0,616436134 | 0,129 | 0,28 | 0,013057433 | eomesa |
| ENSDARG00000101628 | 3,82E-08 | -0,639590235 | 0,052 | 0,21 | 0,000657101 | ascl1b |
| ENSDARG00000101766 | 4,58E-17 | -0,639775378 | 0,759 | 0,92 | 7,89E-13 | ptmab |
| ENSDARG00000030161 | 1,17E-06 | -0,64619577 | 0,247 | 0,41 | 0,020075334 | ppp1r14bb |
| ENSDARG00000098221 | 7,21E-09 | -0,671522024 | 0,069 | 0,24 | 0,000124039 | st8sia1 |
| ENSDARG00000057738 | 1,01E-11 | -0,681164988 | 0,046 | 0,19 | 1,74E-07 | hells |
| ENSDARG00000078155 | 5,17E-11 | -0,716210907 | 0,023 | 0,19 | 8,89E-07 | inavaa |
| ENSDARG00000077473 | 3,62E-09 | -0,7339659 | 0,098 | 0,28 | 6,24E-05 | mych |
| ENSDARG00000020219 | 1,79E-10 | -0,79192243 | 0,029 | 0,2 | 3,09E-06 | dld |
| ENSDARG00000004169 | 2,78E-13 | -0,800565604 | 0,115 | 0,34 | 4,78E-09 | stmn1a |
| ENSDARG00000077811 | 3,63E-10 | -0,825403848 | 0,115 | 0,32 | 6,25E-06 | sox11a |
| ENSDARG00000041607 | 2,43E-09 | -0,827608435 | 0,147 | 0,34 | 4,17E-05 | eif4ebp3l |
| ENSDARG00000006837 | 4,78E-18 | -0,976120599 | 0,098 | 0,39 | 8,22E-14 | mycn |
| ENSDARG00000095743 | 5,72E-23 | -1,123334096 | 0,221 | 0,59 | 9,83E-19 | sox11b |
| ENSDARG00000010791 | 2,40E-33 | -1,139698355 | 0,414 | 0,84 | 4,13E-29 | dla |
| ENSDARG00000004588 | 4,54E-28 | -1,238316712 | 0,328 | 0,73 | 7,82E-24 | sox4a |
| ENSDARG00000101637 | 2,43E-55 | -1,510110603 | 0,353 | 0,88 | 4,17E-51 | ccnd1 |
| ENSDARG00000016725 | 3,38E-41 | -1,749749097 | 0,052 | 0,55 | 5,82E-37 | gadd45gb.1 |
| ENSDARG00000038386 | 1,61E-56 | -1,795603179 | 0,207 | 0,81 | 2,78E-52 | ascl1a |

[illegible]

[illegible]

| | | | | | | |
|---|---|---|---|---|---|---|
| [illegible] | [illegible] | [illegible] | [illegible] | [illegible] | [illegible] | [illegible] |

Table S2
q4b_DMSOvsLY

| | p_val | avg_logFC | pctDMSO | pctLY | p_val_adj | gene_name |
|---|---|---|---|---|---|---|
| ENSDARG00000056732 | 2,54E-69 | 2,883181685 | 0,98 | 0,28 | 4,37E-65 | her4.1 |
| ENSDARG00000009822 | 6,78E-20 | 1,475202043 | 0,791 | 0,356 | 1,17E-15 | her4.4 |
| ENSDARG00000054562 | 3,98E-24 | 1,366021273 | 0,709 | 0,189 | 6,85E-20 | her15.1 |
| ENSDARG00000094426 | 5,26E-22 | 1,1333402 | 0,929 | 0,606 | 9,05E-18 | her4.2 |
| ENSDARG00000056729 | 1,30E-18 | 0,989329219 | 0,883 | 0,47 | 2,24E-14 | her4.2 |
| ENSDARG00000054560 | 7,26E-13 | 0,960749637 | 0,607 | 0,227 | 1,25E-08 | her15.1 |
| ENSDARG00000070538 | 1,83E-13 | 0,923666889 | 0,383 | 0,045 | 3,15E-09 | hey1 |
| ENSDARG00000030367 | 3,43E-13 | 0,812017293 | 0,5 | 0,129 | 5,90E-09 | metrn |
| ENSDARG00000100020 | 3,75E-08 | 0,737361598 | 0,684 | 0,424 | 0,000645448 | pim1 |
| ENSDARG00000056438 | 9,90E-08 | 0,73247785 | 0,388 | 0,114 | 0,001702485 | her9 |
| ENSDARG00000007697 | 5,67E-43 | 0,71719869 | 1 | 1 | 9,76E-39 | fabp7a |
| ENSDARG00000087260 | 8,44E-09 | 0,698989291 | 0,393 | 0,106 | 0,00014524 | mtss1lb |
| ENSDARG00000031044 | 1,41E-09 | 0,689945792 | 0,653 | 0,318 | 2,43E-05 | lipg |
| ENSDARG00000016363 | 4,54E-10 | 0,689607786 | 0,372 | 0,083 | 7,81E-06 | her8a |
| ENSDARG00000070770 | 3,88E-10 | 0,667894115 | 0,597 | 0,235 | 6,67E-06 | her4.3 |
| ENSDARG00000039034 | 6,02E-17 | 0,663155797 | 0,949 | 0,848 | 1,04E-12 | marcksl1a |
| ENSDARG00000023713 | 1,22E-07 | 0,658029909 | 0,48 | 0,197 | 0,002091524 | aqp1a.1 |
| ENSDARG00000035957 | 7,88E-08 | 0,647054956 | 0,255 | 0,038 | 0,001355966 | gmnn |
| ENSDARG00000060072 | 2,84E-08 | 0,590671503 | 0,551 | 0,258 | 0,000488624 | abi3a |
| ENSDARG00000055759 | 1,25E-07 | 0,541782509 | 0,281 | 0,053 | 0,002151521 | efhd2 |
| ENSDARG00000006514 | 1,21E-06 | 0,512984027 | 0,837 | 0,598 | 0,020800908 | her6 |
| ENSDARG00000061817 | 7,16E-07 | 0,510651297 | 0,76 | 0,492 | 0,012321283 | kif1aa |
| ENSDARG00000042690 | 1,35E-10 | 0,508670155 | 0,974 | 0,871 | 2,32E-06 | s1pr1 |
| ENSDARG00000007195 | 3,85E-10 | 0,492909348 | 0,944 | 0,879 | 6,62E-06 | grm2b |
| ENSDARG00000013168 | 1,94E-06 | 0,480389809 | 0,255 | 0,053 | 0,033403482 | jag1b |
| ENSDARG00000068247 | 7,16E-12 | 0,425226367 | 0,995 | 0,97 | 1,23E-07 | luzp2 |
| ENSDARG00000101584 | 9,03E-07 | 0,391623564 | 0,939 | 0,773 | 0,015542747 | CU467822.1 |
| ENSDARG00000019702 | 9,40E-12 | 0,378379196 | 1 | 1 | 1,62E-07 | aldocb |
| ENSDARG00000043446 | 1,28E-07 | 0,31182981 | 1 | 0,992 | 0,002208841 | efhd1 |
| ENSDARG00000099776 | 3,99E-14 | 0,305792825 | 1 | 1 | 6,87E-10 | glula |
| ENSDARG00000019856 | 2,54E-08 | 0,294045628 | 1 | 1 | 0,000437717 | atp1a1b |
| ENSDARG00000041799 | 6,56E-13 | -0,326025114 | 1 | 1 | 1,13E-08 | cx43 |
| ENSDARG00000074057 | 5,10E-07 | -0,384155981 | 0,663 | 0,75 | 0,008779696 | calm3b |
| ENSDARG00000035870 | 5,92E-07 | -0,408151122 | 0,801 | 0,917 | 0,010186967 | laptm4b |
| ENSDARG00000101766 | 8,57E-07 | -0,467045684 | 0,658 | 0,841 | 0,014751906 | ptmab |
| ENSDARG00000101628 | 1,30E-10 | -0,559968929 | 0 | 0,167 | 2,24E-06 | ascl1b |
| ENSDARG00000004169 | 2,45E-06 | -0,643496412 | 0,066 | 0,258 | 0,04207559 | stmn1a |
| ENSDARG00000095743 | 1,05E-07 | -0,699352727 | 0,163 | 0,417 | 0,001808696 | sox11b |
| ENSDARG00000102317 | 1,35E-09 | -0,701121191 | 0,27 | 0,53 | 2,32E-05 | rpl26 |
| ENSDARG00000004588 | 7,08E-11 | -0,849464898 | 0,189 | 0,523 | 1,22E-06 | sox4a |
| ENSDARG00000020239 | 1,17E-09 | -0,920230924 | 0,383 | 0,606 | 2,01E-05 | lpin1 |
| ENSDARG00000010791 | 2,19E-09 | -0,939736559 | 0,168 | 0,447 | 3,76E-05 | dla |
| ENSDARG00000101637 | 1,00E-16 | -1,15166559 | 0,133 | 0,553 | 1,72E-12 | ccnd1 |
| ENSDARG00000038386 | 3,73E-31 | -1,689665744 | 0,071 | 0,644 | 6,42E-27 | ascl1a |

Table S2

q5_DMSOvsLY

| | p_val | avg_logFC | pctDMSO | pctLY | p_val_adj | gene_name |
|---|---|---|---|---|---|---|
| ENSDARG00000056732 | 7,68E-27 | 1,360674022 | 0,923 | 0,66 | 1,32E-22 | her4.1 |
| ENSDARG00000054562 | 5,76E-09 | 0,86667192 | 0,495 | 0,233 | 9,90E-05 | her15.1 |
| ENSDARG00000022303 | 6,04E-10 | 0,805305203 | 0,814 | 0,62 | 1,04E-05 | higd1a |
| ENSDARG00000000551 | 3,62E-07 | 0,719523914 | 0,649 | 0,453 | 0,006228597 | slc1a4 |
| ENSDARG00000027744 | 4,57E-07 | 0,697076375 | 0,753 | 0,507 | 0,007857463 | gadd45ba |
| ENSDARG00000035859 | 1,27E-06 | 0,694137898 | 0,577 | 0,307 | 0,021782311 | angptl4 |
| ENSDARG00000013250 | 1,60E-06 | 0,675698281 | 0,407 | 0,16 | 0,027510755 | tars |
| ENSDARG00000030367 | 8,53E-08 | 0,670255265 | 0,361 | 0,113 | 0,001468301 | metrn |
| ENSDARG00000070426 | 6,78E-07 | 0,655389614 | 0,696 | 0,513 | 0,011663788 | chac1 |
| ENSDARG00000100020 | 5,64E-07 | 0,626028305 | 0,629 | 0,387 | 0,009696124 | pim1 |
| ENSDARG00000069142 | 1,12E-06 | 0,493579284 | 0,304 | 0,08 | 0,019351706 | aars |
| ENSDARG00000035870 | 6,65E-07 | -0,418894849 | 0,747 | 0,847 | 0,011439814 | laptm4b |
| ENSDARG00000020693 | 1,52E-06 | -0,603876312 | 0,515 | 0,687 | 0,026233957 | sesn1 |
| ENSDARG00000079078 | 2,05E-07 | -0,732763076 | 0,201 | 0,353 | 0,003529522 | si:ch211-5k11.8 |
| ENSDARG00000088711 | 3,52E-07 | -0,797525054 | 0,309 | 0,487 | 0,006056896 | lgals1l1 |
| ENSDARG00000102317 | 3,38E-10 | -0,887101638 | 0,206 | 0,447 | 5,82E-06 | rpl26 |

Table S2
q6_DMSOvsLY

| | p_val | avg_logFC | pctDMSO | pctLY | p_val_adj | gene_name |
|---|---|---|---|---|---|---|
| ENSDARG00000056732 | 2,61E-36 | 2,531168225 | 0,713 | 0,044 | 4,49E-32 | her4.1 |
| ENSDARG00000094426 | 9,51E-16 | 1,721671837 | 0,603 | 0,171 | 1,64E-11 | her4.2 |
| ENSDARG00000009822 | 5,94E-11 | 1,544414009 | 0,397 | 0,07 | 1,02E-06 | her4.4 |
| ENSDARG00000007697 | 4,49E-41 | 1,319145864 | 1 | 0,949 | 7,72E-37 | fabp7a |
| ENSDARG00000056729 | 1,64E-12 | 1,133593623 | 0,544 | 0,133 | 2,83E-08 | her4.2 |
| ENSDARG00000054562 | 1,20E-09 | 1,094480179 | 0,368 | 0,063 | 2,06E-05 | her15.1 |
| ENSDARG00000056438 | 5,50E-09 | 0,978634626 | 0,272 | 0,025 | 9,47E-05 | her9 |
| ENSDARG00000054560 | 3,93E-07 | 0,875415451 | 0,265 | 0,038 | 0,006769296 | her15.1 |
| ENSDARG00000043446 | 5,35E-12 | 0,845444086 | 0,846 | 0,601 | 9,21E-08 | efhd1 |
| ENSDARG00000006514 | 9,65E-12 | 0,843056737 | 0,801 | 0,399 | 1,66E-07 | her6 |
| ENSDARG00000095715 | 4,45E-09 | 0,812059244 | 0,618 | 0,259 | 7,66E-05 | BX465834.1 |
| ENSDARG00000036820 | 3,95E-08 | 0,752510248 | 0,588 | 0,228 | 0,000679269 | mgll |
| ENSDARG00000102453 | 5,13E-15 | 0,747007514 | 0,985 | 0,88 | 8,83E-11 | slc1a2b |
| ENSDARG00000030367 | 1,23E-06 | 0,713539281 | 0,235 | 0,032 | 0,02117681 | metrn |
| ENSDARG00000035957 | 4,28E-07 | 0,678718944 | 0,176 | 0,006 | 0,007370451 | gmnn |
| ENSDARG00000101584 | 1,86E-07 | 0,63794146 | 0,816 | 0,62 | 0,003206015 | CU467822.1 |
| ENSDARG00000045248 | 8,90E-07 | -0,518146053 | 0,699 | 0,778 | 0,015315324 | h3f3d |
| ENSDARG00000076532 | 1,04E-06 | -0,822495028 | 0,235 | 0,468 | 0,017952774 | si:ch211-222l21.1 |
| ENSDARG00000020219 | 8,25E-07 | -0,850713359 | 0,029 | 0,215 | 0,014190179 | dld |
| ENSDARG00000041607 | 2,44E-10 | -1,000386979 | 0,206 | 0,557 | 4,20E-06 | eif4ebp3l |
| ENSDARG00000004588 | 2,86E-09 | -1,012650515 | 0,316 | 0,627 | 4,92E-05 | sox4a |
| ENSDARG00000101628 | 6,01E-09 | -1,054063846 | 0,037 | 0,285 | 0,000103362 | ascl1b |
| ENSDARG00000020239 | 1,53E-08 | -1,086188729 | 0,272 | 0,551 | 0,00026355 | lpin1 |
| ENSDARG00000010791 | 9,04E-14 | -1,286016491 | 0,162 | 0,563 | 1,56E-09 | dla |
| ENSDARG00000077811 | 2,11E-10 | -1,300185348 | 0,162 | 0,462 | 3,64E-06 | sox11a |
| ENSDARG00000095743 | 1,01E-15 | -1,423254942 | 0,184 | 0,582 | 1,73E-11 | sox11b |
| ENSDARG00000016725 | 1,06E-11 | -1,540688664 | 0,029 | 0,316 | 1,83E-07 | gadd45gb.1 |
| ENSDARG00000038386 | 1,32E-18 | -1,614728857 | 0,169 | 0,595 | 2,27E-14 | ascl1a |
| ENSDARG00000101637 | 1,55E-20 | -1,668124394 | 0,235 | 0,734 | 2,67E-16 | ccnd1 |

Table S2
q7_DMSOvsLY

| | p_val | avg_logFC | pctDMSO | pctLY | p_val_adj | gene_name |
|---|---|---|---|---|---|---|
| ENSDARG00000056732 | 1,92E-35 | 2,559010659 | 0,874 | 0,177 | 3,30E-31 | her4.1 |
| ENSDARG00000094426 | 9,87E-13 | 1,750590018 | 0,66 | 0,208 | 1,70E-08 | her4.2 |
| ENSDARG00000054562 | 1,48E-11 | 1,250401152 | 0,476 | 0,115 | 2,54E-07 | her15.1 |
| ENSDARG00000100020 | 1,05E-11 | 1,20172021 | 0,612 | 0,229 | 1,81E-07 | pim1 |
| ENSDARG00000056729 | 4,15E-10 | 1,05773543 | 0,544 | 0,146 | 7,14E-06 | her4.2 |
| ENSDARG00000035957 | 1,06E-08 | 1,03187945 | 0,301 | 0,031 | 0,000182368 | gmnn |
| ENSDARG00000006514 | 9,76E-08 | 0,751884887 | 0,68 | 0,365 | 0,001679922 | her6 |
| ENSDARG00000095715 | 2,16E-07 | 0,686755323 | 0,65 | 0,438 | 0,003708522 | BX465834.1 |
| ENSDARG00000007697 | 4,56E-14 | 0,637660832 | 1 | 0,969 | 7,85E-10 | fabp7a |
| ENSDARG00000101584 | 2,04E-06 | 0,586921496 | 0,825 | 0,698 | 0,03517507 | CU467822.1 |
| ENSDARG00000039034 | 2,13E-08 | 0,559936933 | 0,845 | 0,792 | 0,000366244 | marcksl1a |
| ENSDARG00000042690 | 2,02E-06 | 0,529936396 | 0,68 | 0,635 | 0,034743569 | s1pr1 |
| ENSDARG00000018637 | 1,93E-07 | 0,469613198 | 0,883 | 0,865 | 0,003321695 | sec61g |
| ENSDARG00000095743 | 3,28E-07 | -0,894577865 | 0,214 | 0,521 | 0,005643691 | sox11b |
| ENSDARG00000077811 | 2,40E-06 | -0,939275442 | 0,243 | 0,552 | 0,0413493 | sox11a |
| ENSDARG00000020239 | 6,04E-08 | -1,115420911 | 0,417 | 0,74 | 0,001039873 | lpin1 |
| ENSDARG00000010791 | 2,20E-08 | -1,127065076 | 0,146 | 0,51 | 0,000379137 | dla |
| ENSDARG00000004588 | 5,60E-07 | -1,195096032 | 0,282 | 0,573 | 0,009639947 | sox4a |
| ENSDARG00000038386 | 3,33E-10 | -1,198941273 | 0,107 | 0,531 | 5,72E-06 | ascl1a |
| ENSDARG00000078847 | 3,55E-08 | -1,367421959 | 0 | 0,219 | 0,000610431 | si:dkey-238o13.4 |

Table S2

q8_DMSOvsLY

| | p_val | avg_logFC | pctDMSO | pctLY | p_val_adj | gene_name |
|---|---|---|---|---|---|---|
| ENSDARG00000056732 | 2,56E-23 | 3,650761937 | 0,838 | 0,032 | 4,40E-19 | her4.1 |
| ENSDARG00000094426 | 2,73E-09 | 1,72386567 | 0,706 | 0,194 | 4,70E-05 | her4.2 |
| ENSDARG00000007697 | 1,69E-15 | 1,024979935 | 1 | 0,968 | 2,92E-11 | fabp7a |
| ENSDARG00000095743 | 5,53E-07 | -1,380780103 | 0,103 | 0,516 | 0,00951508 | sox11b |
| ENSDARG00000038386 | 7,99E-12 | -2,10477436 | 0,059 | 0,613 | 1,37E-07 | ascl1a |
| ENSDARG00000101637 | 6,26E-08 | -2,119956738 | 0,029 | 0,419 | 0,001077703 | ccnd1 |

Table S3

a1

| | p_val | avg_logFC | pctDMSO | pctLY | p_val_adj | gene_name |
|---|---|---|---|---|---|---|
| ENSDARG00000056732 | 1.64446299119407e-24 | 2.56672400663982 | 0.87 | 0.096 | 2.7266840856989e-20 | her4.1 |
| ENSDARG00000102453 | 4.46512730433824e-22 | 1.4196486623955 | 1 | 0.948 | 7.40362758332323e-18 | slc1a2b |
| ENSDARG00000007697 | 1.63821326247441e-17 | 1.18459699041562 | 1 | 1 | 2.71632141050882e-13 | fabp7a |
| ENSDARG00000101637 | 5.16420961564523e-17 | -1.25360311132974 | 0.63 | 0.991 | 8.56277596370135e-13 | ccnd1 |
| ENSDARG00000099776 | 2.89932096978359e-15 | 1.20761754702497 | 1 | 0.974 | 4.80736409999817e-11 | glula |
| ENSDARG00000102798 | 2.87123152975951e-14 | -1.2291102248989 | 0.261 | 0.878 | 4.76078899949425e-10 | mcm2 |
| ENSDARG00000013730 | 3.36016102460268e-14 | 1.17090173648653 | 0.848 | 0.27 | 5.57148299489371e-10 | slc4a4a |
| ENSDARG00000004169 | 1.59875406143136e-12 | -0.901987623743702 | 0.717 | 0.974 | 2.65089410925933e-08 | stmn1a |
| ENSDARG00000043446 | 6.81890666353711e-12 | 1.00051378678763 | 0.891 | 0.774 | 1.13064291388109e-07 | efhd1 |
| ENSDARG00000057738 | 2.62620692623538e-11 | -1.07109173008056 | 0.5 | 0.896 | 4.35451370439089e-07 | hells |
| ENSDARG00000019856 | 3.3271696986745e-10 | 0.827984224562579 | 0.978 | 0.965 | 5.51678007737219e-06 | atp1a1b |
| ENSDARG00000062326 | 1.69564472652891e-09 | -0.592312214422587 | 0.957 | 0.974 | 2.81154852105759e-05 | hmgn7 |
| ENSDARG00000014817 | 2.42428988245422e-09 | -0.922556860053647 | 0.326 | 0.8 | 4.01971505409735e-05 | ranbp1 |
| ENSDARG00000102317 | 5.98344158623404e-09 | -1.0931431504984 | 0.609 | 0.852 | 9.92114449413466e-05 | rpl26 |
| ENSDARG00000036820 | 8.77955849594338e-09 | 0.951956934091235 | 0.826 | 0.417 | 0.000145573859421237 | mgll |
| ENSDARG00000006837 | 9.61145438338911e-09 | -1.02688956778452 | 0.174 | 0.661 | 0.000159367525130975 | mycn |
| ENSDARG00000040041 | 2.80802522755071e-08 | -0.887396429528581 | 0.283 | 0.774 | 0.000465598662980183 | mcm4 |
| ENSDARG00000009822 | 3.25165311872189e-08 | 1.37380223526939 | 0.717 | 0.27 | 0.000539156603615276 | her4.4 |
| ENSDARG00000101707 | 3.92210449730469e-08 | -0.786823809182054 | 0.609 | 0.783 | 0.00065032414669809 | si:ch211-156b7.4 |
| ENSDARG00000055225 | 4.60846815397243e-08 | 0.757785395143774 | 0.326 | 0.017 | 0.000764130104610169 | dock8 |
| ENSDARG00000019507 | 5.37893060667912e-08 | -0.935647831963882 | 0.261 | 0.739 | 0.000891880483893464 | mcm5 |
| ENSDARG00000038386 | 8.44407954611283e-08 | -0.956170441569981 | 0.522 | 0.861 | 0.00140011282954097 | ascl1a |
| ENSDARG00000028048 | 8.81192988893177e-08 | 0.919357260945294 | 0.5 | 0.104 | 0.00146110609488378 | rdh8a |
| ENSDARG00000101180 | 9.86487860838852e-08 | -0.848726840354275 | 0.326 | 0.765 | 0.0016356955220569 | mcm7 |
| ENSDARG00000006514 | 1.00556105485919e-07 | 1.20548807509411 | 0.587 | 0.235 | 0.00166732078506201 | her6 |
| ENSDARG00000057683 | 1.19351353858726e-07 | -0.872971852580043 | 0.348 | 0.783 | 0.00197896479833154 | mcm6 |
| ENSDARG00000093022 | 1.47100481853418e-07 | -0.792668498192156 | 0.543 | 0.826 | 0.00243907308961152 | selenoh |
| ENSDARG00000005536 | 2.531340425253e-07 | -0.850649563667486 | 0.13 | 0.583 | 0.004197215559112 | ubr7 |
| ENSDARG00000003941 | 2.56787364651189e-07 | -0.893732229840545 | 0.065 | 0.496 | 0.00425779129328136 | rrs1 |
| ENSDARG00000054155 | 3.25635268246884e-07 | -0.689434590279024 | 0.739 | 0.939 | 0.00539935838280158 | pcna |
| ENSDARG00000099865 | 3.45878327951708e-07 | -0.522691410170208 | 0.87 | 0.974 | 0.00573500855576727 | hnrnpabb |
| ENSDARG00000031495 | 4.13917009517169e-07 | -0.547898305858084 | 0.783 | 0.957 | 0.00686315793480418 | seta |
| ENSDARG00000099793 | 5.28908830510359e-07 | 0.960154004310725 | 0.543 | 0.139 | 0.00876983731869226 | cspg5b |
| ENSDARG00000024204 | 5.646719018715e-07 | -0.906126040382222 | 0.348 | 0.748 | 0.00936282480493134 | mcm3 |
| ENSDARG00000019702 | 8.06449759426841e-07 | 0.945164549835832 | 0.783 | 0.487 | 0.0133717434610564 | aldocb |
| ENSDARG00000094426 | 9.0468586715646e-07 | 1.05224100970409 | 0.848 | 0.522 | 0.0150005963633213 | her4.2 |
| ENSDARG00000002304 | 9.66922349860111e-07 | -0.818489478394416 | 0.283 | 0.678 | 0.0160325394830305 | gins2 |
| ENSDARG00000060586 | 1.10595260882869e-06 | -0.797581900592486 | 0.065 | 0.47 | 0.0183378002069885 | ctdspl2b |
| ENSDARG00000077620 | 1.4504188413589e-06 | -0.941028647060825 | 0.196 | 0.617 | 0.024049394808572 | cdca7a |
| ENSDARG00000074572 | 1.8570368751876e-06 | -0.761384841606433 | 0.283 | 0.678 | 0.0307915284274856 | zgc:56304 |
| ENSDARG00000055455 | 2.04148305611113e-06 | 0.591227391252409 | 0.913 | 0.817 | 0.0338498305533787 | gpm6aa |
| ENSDARG00000068247 | 2.62445405652916e-06 | 0.842739854398221 | 0.739 | 0.443 | 0.04351607271131 | luzp2 |
| ENSDARG00000023299 | 2.75528105901957e-06 | -0.793331644124967 | 0.304 | 0.704 | 0.0456853152396035 | snu13b |

Table S3
a2

| | p_val | avg_logFC | pctDMSO | pctLY | p_val_adj | gene_name |
|---|---|---|---|---|---|---|
| ENSDARG00000007697 | 7.48526123253003e-32 | 1.18081327581205 | 1 | 0.958 | 1.2411311649658e-27 | fabp7a |
| ENSDARG00000102453 | 6.09445661645084e-28 | 1.37424217298618 | 0.92 | 0.672 | 1.01052185157371e-23 | slc1a2b |
| ENSDARG00000101637 | 4.75939120205372e-26 | -0.924552143407954 | 0.81 | 0.99 | 7.89154655212527e-22 | ccnd1 |
| ENSDARG00000056732 | 5.61855505293039e-23 | 1.599459122126 | 0.74 | 0.153 | 9.31612613326388e-19 | her4.1 |
| ENSDARG00000038386 | 3.96887553962012e-21 | -1.01935556493133 | 0.63 | 0.941 | 6.58079253224411e-17 | ascl1a |
| ENSDARG00000010791 | 9.98067665757784e-13 | -0.629864664584051 | 0.87 | 0.986 | 1.65489599659298e-08 | dla |
| ENSDARG00000016725 | 1.8890262083137e-11 | -0.817367665658361 | 0.52 | 0.819 | 3.13219435600495e-07 | gadd45gb.1 |
| ENSDARG00000054562 | 2.71101031619789e-10 | 0.923753764699989 | 0.62 | 0.317 | 4.49512620528772e-06 | her15.1 |
| ENSDARG00000099572 | 3.87893188034663e-10 | 0.688062412827492 | 0.99 | 0.913 | 6.43165695080275e-06 | hmgn2 |
| ENSDARG00000103754 | 1.14789356146016e-09 | 0.882496740638911 | 0.27 | 0.017 | 1.90332231425709e-05 | aspm |
| ENSDARG00000091150 | 1.19339272902434e-09 | 1.04424815730291 | 0.43 | 0.087 | 1.97876448399526e-05 | mki67 |
| ENSDARG00000006514 | 1.19524362235137e-09 | 0.855415236157433 | 0.57 | 0.171 | 1.98183345022081e-05 | her6 |
| ENSDARG00000100003 | 1.20120786631565e-09 | -0.661605860107819 | 0.7 | 0.889 | 1.99172276313799e-05 | glulb |
| ENSDARG00000043446 | 1.3758230779518e-09 | 0.693408850451165 | 0.7 | 0.495 | 2.28125224555188e-05 | efhd1 |
| ENSDARG00000102317 | 1.5197370258593e-09 | -0.848779832405124 | 0.69 | 0.77 | 2.5198759625773e-05 | rpl26 |
| ENSDARG00000103919 | 1.95311641557963e-09 | 0.654622719150308 | 1 | 0.941 | 3.23846232867259e-05 | si:ch73-1a9.3 |
| ENSDARG00000057683 | 5.21449388463008e-09 | -0.790633411229049 | 0.32 | 0.634 | 8.64615231010513e-05 | mcm6 |
| ENSDARG00000055133 | 4.22331573821128e-08 | 0.840741753888345 | 0.32 | 0.038 | 0.000700267982552812 | cenpf |
| ENSDARG00000004588 | 7.32385980661801e-08 | -0.546223838852346 | 0.76 | 0.923 | 0.00121436919453533 | sox4a |
| ENSDARG00000024204 | 9.70213337312666e-08 | -0.770540129209973 | 0.33 | 0.571 | 0.00160871073459813 | mcm3 |
| ENSDARG00000094426 | 1.55809811002787e-07 | 0.587681244933905 | 0.88 | 0.638 | 0.00258348247623722 | her4.2 |
| ENSDARG00000007275 | 2.18782495997092e-07 | 0.73845940116163 | 0.97 | 0.861 | 0.00362763256612778 | si:ch211-251b21.1 |
| ENSDARG00000037879 | 4.7438327297659e-07 | -0.551668954341155 | 0.77 | 0.895 | 0.00786574904922484 | lfng |
| ENSDARG00000045167 | 4.77135893820468e-07 | 0.659325038032897 | 0.25 | 0.024 | 0.00791139025543718 | dlgap5 |
| ENSDARG00000041723 | 6.09697794537295e-07 | 0.798844625964949 | 0.37 | 0.111 | 0.0101093991312229 | TUBB4B |
| ENSDARG00000104353 | 7.40477849935743e-07 | -0.545219992441619 | 0.41 | 0.645 | 0.0122778632297845 | nop58 |
| ENSDARG00000002344 | 1.00377326589741e-06 | 0.470479059089211 | 0.98 | 0.948 | 0.016643564521845 | tubb4b |
| ENSDARG00000011094 | 1.64060543523024e-06 | 0.745575891263216 | 0.32 | 0.063 | 0.0272028787215526 | ccna2 |
| ENSDARG00000077811 | 2.2037732268306e-06 | -0.525037689116356 | 0.58 | 0.749 | 0.0365407638740782 | sox11a |

Table S3
a3

| | p_val | avg_logFC | pctDMSO | pctLY | p_val_adj | gene_name |
|---|---|---|---|---|---|---|
| ENSDARG00000007697 | 3.70008348545566e-19 | 1.01107152656666 | 1 | 0.966 | 6.13510842723404e-15 | fabp7a |
| ENSDARG00000038386 | 1.7944729903797e-10 | -0.908605858912313 | 0.632 | 0.903 | 2.97541566534859e-06 | ascl1a |
| ENSDARG00000102453 | 4.10641478619109e-10 | 0.894638409054517 | 0.779 | 0.63 | 6.80884635698345e-06 | slc1a2b |
| ENSDARG00000056732 | 4.01597938817638e-09 | 1.30087660416832 | 0.515 | 0.172 | 6.65889542353526e-05 | her4.1 |

Table S3

a4

| | p_val | avg_logFC | pctDMSO | pctLY | p_val_adj | gene_name |
|---|---|---|---|---|---|---|
| ENSDARG00000102317 | 2.5617664437587e-13 | -1.16535229877411 | 0.592 | 0.779 | 4.24766494039629e-09 | rpl26 |
| ENSDARG00000056732 | 4.14318445347531e-11 | 1.61314912028625 | 0.487 | 0.183 | 6.86981414230741e-07 | her4.1 |
| ENSDARG00000101794 | 1.03150091525372e-06 | 0.483970410821948 | 0.737 | 0.562 | 0.017103316675822 | atp6v0e1 |

Table S3

a5

| | p_val | avg_logFC | pctDMSO | pctLY |
|---|---|---|---|---|
| ENSDARG00000099572 | 4.69909926376722e-09 | 0.391334141600304 | 1 | 1 |
| ENSDARG00000056732 | 6.63420050152163e-08 | 0.71476584477945 | 0.429 | 0.106 |
| ENSDARG00000102317 | 5.49294742441728e-07 | -0.79992446191744 | 0.714 | 0.767 |
| ENSDARG00000059304 | 2.34840589155943e-06 | 0.622451636588065 | 0.494 | 0.183 |

| p_val_adj | gene_name |
|---|---|
| 7.79157648925243e-05 | hmgn2 |
| 0.0011000167851573 | her4.1 |
| 0.00910785612442629 | rpl26 |
| 0.0389389180879469 | chchd2 |

Table S3

a6

| | p_val | avg_logFC | pctDMSO | pctLY |
|---|---|---|---|---|
| ENSDARG00000056732 | 1.67910554529825e-09 | 0.982729097626316 | 0.463 | 0.146 |
| ENSDARG00000007697 | 5.11234448029456e-09 | 0.585378914352631 | 0.979 | 0.943 |
| ENSDARG00000101794 | 9.00007754454406e-09 | 0.576686444431892 | 0.768 | 0.586 |
| ENSDARG00000009822 | 6.62256245792713e-08 | 0.8452251501492 | 0.611 | 0.357 |
| ENSDARG00000043446 | 1.21118362800836e-06 | 0.528288141091376 | 0.358 | 0.178 |
| ENSDARG00000102453 | 2.64521735718982e-06 | 0.75689692024272 | 0.463 | 0.268 |

| p_val_adj | gene_name |
|---|---|
| 2.78412490465903e-05 | her4.1 |
| 8.47677838277641e-05 | fabp7a |
| 0.000149230285766085 | atp6v0e1 |
| 0.0010980870811489 | her4.4 |
| 0.0200826357360066 | efhd1 |
| 0.0438603489995645 | slc1a2b |

Table S3

a7

| | p_val | avg_logFC | pctDMSO | pctLY | p_val_adj | gene_name |
|---|---|---|---|---|---|---|
| ENSDARG00000056732 | 5.72342662785114e-14 | 1.68674459618353 | 0.612 | 0.144 | 9.49001369163997e-10 | her4.1 |
| ENSDARG00000006514 | 6.3368292127775e-11 | 1.013004397035 | 0.418 | 0.031 | 1.05070965177064e-06 | her6 |
| ENSDARG00000102453 | 2.09027176087752e-08 | 1.41741777826453 | 0.597 | 0.258 | 0.000346587960671101 | slc1a2b |
| ENSDARG00000007697 | 8.78282818087312e-08 | 0.760750662380947 | 1 | 0.948 | 0.00145628074067057 | fabp7a |

Table S4

Primary antibodies

| Antibody | Source | Identifier | Dilution |
|---|---|---|---|
| Mouse monoclonal (IgG1) anti- ZO1 | Thermo Fisher | Cat# 33-9100, RRID: AB_2533147 | 1/200 |
| Mouse monoclonal (IgG2a) anti-GS | Millipore | Cat# MAB302, RRID:AB_2110656 | 1/500 |
| Chicken anti-mCherry | EnCor Biotechnology | Cat# CPCA-mCherry, RRID:AB_2572308 | 1/500 |
| Chicken anti-GFP Antibody | Aves lab | Cat# GFP-1020; RRID: AB_10000240 | 1/1000 |
| Mouse anti-PCNA IgG2a (PC10) | Santa Cruz Biotechnology | Cat# sc-56, RRID:AB_628110 | 1/250 |
| Rabbit Polyclonal anti-PCNA | GeneTex | Cat# GTX124496; RRID: AB_11161916 | 1/500 |

Table S4

Secondary antibodies

| Antibody | Source | Identifier | Dilution |
|---|---|---|---|
| Goat anti-Chicken IgG(H+L) Alexa488 Conjugated | Thermo Fisher Scientific | Cat# A-11039, RRID:AB_142924 | 1/1000 |
| Goat anti-Chicken IgG(H+L) Alexa555 Conjugated | Thermo Fisher Scientific | Cat# A-21437, RRID:AB_2535858 | 1/1000 |
| Goat anti-Rabbit IgG Alexa546 | Thermo Fisher Scientific | Cat# A-11010, RRID:AB_2534077 | 1/1000 |
| Goat anti-Mouse IgG Alexa405 conjugated | Thermo Fisher Scientific | Cat# A-31553, RRID:AB_221604 | 1/1000 |
| Goat anti-Mouse IgG1 Alexa647 conjugated | Thermo Fisher Scientific | Cat# A-21240, RRID:AB_2535809 | 1/1000 |
| Goat anti-Mouse IgG2a Alexa555 conjugated | Thermo Fisher Scientific | Cat# A-21137, RRID:AB_2535776 | 1/1000 |
| Goat anti-Mouse IgG2a Alexa633 conjugated | Thermo Fisher Scientific | Cat# A-21136, RRID:AB_2535775 | 1/1000 |

Table S5

| Gene name | Entrez Gene ID | Accession Number | Channel |
|---|---|---|---|
| nr2f1b | 393564 | NM_200592.1 | T2 |
| ascl1a | 30466 | NM_131219.1 | T1 |
| ascl1a | 30466 | NM_131219.1 | T3 |
| timp4.3 | 100124610 | ENSDART00000187110.1 | T2 |
